# CMPK2 restricts *Mycobacterium tuberculosis* replication and regulates macrophage gene expression

**DOI:** 10.64898/2026.05.19.726211

**Authors:** John Neff, Victoria A. Ektnitphong, Priscila C. Campos, Kubra F. Naqvi, Beatriz R. S. Dias, Kathryn C. Rahlwes, Zhenyu Zhong, Michael U. Shiloh

## Abstract

Host cell metabolic pathways influence innate immune responses to intracellular pathogens, but the contribution of nucleotide metabolism to antimicrobial defense remains incompletely defined. Here, we identify the mitochondrial nucleoside monophosphate kinase CMPK2 as a regulator of macrophage responses to *Mycobacterium tuberculosis* (Mtb). Using a targeted genetic screen of candidate host factors, we found that depletion of CMPK2 enhances intracellular Mtb replication in human macrophages. This phenotype was confirmed using both shRNA-mediated knockdown and CRISPR-Cas9-mediated knockout approaches. CMPK2 expression increased following macrophage activation and Mtb infection. Transcriptomic profiling revealed that loss of CMPK2 is associated with broad alterations in gene expression, including reduced expression of genes linked to innate immune and inflammatory responses early after infection. In contrast, myeloid-specific deletion of *Cmpk2* in mice did not significantly alter bacterial burden or survival following aerosol Mtb infection, indicating that the contribution of CMPK2 to host defense is context dependent. Together, these findings identify CMPK2 as a host factor that limits Mtb replication in human macrophages and shapes innate immune gene expression programs.

## Introduction

*Mycobacterium tuberculosis* (Mtb) remains a leading cause of infectious mortality worldwide (1). As an intracellular pathogen, Mtb establishes infection within macrophages, where host cell-intrinsic defenses play a central role in determining bacterial survival and replication (2). Innate immune signaling pathways, including those activated by microbial products and host-derived danger signals, are critical components of this response (2-4).

Among these pathways, cytosolic DNA sensing has emerged as an important mechanism contributing to host responses to Mtb infection. Prior work has demonstrated that Mtb infection can lead to the accumulation of host and bacterial DNA in the cytosol, resulting in activation of downstream signaling pathways that shape inflammatory and antimicrobial responses (3, 5-8). While these pathways have been well described, the upstream cellular processes that influence their activation remain incompletely defined.

We recently used activity-based protein profiling (ABPP) to identify host proteins whose expression or activity changes during Mtb infection (9). Building on these findings, we performed a targeted genetic screen to assess the functional contribution of candidate host factors to intracellular Mtb replication. Through this approach, we identified the mitochondrial nucleoside monophosphate kinase CMPK2 as a regulator of macrophage responses to Mtb.

CMPK2 catalyzes the phosphorylation of pyrimidine nucleotides within mitochondria (10) and has been linked to innate immune responses in other contexts, including antiviral defense (11). However, its role during Mtb infection has not been defined. Here, we investigate the contribution of CMPK2 to macrophage control of Mtb and its impact on host transcriptional responses during infection.

## RESULTS

### CMPK2 knockdown permits increased intracellular *Mycobacterium tuberculosis* replication in human macrophages

To identify host factors that influence intracellular Mtb replication, we generated shRNA-mediated knockdown lines targeting candidate genes identified in a prior activity-based protein profiling screen (9). We infected these cells with a luminescent Mtb strain (12) and quantified bacterial growth over time.

Several knockdowns altered bacterial replication, with *CMPK2* depletion producing one of the most consistent increases in luminescence across independent shRNA constructs (Figure 1A). We confirmed efficient knockdown of CMPK2 at both the transcript and protein levels (Figure 1B-C). To validate these findings using an orthogonal approach, we measured bacterial burden by colony-forming unit (CFU) assays. *CMPK2* knockdown resulted in a reproducible increase in intracellular Mtb replication (Figure 1D).

**Figure 1.**
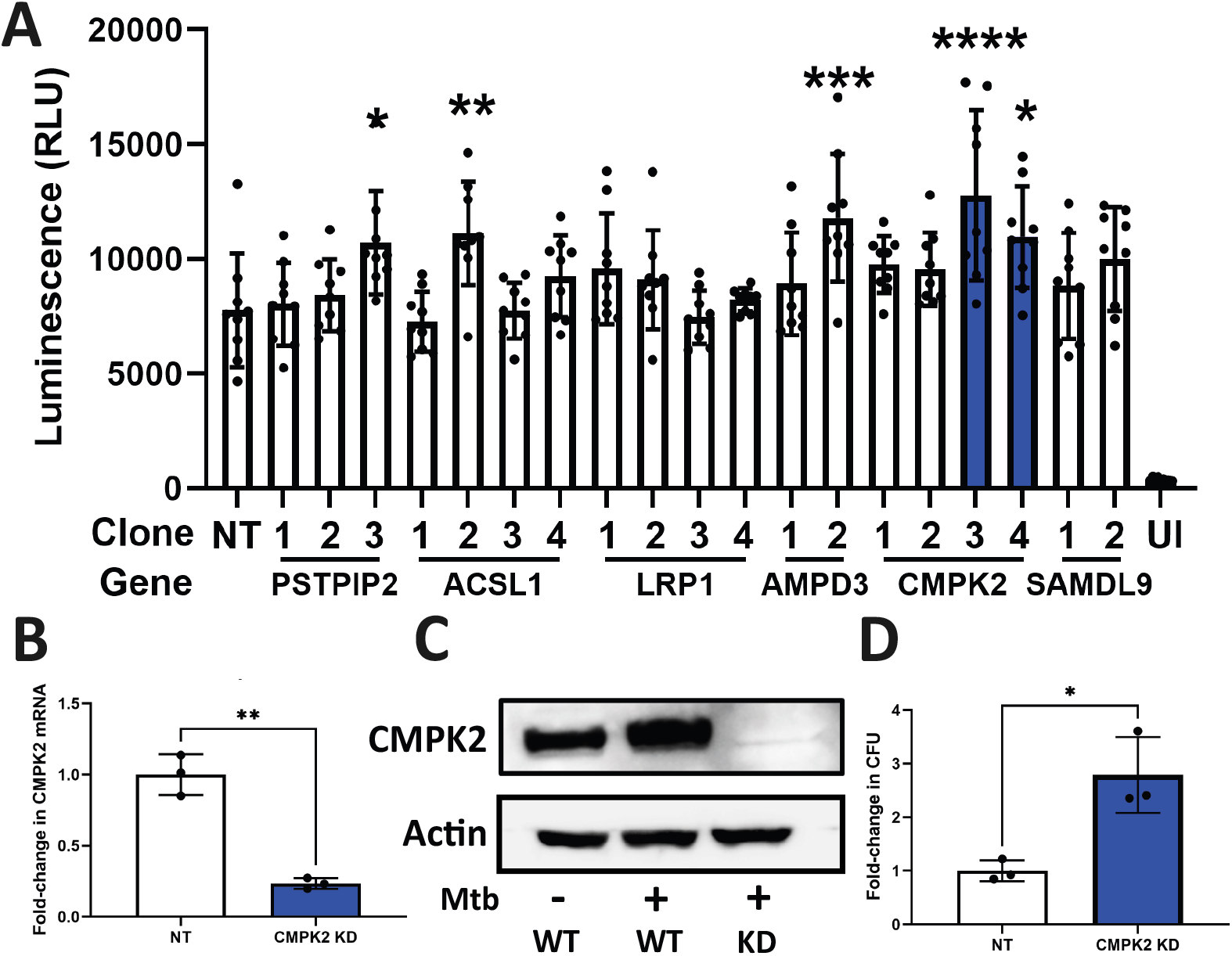
CMPK2 knockdown enhances *M. tuberculosis* replication in human macrophages. (A) Luminescence measurements of THP1 shRNA knockdown lines following infection with Mtb-lux (MOI 3) over 4 days. CMPK2-targeting shRNA lines are highlighted. Statistical comparisons were performed relative to non-targeting (NT) control shRNA. (B) qPCR analysis confirming CMPK2 knockdown in THP1 cells. (C) Immunoblot analysis of CMPK2 expression in wild-type THP1 cells and CMPK2 shRNA knockdown lines. (D) Intracellular bacterial burden measured by CFU assay 4 days following infection (MOI 1) of THP1 wild-type and CMPK2 knockdown cells. Data are shown as mean ± SEM of n = 3 independent biological replicates, each performed in technical triplicate. Statistical significance was determined using a one-way ANOVA with multiple comparisons (A) and an unpaired t-test with Welch’s correction (B, D). *, p < 0.05; **, p < 0.01; ***, p < 0.001; ****, p < 0.0001.

### CMPK2 expression is induced during macrophage activation and Mtb infection

To further confirm the phenotype observed in *CMPK2* knockdown cells, we used a THP1 macrophage cell line with CRISPR-Cas9-mediated deletion of *CMPK2* (Figure 2A) (13). Notably, CMPK2 expression is low under basal conditions but has been reported to increase following innate immune stimulation with lipopolysaccharide (LPS) (14). To confirm this in our system, we treated THP1 macrophages with LPS and observed robust induction of CMPK2 protein expression by immunoblot (Figure 2A). LPS did not induce CMPK2 in THP1 *CMPK2* KO cells, as expected (Figure 2A).

**Figure 2.**
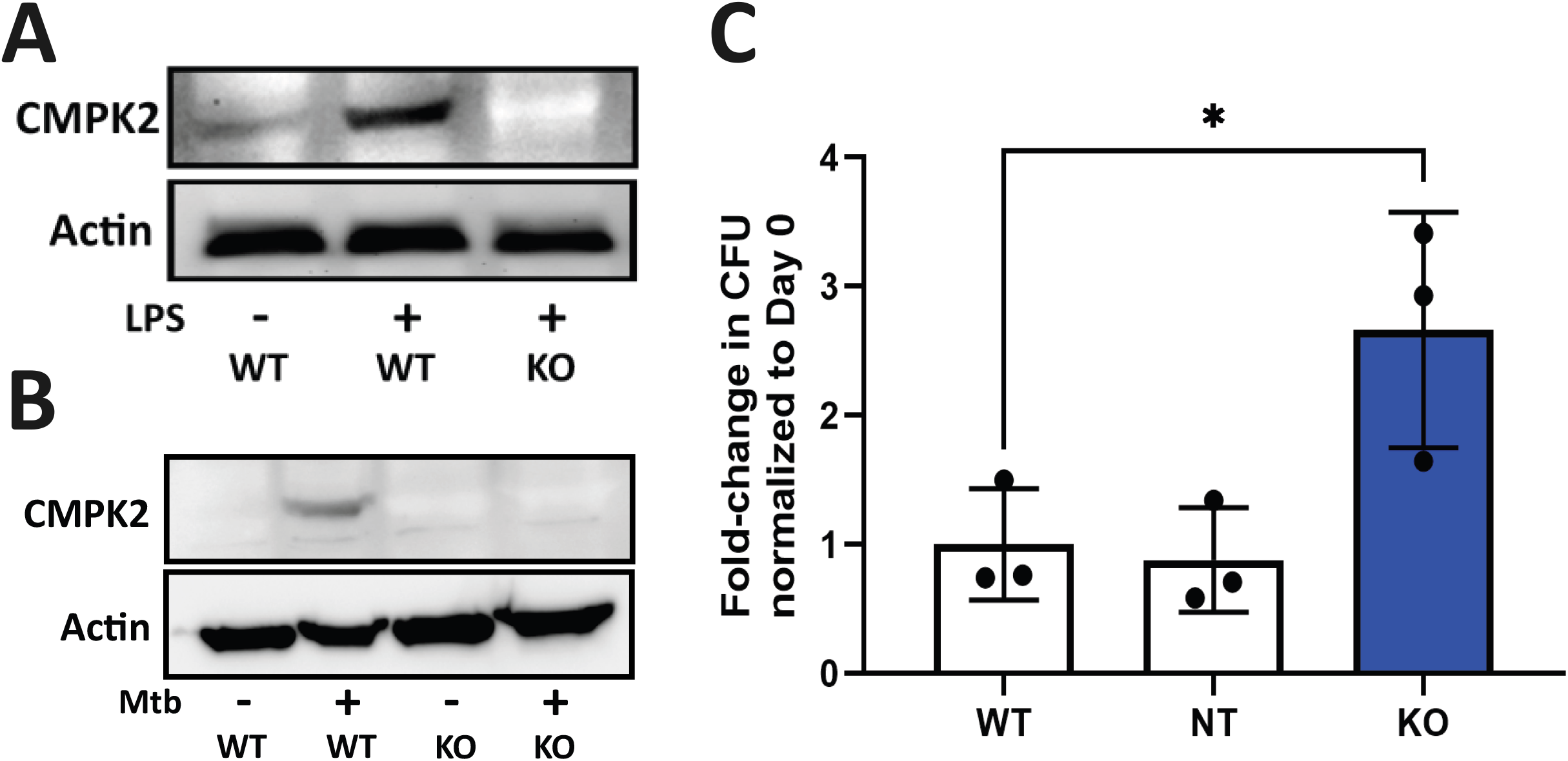
CMPK2 expression is induced by inflammatory stimulation and *M. tuberculosis* infection and restricts intracellular bacterial growth. (A) Immunoblot analysis of CMPK2 expression in wild-type THP1 cells, LPS-treated wild-type THP1 cells, and LPS-treated THP1 *CMPK2* knockout cells. (B) Immunoblot analysis of CMPK2 expression in uninfected and Mtb-infected wild-type and THP1 *CMPK2* knockout cells. (C) Intracellular bacterial burden measured by CFU assay 4 days following infection (MOI 1) of wild-type, non-targeting control, and THP1 *CMPK2* knockout cells. Data are shown as mean ± SEM of n = 3 independent biological replicates, each performed in technical triplicate. Statistical significance was determined using a one-way ANOVA with multiple comparisons. *, p < 0.05.

We next assessed CMPK2 expression during Mtb infection. CMPK2 protein levels increased following infection, with detectable accumulation by 16 hours post-infection (Figure 2B). These findings indicate that CMPK2 is induced in response to both inflammatory stimulation and Mtb infection, consistent with a role in macrophage activation.

### Human THP1 *CMPK2* knockout cells are permissive for Mtb replication

To confirm phenotype of increased growth observed in THP1 *CMPK2* KD cells, we infected THP1 *CMPK2* KO cells with Mtb and quantified Mtb replication. Consistent with the knockdown data, CMPK2-deficient cells exhibited increased Mtb replication compared to control cells (Figure 2C). Together with the data in *CMPK2* KD cells, these results demonstrate that CMPK2 limits intracellular Mtb replication in human macrophages.

### Loss of *CMPK2* alters macrophage transcriptional responses to Mtb infection

To define how CMPK2 influences host responses to infection, we performed bulk RNA sequencing on wild-type and CMPK2-deficient THP1 macrophages under uninfected conditions and at 4 hours post-Mtb infection.

Unsupervised analysis of global gene expression revealed clear separation between wild-type and CMPK2-deficient cells in both uninfected and infected conditions, indicating that loss of CMPK2 is associated with broad transcriptional reprogramming (Figure 3A, C, D). Differential expression analysis identified hundreds of genes that were significantly altered in CMPK2-deficient cells relative to wild-type cells (Figure 3B), with substantial overlap between uninfected and infected conditions.

**Figure 3.**
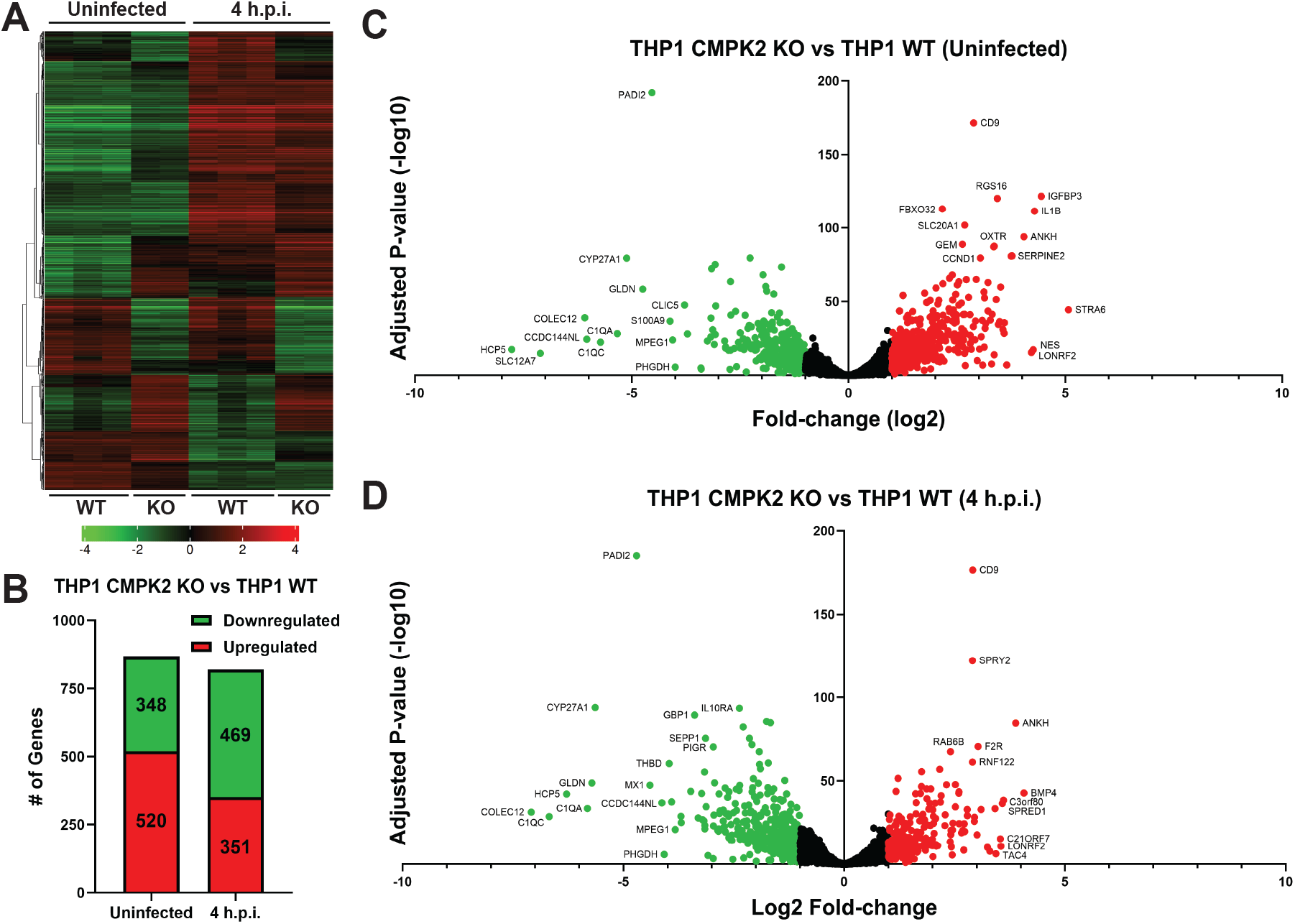
Loss of CMPK2 alters macrophage gene expression in uninfected and infected conditions. (A) Heat map of differentially expressed genes between wild-type and CMPK2 knockout THP1 cells under uninfected and Mtb-infected conditions. Data are centered by subtracting the mean expression level for each gene. Color scale represents gene-level Z-scores, with green indicating downregulation and red indicating upregulation. (B) Number of differentially expressed genes between wild-type and CMPK2 knockout THP1 cells in uninfected and infected conditions. (C, D) Volcano plots showing differential gene expression between wild-type and CMPK2 knockout THP1 cells in uninfected (C) and infected (D) conditions.

Under basal conditions, CMPK2-deficient cells exhibited increased expression of select genes, suggesting that CMPK2 contributes to maintenance of transcriptional homeostasis in resting macrophages. In contrast, following Mtb infection, CMPK2-deficient cells showed a relative reduction in the expression of genes associated with innate immune and inflammatory responses.

Among the genes reduced in CMPK2-deficient cells were multiple transcripts previously implicated in macrophage activation and host defense, including components of complement pathways and interferon-responsive gene programs (Figure 3 C, D; Supplemental Figures 1 and 2). Functional enrichment analysis of differentially expressed genes further supported an association between CMPK2 and pathways linked to innate immunity and responses to infection (Supplemental Tables 1 and 2).

While these changes were not restricted to a single signaling pathway, the overall pattern suggests that CMPK2 contributes to the early transcriptional response of macrophages to Mtb infection. Together, these data indicate that loss of CMPK2 leads to a broad but coordinated alteration in gene expression, consistent with a role in regulating macrophage activation states.

### CMPK2 is dispensable for control of Mtb infection in vivo

To determine whether CMPK2 contributes to host defense in vivo, we generated mice with myeloid-specific deletion of Cmpk2 using a LysMCre driver (15) to delete *Cmpk2* in Cmpk2^fl/fl^ mice (16) (*Cmpk2*^fl/fl LysMCre /+^). We then infected *Cmpk2*^fl/fl LysMCre /+^ mice along with littermate controls with Mtb using an aerosol model and assessed both survival and bacterial burden.

After a high-dose aerosol infection, *Cmpk2*-deficient mice showed a trend toward reduced survival compared to controls, although this did not reach statistical significance (Figure 4A-B). After low-dose aerosol infection, survival was similar between groups (Figure 4C-D).

**Figure 4.**
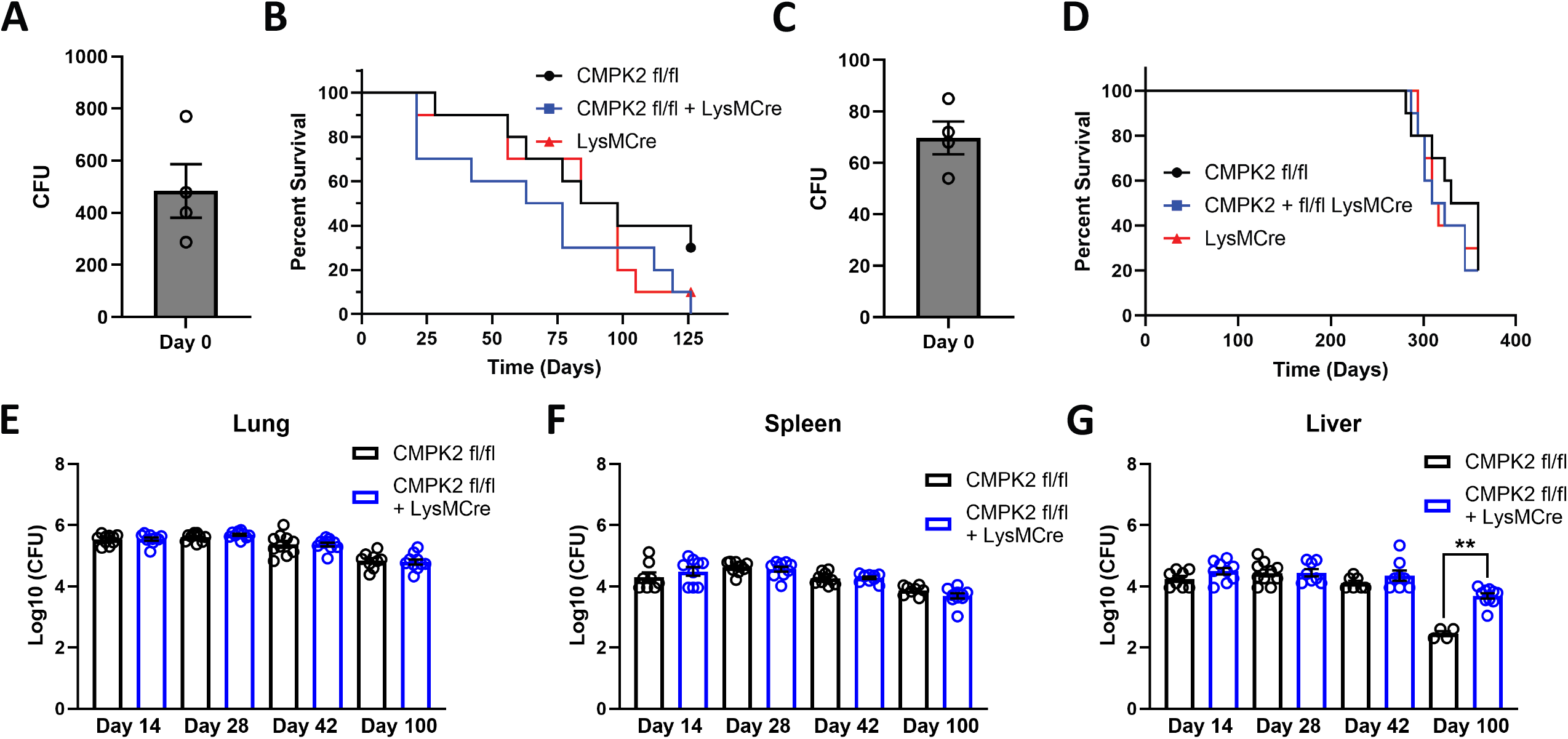
CMPK2 is dispensable for control of *M. tuberculosis* infection in vivo. (A, C) Day-zero bacterial lung inoculum following high-dose (A) and low-dose (C) aerosol infection. (B, D) Survival of WT, *Cmpk2*^fl/fl^, and *Cmpk2*^fl/fl LysMCre /+^ mice following high-dose (B) and low-dose (D) aerosol infection. (E, F, G) Bacterial burden in the lungs (E), spleen (F), and liver (G) of *Cmpk2*^fl/fl^ and *Cmpk2*^fl/fl LysMCre /+^ mice at the indicated time points following low-dose infection. Each symbol represents an individual mouse (n=10 mice/genotype at each time point). Bars indicate mean ± SEM. Statistical significance was determined using Kaplan Meier analysis (B, D) and Mann-Whitney tests of individual timepoints (E, F, G).

Measurement of bacterial burden in the lungs, spleen, and liver over time revealed no significant differences between genotypes (Figure 4E-G).

These findings indicate that, despite its role in human macrophages in vitro, CMPK2 is dispensable for control of aerosol Mtb infection in mice.

## DISCUSSION

In this study, we identify CMPK2 as a host factor that limits Mtb replication in human macrophages and shapes transcriptional responses during infection. Using both knockdown and knockout approaches, we demonstrate that loss of CMPK2 results in increased intracellular bacterial growth in human macrophages, supporting a role for CMPK2 in cell-intrinsic antimicrobial defense in human cells.

Our transcriptional analysis provides additional insight into the consequences of CMPK2 loss. CMPK2-deficient macrophages exhibited widespread changes in gene expression under both resting and infected conditions, indicating that CMPK2 contributes to basal cellular homeostasis as well as infection-induced responses. Notably, following Mtb infection, CMPK2-deficient cells showed reduced expression of genes associated with innate immune and inflammatory pathways. Several of the most strongly downregulated transcripts in CMPK2-deficient cells provide additional context for these changes. For example, expression of *PADI2* was reduced, a gene that has been linked to macrophage inflammatory programs and the abundance of NLRP3+ macrophages (17). Given that NLRP3 inflammasome activation can depend on mitochondrial DNA and is associated with proinflammatory signaling (14), reduced *PADI2* expression is consistent with a diminished inflammatory state in CMPK2-deficient cells. In addition, multiple genes with established roles in host defense were among the top downregulated transcripts, including *MPEG1, MX1*, and *GBP1*, each of which has been implicated in antimicrobial or interferon-driven responses (18-20). Together, these findings support the conclusion that loss of CMPK2 is associated with a broad attenuation of innate immune gene expression programs during Mtb infection.

We also noted reduced expression of *CYP27A1*, a key enzyme involved in the generation of oxysterols that regulate macrophage function and innate immune responses (21). Oxysterols have been implicated in antimicrobial activity and immune cell signaling (22), raising the possibility that altered sterol metabolism contributes to the phenotype observed in CMPK2-deficient macrophages. Taken together, these findings are consistent with a model in which CMPK2 supports early macrophages transcriptional responses to infection, rather than acting within a single defined signaling pathway. The coordinated downregulation of genes involved in antimicrobial defense, inflammatory signaling, and metabolic regulation suggests that CMPK2 contributes to macrophage activation more broadly. However, the specific mechanisms by which CMPK2 regulates gene expression during Mtb infection remain to be defined.

An alternative, non-transcriptional explanation for the cell-intrinsic activity of CMPK2 may relate to its non-catalytic N-terminal domain, which has been shown to mediate antiviral activity independent of its enzymatic function (11). This raises the possibility that CMPK2 may exert direct antimicrobial effects in macrophages, although this was not tested in the present study.

Despite the clear phenotype in human macrophages, we did not detect a significant effect of myeloid-specific *Cmpk2* deletion on bacterial burden or survival in mice. Several explanations may account for this discrepancy. First, the magnitude of the cell-intrinsic phenotype observed in vitro may be insufficient to influence bacterial control at the organismal level. Second, redundant or compensatory pathways may limit the impact of CMPK2 loss in vivo. Third, species-specific differences between human and mouse macrophages may contribute to the observed divergence. Finally, incomplete loss of *Cmpk2* driven by the LysM-Cre transgene, which has been previously reported (23), may have left sufficient residual CMPK2 expression in vivo to preserve cell-intrinsic antimicrobial activity. These possibilities are not mutually exclusive and underscore the challenges of linking cell-intrinsic phenotypes to in vivo outcomes in tuberculosis.

We did not directly assess the role of mitochondrial nucleotide metabolism or cytosolic DNA sensing pathways in this study. CMPK2 has been implicated in both processes in other contexts, and it is plausible that these mechanisms contribute to the phenotypes observed here. However, our data do not distinguish between these possibilities, and further work will be required to define the mechanistic basis of CMPK2 function during Mtb infection.

In summary, our findings identify CMPK2 as a regulator of human macrophage responses to Mtb and highlight a link between host metabolic pathways and innate immune gene expression. These results provide a foundation for future studies aimed at defining how nucleotide metabolism shapes host-pathogen interactions during tuberculosis.

## MATERIALS AND METHODS

### Human cells and culture

THP1 cells were cultured in RPMI supplemented with 10% FBS, 1% HEPES, and 1% sodium pyruvate. Prior to use in assays, THP1 cells were differentiated for two days in 100ng/mL PMA and allowed to rest after a media change to RPMI without PMA for an additional two days. Human MDMs were isolated using Sepmate 50 tubes (StemCell catalog #85450) and the accompanying protocol. CD14^+^ cells were selected using human CD14 micro beads (Miltenyi catalog #130-050-201) with MACS LS columns and quadroMACS separation magnet (Miltenyi) using the provided protocol. MDMs were obtained from buffy coats from anonymous donors provided by a local blood bank. MDMs were differentiated for 24 hours in RPMI supplemented with 10% FBS, 10% heat inactivated donor-specific serum isolated from blood, 1% HEPES, 1% sodium pyruvate, and 50ng/mL GM-CSF (PeproTech catalog #300-03). Media was then changed to RPMI supplemented with 10% FBS, 1% HEPES, 1% sodium pyruvate, and 50ng/mL GM-CSF and cells were incubated for three days. Finally, the media was changed to RPMI supplemented with 10% FBS, 1% HEPES, and 1% sodium pyruvate and cells were allowed to incubate for three additional days before use in experiments.

### CMPK2 KO cell line

The CMPK2 KO THP1 cell line (13) was a generous gift from the lab of Dr. Ling Jun Ho at the Taiwanese Institute of Cellular and System Medicine.

#### Bacterial strains, culture, and infection

Prior to use in assays, either Erdman strain *M. tuberculosis* or Mtb-lux (Erdman strain *M. tuberculosis* constitutively expressing the complete lux luciferase cassette, Addgene plasmid #26161) bacteria were grown to an OD600 of 0.5-0.8 at 37°C on a roller apparatus in 7H9 media supplemented with 20% Middlebrook Oleic Albumin Dextrose Catalase Growth Supplement (OADC), 1% glycerol, and 0.1% Tween 80. To prepare a single-cell bacterial suspension for infection, bacteria were washed with PBS and pelleted three times, sonicated at 90% amplitude for three 7s on/7s off cycles, and then, centrifuged at 300g for 5 minutes to sediment bacterial aggregates. The supernatant of this spin was collected and its OD600 was determined to prepare inoculum. For MOI calculations, we assumed that OD600=1 is equivalent to 3 × 10^8^ bacteria/mL. The volume of bacterial suspension needed for the desired MOI was resuspended in RPMI medium supplemented with 10% FBS, 100 mM HEPES, and 10 mM sodium pyruvate before being added to cells. Cells were incubated with bacteria for 1 hour at 37°C and 5% CO2 to allow for bacterial phagocytosis. Cells were then washed with PBS to remove extracellular bacteria and provided with fresh RPMI medium supplemented with 10% FBS, 100 mM HEPES, and 10 mM sodium pyruvate.

### Luminescent Mtb-lux infections

THP1 cells were differentiated in triplicate in a 96-well plate at 70,000 cells per well and infected at an MOI of three with Mtb-lux. Luminescence readings were taken at days 0, 2, and 4 after a PBS wash and media change. This experiment was performed across two independent biological replicates containing three individually infected wells each.

### CFU assays

THP1 macrophages were differentiated in 48-well plates at 250,000 cells per well and infected at an MOI of one in three independent biological replicates, each containing three technical replicates. After removing growth media, cells were lysed at the indicated time-points with 0.5% Triton X-100 (Sigma) in water and serial dilutions were plated in technical triplicate on 7H11 plates. Plates were allowed to grow for 3-4 weeks at 37°C before being counted.

### Antibodies and immunoblotting

The following antibodies were used in this study: CMPK2 (1:1000, Abcam ab194567); beta-Actin (1:5000, Santa Cruz SC-47778); Western blots were run using a 12% gel and transferred to PVDF membranes using the mixed molecular weight setting on a Biorad semidry transfer apparatus. Membranes were blocked for 1 hour in TBST+ 5% fat-free milk and incubated with primary antibody overnight. The following day, membranes were washed with TBST, incubated in TBST + 5% milk with secondary antibody for one hour, washed again, and imaged on a BioRad ChemiDoc imaging system.

### RNA isolation and qPCR

RNA was collected from cells by lysing them in TRIzol reagent (ThermoFisher catalog #15596026). RNA was then isolated using chloroform extraction according to the reagent user guide. cDNA was synthesized using the iScript cDNA synthesis kit (BioRad catalog # 1708890) according to manufacturer’s instructions. qPCR was performed on an Applied Biosystems 7500 Fast Real-time PCR system using Fast SYBR Green Master Mix (ThermoFisher catalog # 4385612) according to the provided reagent protocol.

### RNA Sequencing

RNA sequencing was performed on WT THP1 and CMPK2 KO cells 4 hours post infection with *M. tuberculosis* at an MOI of 5. RNA was extracted as above. Samples were run on an Agilent Tapestation 4200 to determine level of degradation and ensure only high quality RNA was used (RIN Score 8 or higher). A Qubit fluorimeter was used to determine the concentration prior to staring library prep. One microgram of total DNAse treated RNA was then prepared with the TruSeq Stranded mRNA Library Prep Kit (Illumina). Poly-A RNA was purified and fragmented before strand specific cDNA synthesis. cDNA were then a-tailed and indexed adapters were ligated. After adapter ligation, samples were PCR amplified and purified with AmpureXP beads, then validated again on an Agilent Tapestation 4200. Before being normalized and pooled, samples were quantified by Qubit then run on the Illumina NextSeq 2000. Analysis of RNA sequencing data was performed using iDEP2.0 with the quality control cutoff of 5 CPM in at least six libraries (24). EdgeR was used to transform counts data for clustering & PCA. Signaling and immune pathways depicted are curated by KEGG and visualized using Pathview (25-27). Genes marked as changing in expression were required to have a two-fold increase or decrease in RNA transcripts. Functional enrichment analysis was performed on all genes upregulated or downregulated by at least two-fold in *CMPK2* KO THP1 cells as compared to WT THP1 cells 4 h.p.i. using the default settings on the DAVID bioinformatics web server (28, 29)

### Mouse lines

The C57BL/6J CMPK2^fl/fl^ mouse line was a generous gift from the laboratory of Dr. Zhenyu Zhong at UTSW. The CMPK2^fl/fl LysMCre/+^ line was generated by crossing the CMPK2^fl/fl^ mouse line with a C57BL/6J LysMCre homozygous mouse line.

### Aerosol infections

Mtb aerosol infection was performed as described (5). Briefly, cultures were washed twice with PBS, sonicated, and resuspended in PBS at an OD600 of 0.1. Bacterial suspension was introduced into the nebulizer of a GlasCol aerosol chamber to infect mice with the desired number of bacteria per animal. On day zero of infection in CFU assays, lungs were collected from 3 mice and the entire lung homogenate was plated on 7H11 plates to determine the initial inoculum. In subsequent time points, left lungs, livers and spleens were collected, homogenized and plated on 7H11 plates for quantitation of CFUs at the indicated times post infection. For survival studies, infected mice were monitored and sacrificed when they had lost 15% of their maximal body weight.

### Statistical Analysis

Statistical analysis was performed using GraphPad Prism software (version 10). After verifying data normality, the unpaired Student’s t-test, or the Mann-Whitney test was used to compare two groups. Comparisons among three or more groups were assessed with one-way ANOVA. Differences among groups were considered statistically significant when p < 0.05, (*) p < 0.01 (**), p < 0.001(***), and p < 0.0001 (****).

## Acknowledgments

The authors would like to thank Dr. Ling Jun Ho for providing the THP1 *CMPK2* KO cell line. We thank the UTSW Genomics Sequencing Core for bulk RNA-seq library preparation and sequencing.

## Data availability

All RNA-Seq data have been deposited in the Gene Expression Omnibus (GEO) under accession number: GSE331078.

## Author contributions

Conceptualization, J.N., M.U.S.; Formal analysis, J.N., M.U.S.; Investigation, J.N., V.A.E., P.C.C., K.F.N. B.R.S.D., K.C.R.; Funding acquisition, J.N., M.U.S.; Project administration, M.U.S.; Resources, Z.Z.; Supervision, M.U.S.; Writing - original draft, J.N., M.U.S.; Writing - review and editing, J.N., Z.Z., M.U.S..

