## Supplemental Figures for "CMPK2 restricts *Mycobacterium tuberculosis* replication and regulates macrophage gene expression"

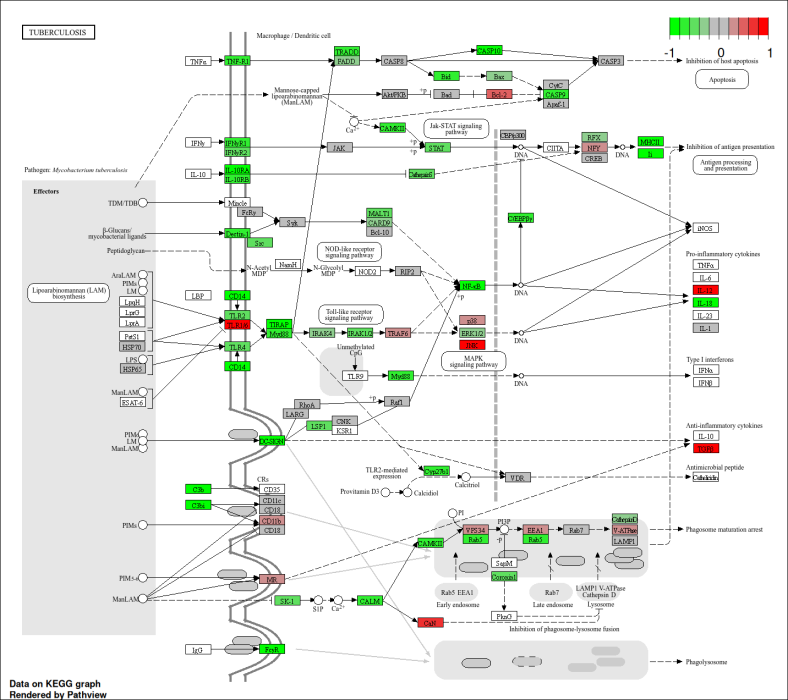

**Supplementary Figure 1. Differential gene expression mapped onto innate immune pathways during Mtb infection.** KEGG pathway map of host innate immune responses to Mtb, with genes identified as differentially expressed in Mtb-infected CMPK2 knockout cells relative to wild-type THP1 cells by RNA sequencing. Genes with increased expression are shown in red, and genes with reduced expression are shown in green.

­­­­­­
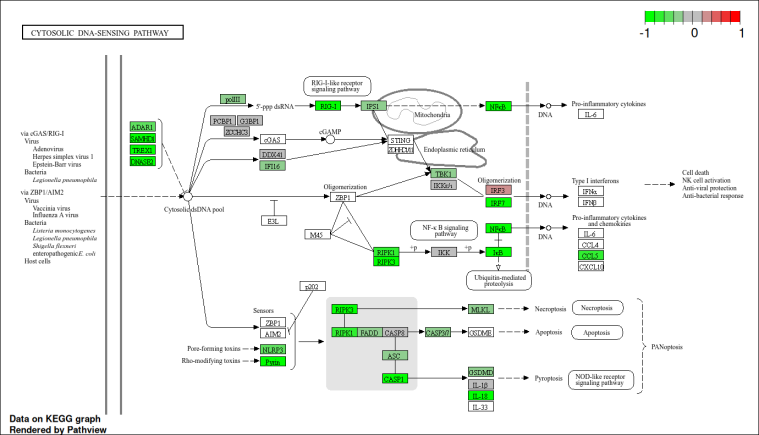

**Supplementary Figure 2. Differential expression of genes associated with DNA sensing and innate immune pathways.** KEGG pathway map highlighting genes associated with DNA sensing and related innate immune responses, with differential expression in Mtb-infected THP1 *CMPK2* knockout cells relative to wild-type THP1 cells. Genes with increased expression are shown in red, and genes with reduced expression are shown in green.

**Supplemental Table 1. Functional enrichment analysis of genes downregulated in THP1 *CMPK2* knockout cells.** Functional enrichment results generated using the DAVID bioinformatics webserver for genes downregulated by at least two-fold in THP1 *CMPK2* knockout cells compared to wild-type cells at 4 hours post-infection. Enrichment significance was calculated using default DAVID parameters.

**Supplementary Table 2: Functional enrichment analysis of genes upregulated in THP1 *CMPK2* knockout cells.** Functional enrichment results generated using the DAVID bioinformatics webserver for genes upregulated by at least two-fold in THP1 *CMPK2* knockout cells compared to wild-type cells at 4 hours post-infection. Enrichment significance was calculated using default DAVID parameters.
