## Supplemental Table 1 for "CMPK2 restricts *Mycobacterium tuberculosis* replication and regulates macrophage gene expression"

| Cluster | Cluster Enrichment Score | Category | Term | Count | List Total | Pop Hits | Pop Total | % |
|---|---|---|---|---|---|---|---|---|
| 1 | 14.52 | GOTERM_BP_DIRECT | immune system process | 66 | 409 | 953 | 19512 | 15.1[illegible] |
| 1 | 14.52 | UP_KW_BIOLOGICAL_PROCESS | Innate immunity | 44 | 278 | 432 | 11605 | 10.0[illegible] |
| 1 | 14.52 | GOTERM_BP_DIRECT | innate immune response | 50 | 409 | 621 | 19512 | 11.4[illegible] |
| 1 | 14.52 | GOTERM_BP_DIRECT | defense response to virus | 32 | 409 | 256 | 19512 | 7.34[illegible] |
| 1 | 14.52 | UP_KW_BIOLOGICAL_PROCESS | Immunity | 65 | 278 | 988 | 11605 | 14.9[illegible] |
| 1 | 14.52 | UP_KW_BIOLOGICAL_PROCESS | Antiviral defense | 24 | 278 | 139 | 11605 | 5.50[illegible] |
| 2 | 8.47 | UP_KW_BIOLOGICAL_PROCESS | Antiviral defense | 24 | 278 | 139 | 11605 | 5.50[illegible] |
| 2 | 8.47 | GOTERM_BP_DIRECT | negative regulation of viral genome replication | 10 | 409 | 46 | 19512 | 2.29[illegible] |
| 2 | 8.47 | GOTERM_BP_DIRECT | response to virus | 15 | 409 | 129 | 19512 | 3.44[illegible] |
| 3 | 4.79 | GOTERM_CC_DIRECT | plasma membrane | 185 | 415 | 5927 | 20808 | 42.4[illegible] |
| 3 | 4.79 | UP_SEQ_FEATURE | CARBOHYD:N-linked (GlcNAc...) asparagine | 140 | 422 | 4425 | 20669 | 32.1[illegible] |
| 3 | 4.79 | GOTERM_CC_DIRECT | membrane | 233 | 415 | 9038 | 20808 | 53.4[illegible] |
| 3 | 4.79 | UP_SEQ_FEATURE | TOPO_DOM:Extracellular | 97 | 422 | 3012 | 20669 | 22.2[illegible] |
| 3 | 4.79 | UP_KW_CELLULAR_COMPONENT | Cell membrane | 128 | 387 | 4200 | 18195 | 29.3[illegible] |
| 3 | 4.79 | UP_KW_PTM | Disulfide bond | 133 | 343 | 4060 | 14452 | 30.5[illegible] |
| 3 | 4.79 | UP_SEQ_FEATURE | TOPO_DOM:Cytoplasmic | 116 | 422 | 3928 | 20669 | 26.6[illegible] |
| 3 | 4.79 | UP_KW_CELLULAR_COMPONENT | Membrane | 215 | 387 | 8428 | 18195 | 49.3[illegible] |
| 3 | 4.79 | UP_KW_PTM | Glycoprotein | 148 | 343 | 4892 | 14452 | 33.9[illegible] |
| 3 | 4.79 | UP_KW_DOMAIN | Signal | 118 | 313 | 4426 | 14656 | 27.0[illegible] |
| 3 | 4.79 | UP_SEQ_FEATURE | TRANSMEM:Helical | 135 | 422 | 5434 | 20669 | 30.9[illegible] |
| 3 | 4.79 | UP_KW_DOMAIN | Transmembrane helix | 146 | 313 | 5895 | 14656 | 33.4[illegible] |
| 3 | 4.79 | UP_KW_DOMAIN | Transmembrane | 146 | 313 | 5971 | 14656 | 33.4[illegible] |
| 4 | 4.14 | KEGG_PATHWAY | Staphylococcus aureus infection | 14 | 253 | 102 | 9496 | 3.21[illegible] |
| 4 | 4.14 | GOTERM_MF_DIRECT | IgG binding | 5 | 417 | 16 | 19272 | 1.15[illegible] |
| 4 | 4.14 | KEGG_PATHWAY | Systemic lupus erythematosus | 13 | 253 | 144 | 9496 | 2.98[illegible] |
| 5 | 4.14 | GOTERM_CC_DIRECT | extracellular region | 90 | 415 | 2693 | 20808 | 20.6[illegible] |
| 5 | 4.14 | UP_KW_CELLULAR_COMPONENT | Secreted | 74 | 387 | 2233 | 18195 | 16.9[illegible] |
| 5 | 4.14 | GOTERM_CC_DIRECT | extracellular space | 62 | 415 | 1895 | 20808 | 14.2[illegible] |

| Cluster | Cluster Enrichment Score | Category | Term | Count | List Total | Pop Hits | Pop Total | % |
|---|---|---|---|---|---|---|---|---|
| 5 | 4.14 | UP_KW_DOMAIN | Signal | 118 | 313 | 4426 | 14656 | 27.0[illegible] |
| 6 | 4.13 | GOTERM_BP_DIRECT | lipopolysaccharide-mediated signaling pathway | 9 | 409 | 38 | 19512 | 2.06[illegible] |
| 6 | 4.13 | GOTERM_MF_DIRECT | lipopolysaccharide immune receptor activity | 4 | 417 | 7 | 19272 | 0.92[illegible] |
| 6 | 4.13 | GOTERM_BP_DIRECT | positive regulation of lipopolysaccharide-mediated signaling pathway | 4 | 409 | 11 | 19512 | 0.92[illegible] |
| 7 | 3.76 | GOTERM_CC_DIRECT | lysosome | 27 | 415 | 532 | 20808 | 6.19[illegible] |
| 7 | 3.76 | UP_KW_CELLULAR_COMPONENT | Lysosome | 24 | 387 | 450 | 18195 | 5.50[illegible] |
| 7 | 3.76 | GOTERM_CC_DIRECT | lysosomal membrane | 19 | 415 | 418 | 20808 | 4.36[illegible] |
| 8 | 3.38 | GOTERM_CC_DIRECT | cytosol | 156 | 415 | 5677 | 20808 | 35.7[illegible] |
| 8 | 3.38 | GOTERM_CC_DIRECT | cytoplasm | 190 | 415 | 7749 | 20808 | 43.5[illegible] |
| 8 | 3.38 | UP_KW_CELLULAR_COMPONENT | Cytoplasm | 141 | 387 | 5979 | 18195 | 32.3[illegible] |
| 9 | 3.27 | KEGG_PATHWAY | Leishmaniasis | 11 | 253 | 79 | 9496 | 2.52[illegible] |
| 9 | 3.27 | KEGG_PATHWAY | Tuberculosis | 14 | 253 | 182 | 9496 | 3.21[illegible] |
| 9 | 3.27 | KEGG_PATHWAY | Phagosome | 12 | 253 | 159 | 9496 | 2.75[illegible] |
| 10 | 3.11 | UP_KW_PTM | Zymogen | 18 | 343 | 227 | 14452 | 4.13[illegible] |
| 10 | 3.11 | INTERPRO | TRYPSIN_HIS | 11 | 427 | 109 | 20827 | 2.52[illegible] |
| 10 | 3.11 | INTERPRO | Peptidase_S1A | 11 | 427 | 118 | 20827 | 2.52[illegible] |
| 10 | 3.11 | INTERPRO | TRYPSIN_SER | 10 | 427 | 99 | 20827 | 2.29[illegible] |
| 10 | 3.11 | UP_SEQ_FEATURE | DOMAIN:Peptidase S1 | 11 | 422 | 123 | 20669 | 2.52[illegible] |
| 10 | 3.11 | INTERPRO | Trypsin_dom | 11 | 427 | 126 | 20827 | 2.52[illegible] |
| 10 | 3.11 | INTERPRO | Peptidase_S1_PA_chymotrypsin | 11 | 427 | 127 | 20827 | 2.52[illegible] |
| 10 | 3.11 | INTERPRO | Peptidase_S1_PA | 11 | 427 | 135 | 20827 | 2.52[illegible] |
| 10 | 3.11 | SMART | Tryp_SPc | 11 | 246 | 123 | 10704 | 2.52[illegible] |
| 10 | 3.11 | GOTERM_MF_DIRECT | hydrolase activity | 61 | 417 | 1847 | 19272 | 13.9[illegible] |
| 10 | 3.11 | GOTERM_BP_DIRECT | proteolysis | 27 | 409 | 671 | 19512 | 6.19[illegible] |
| 10 | 3.11 | UP_KW_MOLECULAR_FUNCTION | Serine protease | 11 | 284 | 147 | 12059 | 2.52[illegible] |
| 10 | 3.11 | GOTERM_MF_DIRECT | serine-type endopeptidase activity | 12 | 417 | 192 | 19272 | 2.75[illegible] |
| 10 | 3.11 | GOTERM_MF_DIRECT | peptidase activity | 23 | 417 | 544 | 19272 | 5.28[illegible] |

| Cluster | Cluster Enrichment Score | Category | Term | Count | List Total | Pop Hits | Pop Total | % |
|---|---|---|---|---|---|---|---|---|
| 10 | 3.11 | UP_SEQ_FEATURE | ACT_SITE:Charge relay system | 12 | 422 | 209 | 20669 | 2.7[illegible] |
| 10 | 3.11 | UP_KW_MOLECULAR_FUNCTION | Protease | 25 | 284 | 586 | 12059 | 5.7[illegible] |
| 10 | 3.11 | UP_KW_MOLECULAR_FUNCTION | Hydrolase | 63 | 284 | 1940 | 12059 | 14.[illegible] |
| 10 | 3.11 | GOTERM_MF_DIRECT | serine-type peptidase activity | 10 | 417 | 152 | 19272 | 2.2[illegible] |
| 11 | 2.99 | INTERPRO | Ig_sub | 27 | 427 | 528 | 20827 | 6.1[illegible] |
| 11 | 2.99 | INTERPRO | Ig_sub2 | 17 | 427 | 262 | 20827 | 3.9[illegible] |
| 11 | 2.99 | SMART | IG | 27 | 246 | 528 | 10704 | 6.1[illegible] |
| 11 | 2.99 | INTERPRO | Ig-like_dom_sf | 35 | 427 | 869 | 20827 | 8.0[illegible] |
| 11 | 2.99 | INTERPRO | Ig-like_fold | 42 | 427 | 1123 | 20827 | 9.6[illegible] |
| 11 | 2.99 | SMART | IGc2 | 17 | 246 | 262 | 10704 | 3.9[illegible] |
| 11 | 2.99 | UP_SEQ_FEATURE | DOMAIN:Ig-like V-type | 11 | 422 | 139 | 20669 | 2.5[illegible] |
| 11 | 2.99 | INTERPRO | Ig-like_dom | 31 | 427 | 768 | 20827 | 7.1[illegible] |
| 11 | 2.99 | UP_SEQ_FEATURE | DOMAIN:Ig-like C2-type 3 | 10 | 422 | 133 | 20669 | 2.2[illegible] |
| 11 | 2.99 | UP_SEQ_FEATURE | DOMAIN:Ig-like C2-type 1 | 13 | 422 | 219 | 20669 | 2.9[illegible] |
| 11 | 2.99 | UP_SEQ_FEATURE | DOMAIN:Ig-like C2-type 2 | 13 | 422 | 219 | 20669 | 2.9[illegible] |
| 11 | 2.99 | INTERPRO | Ig_I-set | 10 | 427 | 143 | 20827 | 2.2[illegible] |
| 11 | 2.99 | UP_KW_DOMAIN | Immunoglobulin domain | 31 | 313 | 828 | 14656 | 7.1[illegible] |
| 11 | 2.99 | UP_SEQ_FEATURE | DOMAIN:Ig-like | 24 | 422 | 676 | 20669 | 5.5[illegible] |
| 11 | 2.99 | INTERPRO | Ig_V-set | 12 | 427 | 484 | 20827 | 2.7[illegible] |
| 12 | 2.57 | GOTERM_BP_DIRECT | cellular response to cadmium ion | 6 | 409 | 23 | 19512 | 1.3[illegible] |
| 12 | 2.57 | INTERPRO | Metalthion_vert_metal_BS | 4 | 427 | 10 | 20827 | 0.9[illegible] |
| 12 | 2.57 | KEGG_PATHWAY | Mineral absorption | 8 | 253 | 61 | 9496 | 1.8[illegible] |
| 12 | 2.57 | UP_KW_LIGAND | Cadmium | 4 | 165 | 10 | 7011 | 0.9[illegible] |
| 12 | 2.57 | UP_SEQ_FEATURE | REGION:Alpha | 4 | 422 | 12 | 20669 | 0.9[illegible] |
| 12 | 2.57 | UP_SEQ_FEATURE | REGION:Beta | 4 | 422 | 12 | 20669 | 0.9[illegible] |
| 12 | 2.57 | GOTERM_BP_DIRECT | cellular response to copper ion | 5 | 409 | 26 | 19512 | 1.1[illegible] |
| 12 | 2.57 | INTERPRO | Metalthion_dom_sf | 4 | 427 | 14 | 20827 | 0.9[illegible] |
| 12 | 2.57 | INTERPRO | Metalthion_dom_sf_vert | 4 | 427 | 14 | 20827 | 0.9[illegible] |
| 12 | 2.57 | INTERPRO | Metalthion_vert | 4 | 427 | 14 | 20827 | 0.9[illegible] |

| Cluster | Cluster Enrichment Score | Category | Term | Count | List Total | Pop Hits | Pop Total | % |
|---|---|---|---|---|---|---|---|---|
| 12 | 2.57 | UP_KW_LIGAND | Metal-thiolate cluster | 4 | 165 | 13 | 7011 | 0.92 |
| 12 | 2.57 | GOTERM_BP_DIRECT | detoxification of copper ion | 4 | 409 | 16 | 19512 | 0.92 |
| 12 | 2.57 | GOTERM_BP_DIRECT | negative regulation of growth | 4 | 409 | 18 | 19512 | 0.92 |
| 12 | 2.57 | GOTERM_BP_DIRECT | intracellular zinc ion homeostasis | 5 | 409 | 35 | 19512 | 1.15 |
| 12 | 2.57 | GOTERM_BP_DIRECT | cellular response to zinc ion | 4 | 409 | 25 | 19512 | 0.92 |
| 12 | 2.57 | UP_KW_LIGAND | Copper | 5 | 165 | 71 | 7011 | 1.15 |
| 13 | 2.31 | UP_SEQ_FEATURE | DOMAIN:bZIP | 7 | 422 | 55 | 20669 | 1.61 |
| 13 | 2.31 | INTERPRO | bZIP_sf | 7 | 427 | 55 | 20827 | 1.61 |
| 13 | 2.31 | INTERPRO | bZIP | 7 | 427 | 56 | 20827 | 1.61 |
| 13 | 2.31 | SMART | BRLZ | 7 | 246 | 53 | 10704 | 1.61 |
| 13 | 2.31 | UP_SEQ_FEATURE | REGION:Leucine-zipper | 9 | 422 | 118 | 20669 | 2.06 |
| 13 | 2.31 | GOTERM_CC_DIRECT | RNA polymerase II transcription regulator complex | 9 | 415 | 135 | 20808 | 2.06 |
| 13 | 2.31 | UP_SEQ_FEATURE | REGION:Basic motif | 6 | 422 | 57 | 20669 | 1.38 |
| 13 | 2.31 | INTERPRO | AP-1 | 3 | 427 | 9 | 20827 | 0.69 |
| 13 | 2.31 | UP_SEQ_FEATURE | DOMAIN:BZIP | 5 | 422 | 46 | 20669 | 1.15 |
| 13 | 2.31 | GOTERM_BP_DIRECT | integrated stress response signaling | 4 | 409 | 25 | 19512 | 0.92 |
| 13 | 2.31 | GOTERM_BP_DIRECT | fat cell differentiation | 5 | 409 | 95 | 19512 | 1.15 |
| 14 | 2.22 | GOTERM_BP_DIRECT | Fc-gamma receptor signaling pathway | 5 | 409 | 15 | 19512 | 1.15 |
| 14 | 2.22 | GOTERM_MF_DIRECT | IgG binding | 5 | 417 | 16 | 19272 | 1.15 |
| 14 | 2.22 | GOTERM_MF_DIRECT | IgG receptor activity | 3 | 417 | 8 | 19272 | 0.69 |
| 14 | 2.22 | KEGG_PATHWAY | Fc gamma R-mediated phagocytosis | 8 | 253 | 99 | 9496 | 1.83 |
| 14 | 2.22 | GOTERM_BP_DIRECT | antibody-dependent cellular cytotoxicity | 3 | 409 | 10 | 19512 | 0.69 |
| 14 | 2.22 | UP_KW_MOLECULAR_FUNCTION | IgG-binding protein | 3 | 284 | 10 | 12059 | 0.69 |
| 14 | 2.22 | INTERPRO | Ig_Fc_receptor | 3 | 427 | 20 | 20827 | 0.69 |
| 15 | 2.20 | UP_KW_LIGAND | NADP | 12 | 165 | 186 | 7011 | 2.75 |
| 15 | 2.20 | GOTERM_MF_DIRECT | oxidoreductase activity | 25 | 417 | 625 | 19272 | 5.73 |

| Cluster | Cluster Enrichment Score | Category | Term | Count | List Total | Pop Hits | Pop Total | % |
|---|---|---|---|---|---|---|---|---|
| 15 | 2.20 | UP_KW_MOLECULAR_FUNCTION | Oxidoreductase | 25 | 284 | 627 | 12059 | 5.73[illegible] |
| 16 | 2.15 | GOTERM_BP_DIRECT | pyridine nucleotide biosynthetic process | 4 | 409 | 11 | 19512 | 0.92[illegible] |
| 16 | 2.15 | UP_KW_BIOLOGICAL_PROCESS | Pyridine nucleotide biosynthesis | 4 | 278 | 11 | 11605 | 0.92[illegible] |
| 16 | 2.15 | GOTERM_BP_DIRECT | NAD+ biosynthetic process | 4 | 409 | 13 | 19512 | 0.92[illegible] |
| 16 | 2.15 | GOTERM_BP_DIRECT | 'de novo' NAD+ biosynthetic process from L-tryptophan | 3 | 409 | 9 | 19512 | 0.69[illegible] |
| 16 | 2.15 | KEGG_PATHWAY | Biosynthesis of cofactors | 7 | 253 | 154 | 9496 | 1.61[illegible] |
| 17 | 2.08 | KEGG_PATHWAY | Pertussis | 11 | 253 | 78 | 9496 | 2.52[illegible] |
| 17 | 2.08 | GOTERM_CC_DIRECT | symbiont cell surface | 5 | 415 | 20 | 20808 | 1.15[illegible] |
| 17 | 2.08 | KEGG_PATHWAY | Complement and coagulation cascades | 9 | 253 | 88 | 9496 | 2.06[illegible] |
| 17 | 2.08 | KEGG_PATHWAY | Alcoholic liver disease | 11 | 253 | 144 | 9496 | 2.52[illegible] |
| 17 | 2.08 | UP_KW_BIOLOGICAL_PROCESS | Complement pathway | 4 | 278 | 36 | 11605 | 0.92[illegible] |
| 17 | 2.08 | GOTERM_BP_DIRECT | complement activation, classical pathway | 4 | 409 | 45 | 19512 | 0.92[illegible] |
| 17 | 2.08 | KEGG_PATHWAY | Chagas disease | 6 | 253 | 103 | 9496 | 1.38[illegible] |
| 17 | 2.08 | GOTERM_BP_DIRECT | complement activation | 3 | 409 | 38 | 19512 | 0.69[illegible] |
| 18 | 1.95 | UP_SEQ_FEATURE | BINDING:axial binding residue | 9 | 422 | 114 | 20669 | 2.06[illegible] |
| 18 | 1.95 | UP_KW_LIGAND | Heme | 9 | 165 | 142 | 7011 | 2.06[illegible] |
| 18 | 1.95 | GOTERM_MF_DIRECT | heme binding | 9 | 417 | 154 | 19272 | 2.06[illegible] |
| 18 | 1.95 | UP_KW_LIGAND | Iron | 16 | 165 | 357 | 7011 | 3.67[illegible] |
| 19 | 1.88 | GOTERM_BP_DIRECT | cellular response to interleukin-1 | 6 | 409 | 56 | 19512 | 1.38[illegible] |
| 19 | 1.88 | UP_SEQ_FEATURE | REGION:GTPase domain (Globular) | 3 | 422 | 7 | 20669 | 0.69[illegible] |
| 19 | 1.88 | INTERPRO | GBP_C | 3 | 427 | 7 | 20827 | 0.69[illegible] |
| 19 | 1.88 | INTERPRO | Guanylate-bd/ATL_C | 3 | 427 | 9 | 20827 | 0.69[illegible] |
| 19 | 1.88 | GOTERM_BP_DIRECT | cytolysis in another organism | 3 | 409 | 9 | 19512 | 0.69[illegible] |
| 19 | 1.88 | GOTERM_BP_DIRECT | positive regulation of pyroptotic inflammatory response | 3 | 409 | 9 | 19512 | 0.69[illegible] |
| 19 | 1.88 | INTERPRO | Guanylate-bd_C_sf | 3 | 427 | 10 | 20827 | 0.69[illegible] |

| Cluster | Cluster Enrichment Score | Category | Term | Count | List Total | Pop Hits | Pop Total | % |
|---|---|---|---|---|---|---|---|---|
| 19 | 1.88 | UP_SEQ_FEATURE | DOMAIN:GB1/RHD3-type G | 3 | 422 | 11 | 20669 | 0.69 |
| 19 | 1.88 | INTERPRO | Guanylate-bd_N | 3 | 427 | 11 | 20827 | 0.69 |
| 19 | 1.88 | INTERPRO | G_GB1_RHD3_dom | 3 | 427 | 11 | 20827 | 0.69 |
| 20 | 1.84 | GOTERM_BP_DIRECT | amino acid biosynthetic process | 7 | 409 | 27 | 19512 | 1.61 |
| 20 | 1.84 | UP_KW_BIOLOGICAL_PROCESS | Amino-acid biosynthesis | 7 | 278 | 27 | 11605 | 1.61 |
| 20 | 1.84 | GOTERM_BP_DIRECT | L-serine metabolic process | 3 | 409 | 10 | 19512 | 0.69 |
| 20 | 1.84 | KEGG_PATHWAY | Cysteine and methionine metabolism | 5 | 253 | 52 | 9496 | 1.15 |
| 20 | 1.84 | KEGG_PATHWAY | Biosynthesis of amino acids | 6 | 253 | 75 | 9496 | 1.38 |
| 20 | 1.84 | KEGG_PATHWAY | Alanine, aspartate and glutamate metabolism | 4 | 253 | 37 | 9496 | 0.92 |
| 20 | 1.84 | KEGG_PATHWAY | Glycine, serine and threonine metabolism | 4 | 253 | 41 | 9496 | 0.92 |
| 20 | 1.84 | GOTERM_MF_DIRECT | pyridoxal phosphate binding | 3 | 417 | 58 | 19272 | 0.69 |
| 20 | 1.84 | UP_KW_LIGAND | Pyridoxal phosphate | 3 | 165 | 63 | 7011 | 0.69 |
| 21 | 1.60 | GOTERM_CC_DIRECT | collagen trimer | 9 | 415 | 104 | 20808 | 2.06 |
| 21 | 1.60 | UP_KW_DOMAIN | Collagen | 9 | 313 | 102 | 14656 | 2.06 |
| 21 | 1.60 | INTERPRO | Collagen | 7 | 427 | 86 | 20827 | 1.61 |
| 21 | 1.60 | INTERPRO | Collagen/C1q_domain | 4 | 427 | 28 | 20827 | 0.92 |
| 21 | 1.60 | UP_SEQ_FEATURE | DOMAIN:C1q | 4 | 422 | 36 | 20669 | 0.92 |
| 21 | 1.60 | INTERPRO | C1q_dom | 4 | 427 | 37 | 20827 | 0.92 |
| 21 | 1.60 | SMART | C1Q | 4 | 246 | 35 | 10704 | 0.92 |
| 21 | 1.60 | INTERPRO | Tumour_necrosis_fac-like_dom | 4 | 427 | 58 | 20827 | 0.92 |
| 21 | 1.60 | UP_SEQ_FEATURE | DOMAIN:Collagen-like | 3 | 422 | 33 | 20669 | 0.69 |
| 21 | 1.60 | UP_KW_PTM | Hydroxylation | 7 | 343 | 210 | 14452 | 1.61 |
| 22 | 1.58 | INTERPRO | SH3-like_dom_sf | 11 | 427 | 220 | 20827 | 2.52 |
| 22 | 1.58 | INTERPRO | SH3_domain | 11 | 427 | 229 | 20827 | 2.52 |
| 22 | 1.58 | UP_KW_DOMAIN | SH3 domain | 11 | 313 | 229 | 14656 | 2.52 |
| 22 | 1.58 | UP_SEQ_FEATURE | DOMAIN:SH3 | 10 | 422 | 217 | 20669 | 2.29 |
| 22 | 1.58 | SMART | SH3 | 10 | 246 | 208 | 10704 | 2.29 |

| Cluster | Cluster Enrichment Score | Category | Term | Count | List Total | Pop Hits | Pop Total | % |
|---|---|---|---|---|---|---|---|---|
| 23 | 1.55 | GOTERM_BP_DIRECT | L-amino acid transport | 3 | 409 | 6 | 19512 | 0.69 |
| 23 | 1.55 | GOTERM_MF_DIRECT | neutral L-amino acid transmembrane transporter activity | 4 | 417 | 18 | 19272 | 0.92 |
| 23 | 1.55 | GOTERM_MF_DIRECT | L-leucine transmembrane transporter activity | 3 | 417 | 6 | 19272 | 0.69 |
| 23 | 1.55 | GOTERM_BP_DIRECT | neutral amino acid transport | 4 | 409 | 23 | 19512 | 0.92 |
| 23 | 1.55 | GOTERM_MF_DIRECT | L-amino acid transmembrane transporter activity | 4 | 417 | 25 | 19272 | 0.92 |
| 23 | 1.55 | GOTERM_BP_DIRECT | L-leucine transport | 3 | 409 | 10 | 19512 | 0.69 |
| 23 | 1.55 | GOTERM_BP_DIRECT | amino acid transmembrane transport | 4 | 409 | 30 | 19512 | 0.92 |
| 23 | 1.55 | UP_KW_BIOLOGICAL_PROCESS | Amino-acid transport | 5 | 278 | 55 | 11605 | 1.15 |
| 23 | 1.55 | GOTERM_CC_DIRECT | basal plasma membrane | 5 | 415 | 73 | 20808 | 1.15 |
| 23 | 1.55 | GOTERM_BP_DIRECT | amino acid transport | 5 | 409 | 70 | 19512 | 1.15 |
| 23 | 1.55 | GOTERM_BP_DIRECT | L-alpha-amino acid transmembrane transport | 3 | 409 | 19 | 19512 | 0.69 |
| 23 | 1.55 | GOTERM_MF_DIRECT | amino acid transmembrane transporter activity | 3 | 417 | 34 | 19272 | 0.69 |
| 23 | 1.55 | GOTERM_BP_DIRECT | transport across blood-brain barrier | 4 | 409 | 85 | 19512 | 0.92 |
| 24 | 1.54 | UP_SEQ_FEATURE | DOMAIN:SH2 | 7 | 422 | 110 | 20669 | 1.61 |
| 24 | 1.54 | INTERPRO | SH2 | 7 | 427 | 111 | 20827 | 1.61 |
| 24 | 1.54 | UP_KW_DOMAIN | SH2 domain | 7 | 313 | 109 | 14656 | 1.61 |
| 24 | 1.54 | INTERPRO | SH2_dom_sf | 7 | 427 | 116 | 20827 | 1.61 |
| 24 | 1.54 | SMART | SH2 | 7 | 246 | 105 | 10704 | 1.61 |
| 25 | 1.48 | GOTERM_MF_DIRECT | cytokine binding | 7 | 417 | 47 | 19272 | 1.61 |
| 25 | 1.48 | GOTERM_BP_DIRECT | pyroptotic inflammatory response | 3 | 409 | 41 | 19512 | 0.69 |
| 25 | 1.48 | GOTERM_MF_DIRECT | endopeptidase activity | 4 | 417 | 96 | 19272 | 0.92 |
| 26 | 1.37 | GOTERM_BP_DIRECT | cholesterol metabolic process | 8 | 409 | 109 | 19512 | 1.83 |
| 26 | 1.37 | UP_KW_BIOLOGICAL_PROCESS | Steroid metabolism | 8 | 278 | 109 | 11605 | 1.83 |
| 26 | 1.37 | GOTERM_BP_DIRECT | steroid metabolic process | 8 | 409 | 138 | 19512 | 1.83 |
| 26 | 1.37 | UP_KW_BIOLOGICAL_PROCESS | Lipid biosynthesis | 10 | 278 | 178 | 11605 | 2.29 |

| Cluster | Cluster Enrichment Score | Category | Term | Count | List Total | Pop Hits | Pop Total | % |
|---|---|---|---|---|---|---|---|---|
| 26 | 1.37 | GOTERM_BP_DIRECT | steroid biosynthetic process | 5 | 409 | 65 | 19512 | 1.15 |
| 26 | 1.37 | UP_KW_BIOLOGICAL_PROCESS | Steroid biosynthesis | 4 | 278 | 47 | 11605 | 0.92 |
| 26 | 1.37 | UP_KW_BIOLOGICAL_PROCESS | Sterol metabolism | 5 | 278 | 79 | 11605 | 1.15 |
| 26 | 1.37 | UP_KW_BIOLOGICAL_PROCESS | Cholesterol metabolism | 4 | 278 | 70 | 11605 | 0.92 |
| 27 | 1.31 | INTERPRO | Collagen | 7 | 427 | 86 | 20827 | 1.61 |
| 27 | 1.31 | UP_SEQ_FEATURE | DOMAIN:Collagen-like 1 | 3 | 422 | 29 | 20669 | 0.69 |
| 27 | 1.31 | UP_SEQ_FEATURE | DOMAIN:Collagen-like 2 | 3 | 422 | 29 | 20669 | 0.69 |
| 28 | 1.29 | GOTERM_BP_DIRECT | regulation of cell growth | 6 | 409 | 81 | 19512 | 1.38 |
| 28 | 1.29 | GOTERM_MF_DIRECT | insulin-like growth factor binding | 3 | 417 | 14 | 19272 | 0.69 |
| 28 | 1.29 | UP_SEQ_FEATURE | DOMAIN:IGFBP N-terminal | 3 | 422 | 20 | 20669 | 0.69 |
| 28 | 1.29 | INTERPRO | IGFBP-like | 3 | 427 | 20 | 20827 | 0.69 |
| 28 | 1.29 | SMART | IB | 3 | 246 | 18 | 10704 | 0.69 |
| 28 | 1.29 | INTERPRO | Growth_fac_rcpt_cys_sf | 7 | 427 | 145 | 20827 | 1.61 |
| 29 | 1.26 | KEGG_PATHWAY | Cell adhesion molecules | 13 | 253 | 160 | 9496 | 2.98 |
| 29 | 1.26 | GOTERM_MF_DIRECT | peptide antigen binding | 6 | 417 | 46 | 19272 | 1.38 |
| 29 | 1.26 | UP_SEQ_FEATURE | REGION:Connecting peptide | 5 | 422 | 38 | 20669 | 1.15 |
| 29 | 1.26 | INTERPRO | Ig/MHC_CS | 6 | 427 | 70 | 20827 | 1.38 |
| 29 | 1.26 | KEGG_PATHWAY | Allograft rejection | 5 | 253 | 39 | 9496 | 1.15 |
| 29 | 1.26 | KEGG_PATHWAY | Viral myocarditis | 6 | 253 | 70 | 9496 | 1.38 |
| 29 | 1.26 | KEGG_PATHWAY | Intestinal immune network for IgA production | 5 | 253 | 50 | 9496 | 1.15 |
| 29 | 1.26 | KEGG_PATHWAY | Autoimmune thyroid disease | 5 | 253 | 54 | 9496 | 1.15 |
| 29 | 1.26 | KEGG_PATHWAY | Rheumatoid arthritis | 6 | 253 | 95 | 9496 | 1.38 |
| 29 | 1.26 | KEGG_PATHWAY | Type I diabetes mellitus | 4 | 253 | 44 | 9496 | 0.92 |
| 29 | 1.26 | KEGG_PATHWAY | Graft-versus-host disease | 4 | 253 | 45 | 9496 | 0.92 |
| 29 | 1.26 | KEGG_PATHWAY | Antigen processing and presentation | 5 | 253 | 82 | 9496 | 1.15 |
| 29 | 1.26 | UP_SEQ_FEATURE | DOMAIN:Ig-like C1-type | 3 | 422 | 40 | 20669 | 0.69 |
| 29 | 1.26 | KEGG_PATHWAY | Asthma | 3 | 253 | 32 | 9496 | 0.69 |
| 29 | 1.26 | INTERPRO | MHC_I/II-like_Ag-recog | 3 | 427 | 47 | 20827 | 0.69 |

| Cluster | Cluster Enrichment Score | Category | Term | Count | List Total | Pop Hits | Pop Total | % |
|---|---|---|---|---|---|---|---|---|
| 29 | 1.26 | INTERPRO | Ig_C1-set | 3 | 427 | 78 | 20827 | 0.69[illegible] |
| 29 | 1.26 | SMART | IGc1 | 3 | 246 | 75 | 10704 | 0.69[illegible] |
| 30 | 1.22 | INTERPRO | Ald_DH_CS_CYS | 3 | 427 | 16 | 20827 | 0.69[illegible] |
| 30 | 1.22 | INTERPRO | Ald_DH_C | 3 | 427 | 19 | 20827 | 0.69[illegible] |
| 30 | 1.22 | INTERPRO | Ald_DH_N | 3 | 427 | 20 | 20827 | 0.69[illegible] |
| 30 | 1.22 | INTERPRO | Aldehyde_DH_dom | 3 | 427 | 20 | 20827 | 0.69[illegible] |
| 30 | 1.22 | INTERPRO | Ald_DH/histidinol_DH | 3 | 427 | 20 | 20827 | 0.69[illegible] |
| 30 | 1.22 | GOTERM_MF_DIRECT | oxidoreductase activity, acting on the aldehyde or oxo group of donors, NAD or NADP as acceptor | 3 | 417 | 23 | 19272 | 0.69[illegible] |
| 31 | 1.18 | GOTERM_BP_DIRECT | positive regulation of T cell activation | 5 | 409 | 45 | 19512 | 1.15[illegible] |
| 31 | 1.18 | GOTERM_BP_DIRECT | antigen processing and presentation | 5 | 409 | 49 | 19512 | 1.15[illegible] |
| 31 | 1.18 | GOTERM_MF_DIRECT | MHC class II protein complex binding | 4 | 417 | 27 | 19272 | 0.92[illegible] |
| 31 | 1.18 | GOTERM_CC_DIRECT | MHC class II protein complex | 3 | 415 | 26 | 20808 | 0.69[illegible] |
| 31 | 1.18 | GOTERM_BP_DIRECT | antigen processing and presentation of exogenous peptide antigen via MHC class II | 3 | 409 | 32 | 19512 | 0.69[illegible] |
| 31 | 1.18 | KEGG_PATHWAY | Th17 cell differentiation | 6 | 253 | 109 | 9496 | 1.38[illegible] |
| 31 | 1.18 | GOTERM_CC_DIRECT | clathrin-coated endocytic vesicle membrane | 4 | 415 | 71 | 20808 | 0.92[illegible] |
| 31 | 1.18 | KEGG_PATHWAY | Antigen processing and presentation | 5 | 253 | 82 | 9496 | 1.15[illegible] |
| 32 | 1.17 | INTERPRO | HD/PDEase_dom | 4 | 427 | 24 | 20827 | 0.92[illegible] |
| 32 | 1.17 | GOTERM_BP_DIRECT | negative regulation of cAMP/ PKA signal transduction | 4 | 409 | 28 | 19512 | 0.92[illegible] |
| 32 | 1.17 | UP_KW_LIGAND | cAMP | 4 | 165 | 37 | 7011 | 0.92[illegible] |
| 32 | 1.17 | INTERPRO | PDEase | 3 | 427 | 19 | 20827 | 0.69[illegible] |
| 32 | 1.17 | UP_SEQ_FEATURE | DOMAIN:PDEase | 3 | 422 | 21 | 20669 | 0.69[illegible] |
| 32 | 1.17 | INTERPRO | PDEase_catalytic_dom | 3 | 427 | 21 | 20827 | 0.69[illegible] |
| 32 | 1.17 | INTERPRO | PDEase_CS | 3 | 427 | 21 | 20827 | 0.69[illegible] |

| Cluster | Cluster Enrichment Score | Category | Term | Count | List Total | Pop Hits | Pop Total | % |
|---|---|---|---|---|---|---|---|---|
| 32 | 1.17 | INTERPRO | PDEase_catalytic_dom_sf | 3 | 427 | 21 | 20827 | 0.69 |
| 32 | 1.17 | GOTERM_MF_DIRECT | 3',5'-cyclic-AMP phosphodiesterase activity | 3 | 417 | 22 | 19272 | 0.69 |
| 32 | 1.17 | GOTERM_MF_DIRECT | 3',5'-cyclic-GMP phosphodiesterase activity | 3 | 417 | 23 | 19272 | 0.69 |
| 32 | 1.17 | GOTERM_MF_DIRECT | 3',5'-cyclic-nucleotide phosphodiesterase activity | 3 | 417 | 23 | 19272 | 0.69 |
| 32 | 1.17 | KEGG_PATHWAY | Morphine addiction | 6 | 253 | 91 | 9496 | 1.38 |
| 32 | 1.17 | GOTERM_MF_DIRECT | cAMP binding | 3 | 417 | 25 | 19272 | 0.69 |
| 32 | 1.17 | GOTERM_MF_DIRECT | phosphoric diester hydrolase activity | 4 | 417 | 58 | 19272 | 0.92 |
| 32 | 1.17 | KEGG_PATHWAY | Purine metabolism | 6 | 253 | 128 | 9496 | 1.38 |
| 33 | 1.16 | UP_SEQ_FEATURE | DOMAIN:Caspase family p10 | 3 | 422 | 11 | 20669 | 0.69 |
| 33 | 1.16 | INTERPRO | Caspase_his_AS | 3 | 427 | 11 | 20827 | 0.69 |
| 33 | 1.16 | UP_SEQ_FEATURE | DOMAIN:Caspase family p20 | 3 | 422 | 13 | 20669 | 0.69 |
| 33 | 1.16 | INTERPRO | Caspase_cys_AS | 3 | 427 | 13 | 20827 | 0.69 |
| 33 | 1.16 | INTERPRO | Pept_C14_p10 | 3 | 427 | 13 | 20827 | 0.69 |
| 33 | 1.16 | INTERPRO | Pept_C14A | 3 | 427 | 14 | 20827 | 0.69 |
| 33 | 1.16 | INTERPRO | Pept_C14_caspase | 3 | 427 | 15 | 20827 | 0.69 |
| 33 | 1.16 | INTERPRO | Pept_C14_p20 | 3 | 427 | 15 | 20827 | 0.69 |
| 33 | 1.16 | INTERPRO | Caspase-like_dom_sf | 3 | 427 | 15 | 20827 | 0.69 |
| 33 | 1.16 | SMART | CASc | 3 | 246 | 14 | 10704 | 0.69 |
| 33 | 1.16 | INTERPRO | DEATH-like_dom_sf | 5 | 427 | 105 | 20827 | 1.15 |
| 33 | 1.16 | GOTERM_MF_DIRECT | cysteine-type endopeptidase activity | 4 | 417 | 66 | 19272 | 0.92 |
| 33 | 1.16 | GOTERM_BP_DIRECT | regulation of apoptotic process | 8 | 409 | 291 | 19512 | 1.83 |
| 33 | 1.16 | UP_KW_MOLECULAR_FUNCTION | Thiol protease | 4 | 284 | 178 | 12059 | 0.92 |
| 33 | 1.16 | GOTERM_MF_DIRECT | cysteine-type peptidase activity | 3 | 417 | 155 | 19272 | 0.69 |
| 34 | 1.15 | GOTERM_BP_DIRECT | monocyte chemotaxis | 5 | 409 | 25 | 19512 | 1.15 |
| 34 | 1.15 | KEGG_PATHWAY | Viral protein interaction with cytokine and cytokine receptor | 7 | 253 | 100 | 9496 | 1.61 |
| 34 | 1.15 | GOTERM_BP_DIRECT | positive regulation of cytosolic calcium ion concentration | 7 | 409 | 139 | 19512 | 1.61 |

| Cluster | Cluster Enrichment Score | Category | Term | Count | List Total | Pop Hits | Pop Total | % |
|---|---|---|---|---|---|---|---|---|
| 34 | 1.15 | GOTERM_MF_DIRECT | C-C chemokine receptor activity | 3 | 417 | 23 | 19272 | 0.69 |
| 34 | 1.15 | GOTERM_MF_DIRECT | C-C chemokine binding | 3 | 417 | 23 | 19272 | 0.69 |
| 34 | 1.15 | GOTERM_BP_DIRECT | intracellular calcium ion homeostasis | 6 | 409 | 115 | 19512 | 1.38 |
| 34 | 1.15 | GOTERM_BP_DIRECT | cell chemotaxis | 5 | 409 | 87 | 19512 | 1.15 |
| 34 | 1.15 | INTERPRO | CCR1-9-like | 3 | 427 | 30 | 20827 | 0.69 |
| 34 | 1.15 | GOTERM_BP_DIRECT | chemokine-mediated signaling pathway | 4 | 409 | 64 | 19512 | 0.92 |
| 34 | 1.15 | KEGG_PATHWAY | Cytokine-cytokine receptor interaction | 10 | 253 | 298 | 9496 | 2.29 |
| 35 | 1.12 | GOTERM_BP_DIRECT | circadian regulation of gene expression | 6 | 409 | 68 | 19512 | 1.38 |
| 35 | 1.12 | GOTERM_BP_DIRECT | rhythmic process | 7 | 409 | 151 | 19512 | 1.61 |
| 35 | 1.12 | UP_KW_BIOLOGICAL_PROCESS | Biological rhythms | 7 | 278 | 149 | 11605 | 1.61 |
| 35 | 1.12 | GOTERM_BP_DIRECT | regulation of circadian rhythm | 4 | 409 | 69 | 19512 | 0.92 |
| 36 | 1.11 | GOTERM_MF_DIRECT | phosphatidylinositol-3,4-bisphosphate binding | 4 | 417 | 30 | 19272 | 0.92 |
| 36 | 1.11 | GOTERM_MF_DIRECT | phosphatidylinositol-3,4,5-trisphosphate binding | 4 | 417 | 42 | 19272 | 0.92 |
| 36 | 1.11 | GOTERM_MF_DIRECT | phosphatidylinositol-4,5-bisphosphate binding | 4 | 417 | 86 | 19272 | 0.92 |
| 37 | 1.08 | GOTERM_BP_DIRECT | positive regulation of T cell proliferation | 5 | 409 | 67 | 19512 | 1.15 |
| 37 | 1.08 | GOTERM_BP_DIRECT | symbiont entry into host cell | 6 | 409 | 112 | 19512 | 1.38 |
| 37 | 1.08 | UP_KW_MOLECULAR_FUNCTION | Host cell receptor for virus entry | 5 | 284 | 75 | 12059 | 1.15 |
| 37 | 1.08 | GOTERM_MF_DIRECT | virus receptor activity | 5 | 417 | 83 | 19272 | 1.15 |
| 38 | 1.01 | GOTERM_BP_DIRECT | angiogenesis | 16 | 409 | 292 | 19512 | 3.67 |
| 38 | 1.01 | UP_KW_BIOLOGICAL_PROCESS | Angiogenesis | 11 | 278 | 145 | 11605 | 2.52 |
| 38 | 1.01 | GOTERM_BP_DIRECT | cell differentiation | 24 | 409 | 1094 | 19512 | 5.50 |
| 38 | 1.01 | UP_KW_MOLECULAR_FUNCTION | Developmental protein | 23 | 284 | 1046 | 12059 | 5.28 |
| 38 | 1.01 | UP_KW_MOLECULAR_FUNCTION | Developmental protein | 23 | 284 | 1046 | 12059 | 5.28 |
| 38 | 1.01 | UP_KW_BIOLOGICAL_PROCESS | Differentiation | 15 | 278 | 825 | 11605 | 3.44 |
| 39 | 1.01 | INTERPRO | Collagen | 7 | 427 | 86 | 20827 | 1.61 |

| Cluster | Cluster Enrichment Score | Category | Term | Count | List Total | Pop Hits | Pop Total | % |
|---|---|---|---|---|---|---|---|---|
| 39 | 1.01 | UP_SEQ_FEATURE | REGION:Triple-helical region | 3 | 422 | 23 | 20669 | 0.69 |
| 39 | 1.01 | INTERPRO | Collagen_superfamily | 3 | 427 | 38 | 20827 | 0.69 |
| 39 | 1.01 | GOTERM_MF_DIRECT | extracellular matrix structural constituent conferring tensile strength | 3 | 417 | 44 | 19272 | 0.69 |
| 39 | 1.01 | KEGG_PATHWAY | Protein digestion and absorption | 5 | 253 | 105 | 9496 | 1.15 |
| 40 | 1.01 | GOTERM_BP_DIRECT | response to other organism | 6 | 409 | 19 | 19512 | 1.38 |
| 40 | 1.01 | INTERPRO | TPR_2 | 3 | 427 | 16 | 20827 | 0.69 |
| 40 | 1.01 | UP_SEQ_FEATURE | REPEAT:TPR 8 | 4 | 422 | 54 | 20669 | 0.92 |
| 40 | 1.01 | UP_SEQ_FEATURE | REPEAT:TPR | 5 | 422 | 99 | 20669 | 1.15 |
| 40 | 1.01 | UP_SEQ_FEATURE | REPEAT:TPR 7 | 4 | 422 | 66 | 20669 | 0.92 |
| 40 | 1.01 | INTERPRO | TPR_rpt | 6 | 427 | 140 | 20827 | 1.38 |
| 40 | 1.01 | UP_SEQ_FEATURE | REPEAT:TPR 3 | 6 | 422 | 146 | 20669 | 1.38 |
| 40 | 1.01 | SMART | TPR | 6 | 246 | 132 | 10704 | 1.38 |
| 40 | 1.01 | INTERPRO | TPR-like_helical_dom_sf | 8 | 427 | 230 | 20827 | 1.83 |
| 40 | 1.01 | UP_SEQ_FEATURE | REPEAT:TPR 6 | 4 | 422 | 77 | 20669 | 0.92 |
| 40 | 1.01 | UP_SEQ_FEATURE | REPEAT:TPR 1 | 6 | 422 | 160 | 20669 | 1.38 |
| 40 | 1.01 | UP_SEQ_FEATURE | REPEAT:TPR 2 | 6 | 422 | 160 | 20669 | 1.38 |
| 40 | 1.01 | UP_SEQ_FEATURE | REPEAT:TPR 5 | 4 | 422 | 82 | 20669 | 0.92 |
| 40 | 1.01 | UP_KW_DOMAIN | TPR repeat | 6 | 313 | 171 | 14656 | 1.38 |
| 40 | 1.01 | UP_SEQ_FEATURE | REPEAT:TPR 4 | 4 | 422 | 101 | 20669 | 0.92 |
| 41 | 0.96 | UP_SEQ_FEATURE | DOMAIN:Fibronectin type-III 8 | 4 | 422 | 23 | 20669 | 0.92 |
| 41 | 0.96 | UP_SEQ_FEATURE | DOMAIN:Fibronectin type-III 7 | 4 | 422 | 24 | 20669 | 0.92 |
| 41 | 0.96 | UP_SEQ_FEATURE | DOMAIN:Fibronectin type-III 5 | 5 | 422 | 45 | 20669 | 1.15 |
| 41 | 0.96 | UP_SEQ_FEATURE | DOMAIN:Fibronectin type-III 6 | 4 | 422 | 31 | 20669 | 0.92 |
| 41 | 0.96 | UP_SEQ_FEATURE | DOMAIN:Fibronectin type-III 4 | 5 | 422 | 64 | 20669 | 1.15 |
| 41 | 0.96 | UP_SEQ_FEATURE | DOMAIN:Fibronectin type-III 3 | 5 | 422 | 86 | 20669 | 1.15 |
| 41 | 0.96 | UP_SEQ_FEATURE | DOMAIN:Fibronectin type-III | 6 | 422 | 174 | 20669 | 1.38 |
| 41 | 0.96 | UP_SEQ_FEATURE | DOMAIN:Fibronectin type-III 1 | 5 | 422 | 139 | 20669 | 1.15 |
| 41 | 0.96 | UP_SEQ_FEATURE | DOMAIN:Fibronectin type-III 2 | 5 | 422 | 139 | 20669 | 1.15 |

| Cluster | Cluster Enrichment Score | Category | Term | Count | List Total | Pop Hits | Pop Total | % |
|---|---|---|---|---|---|---|---|---|
| 41 | 0.96 | INTERPRO | FN3_dom | 6 | 427 | 209 | 20827 | 1.38 |
| 41 | 0.96 | INTERPRO | FN3_sf | 6 | 427 | 211 | 20827 | 1.38 |
| 41 | 0.96 | SMART | FN3 | 5 | 246 | 150 | 10704 | 1.15 |
| 41 | 0.96 | GOTERM_CC_DIRECT | neuron projection | 6 | 415 | 372 | 20808 | 1.38 |
| 42 | 0.95 | KEGG_PATHWAY | Th1 and Th2 cell differentiation | 8 | 253 | 93 | 9496 | 1.83 |
| 42 | 0.95 | KEGG_PATHWAY | Yersinia infection | 9 | 253 | 138 | 9496 | 2.06 |
| 42 | 0.95 | KEGG_PATHWAY | Human T-cell leukemia virus 1 infection | 11 | 253 | 224 | 9496 | 2.52 |
| 42 | 0.95 | KEGG_PATHWAY | PD-L1 expression and PD-1 checkpoint pathway in cancer | 6 | 253 | 90 | 9496 | 1.38 |
| 42 | 0.95 | KEGG_PATHWAY | Th17 cell differentiation | 6 | 253 | 109 | 9496 | 1.38 |
| 42 | 0.95 | KEGG_PATHWAY | Human immunodeficiency virus 1 infection | 8 | 253 | 214 | 9496 | 1.83 |
| 42 | 0.95 | KEGG_PATHWAY | T cell receptor signaling pathway | 5 | 253 | 122 | 9496 | 1.15 |
| 42 | 0.95 | KEGG_PATHWAY | Hepatitis B | 6 | 253 | 163 | 9496 | 1.38 |
| 43 | 0.94 | UP_KW_LIGAND | FAD | 7 | 165 | 118 | 7011 | 1.61 |
| 43 | 0.94 | GOTERM_MF_DIRECT | FAD binding | 4 | 417 | 41 | 19272 | 0.92 |
| 43 | 0.94 | UP_KW_LIGAND | Flavoprotein | 7 | 165 | 130 | 7011 | 1.61 |
| 43 | 0.94 | INTERPRO | FAD/NAD-bd_sf | 4 | 427 | 62 | 20827 | 0.92 |
| 43 | 0.94 | UP_KW_BIOLOGICAL_PROCESS | Electron transport | 4 | 278 | 114 | 11605 | 0.92 |
| 44 | 0.94 | GOTERM_MF_DIRECT | transmembrane transporter activity | 10 | 417 | 231 | 19272 | 2.29 |
| 44 | 0.94 | UP_SEQ_FEATURE | TRANSMEM:Helical; Name=8 | 6 | 422 | 119 | 20669 | 1.38 |
| 44 | 0.94 | UP_SEQ_FEATURE | TRANSMEM:Helical; Name=12 | 5 | 422 | 89 | 20669 | 1.15 |
| 44 | 0.94 | UP_SEQ_FEATURE | TRANSMEM:Helical; Name=11 | 5 | 422 | 94 | 20669 | 1.15 |
| 44 | 0.94 | UP_SEQ_FEATURE | TRANSMEM:Helical; Name=10 | 5 | 422 | 101 | 20669 | 1.15 |
| 44 | 0.94 | UP_SEQ_FEATURE | TRANSMEM:Helical; Name=9 | 5 | 422 | 111 | 20669 | 1.15 |
| 45 | 0.90 | UP_SEQ_FEATURE | DOMAIN:Major facilitator superfamily (MFS) profile | 5 | 422 | 68 | 20669 | 1.15 |
| 45 | 0.90 | GOTERM_MF_DIRECT | transmembrane transporter activity | 10 | 417 | 231 | 19272 | 2.29 |
| 45 | 0.90 | INTERPRO | MFS_dom | 5 | 427 | 99 | 20827 | 1.15 |

| Cluster | Cluster Enrichment Score | Category | Term | Count | List Total | Pop Hits | Pop Total | % |
|---|---|---|---|---|---|---|---|---|
| 45 | 0.90 | INTERPRO | Sugar_transporter_CS | 3 | 427 | 33 | 20827 | 0.69 |
| 45 | 0.90 | INTERPRO | MFS_trans_sf | 6 | 427 | 147 | 20827 | 1.38 |
| 45 | 0.90 | INTERPRO | MFS_sugar_transport-like | 3 | 427 | 40 | 20827 | 0.69 |
| 45 | 0.90 | GOTERM_BP_DIRECT | transmembrane transport | 16 | 409 | 569 | 19512 | 3.67 |
| 46 | 0.89 | INTERPRO | EGF-like_dom | 11 | 427 | 254 | 20827 | 2.52 |
| 46 | 0.89 | UP_SEQ_FEATURE | DOMAIN:EGF-like 5 | 4 | 422 | 39 | 20669 | 0.92 |
| 46 | 0.89 | INTERPRO | EGF-like_Ca-bd_dom | 7 | 427 | 134 | 20827 | 1.61 |
| 46 | 0.89 | UP_KW_DOMAIN | EGF-like domain | 11 | 313 | 269 | 14656 | 2.52 |
| 46 | 0.89 | INTERPRO | Growth_fac_rcpt_cys_sf | 7 | 427 | 145 | 20827 | 1.61 |
| 46 | 0.89 | UP_SEQ_FEATURE | DOMAIN:EGF-like 6; calcium-binding | 3 | 422 | 23 | 20669 | 0.69 |
| 46 | 0.89 | UP_SEQ_FEATURE | DOMAIN:EGF-like 8 | 3 | 422 | 23 | 20669 | 0.69 |
| 46 | 0.89 | SMART | EGF_CA | 7 | 246 | 134 | 10704 | 1.61 |
| 46 | 0.89 | UP_SEQ_FEATURE | DOMAIN:EGF-like 7 | 3 | 422 | 26 | 20669 | 0.69 |
| 46 | 0.89 | UP_SEQ_FEATURE | DOMAIN:EGF-like 2; calcium-binding | 4 | 422 | 57 | 20669 | 0.92 |
| 46 | 0.89 | UP_SEQ_FEATURE | DOMAIN:EGF-like 1 | 6 | 422 | 129 | 20669 | 1.38 |
| 46 | 0.89 | INTERPRO | cEGF | 3 | 427 | 30 | 20827 | 0.69 |
| 46 | 0.89 | UP_SEQ_FEATURE | DOMAIN:EGF-like 4 | 4 | 422 | 66 | 20669 | 0.92 |
| 46 | 0.89 | UP_SEQ_FEATURE | DOMAIN:EGF-like | 8 | 422 | 221 | 20669 | 1.83 |
| 46 | 0.89 | SMART | EGF | 8 | 246 | 200 | 10704 | 1.83 |
| 46 | 0.89 | INTERPRO | EGF-type_Asp/ Asn_hydroxyl_site | 5 | 427 | 107 | 20827 | 1.15 |
| 46 | 0.89 | UP_SEQ_FEATURE | DOMAIN:EGF-like 3; calcium-binding | 3 | 422 | 38 | 20669 | 0.69 |
| 46 | 0.89 | UP_SEQ_FEATURE | DOMAIN:EGF-like 3 | 4 | 422 | 82 | 20669 | 0.92 |
| 46 | 0.89 | UP_SEQ_FEATURE | DOMAIN:EGF-like 6 | 3 | 422 | 53 | 20669 | 0.69 |
| 46 | 0.89 | UP_SEQ_FEATURE | DOMAIN:EGF-like 2 | 4 | 422 | 99 | 20669 | 0.92 |
| 46 | 0.89 | INTERPRO | EGF_Ca-bd_CS | 4 | 427 | 109 | 20827 | 0.92 |
| 46 | 0.89 | INTERPRO | NOTCH1_EGF-like | 3 | 427 | 80 | 20827 | 0.69 |
| 47 | 0.89 | GOTERM_MF_DIRECT | non-membrane spanning protein tyrosine phosphatase | 3 | 417 | 13 | 19272 | 0.69 |

| Cluster | Cluster Enrichment Score | Category | Term | Count | List Total | Pop Hits | Pop Total | % |
|---|---|---|---|---|---|---|---|---|
| | | | activity | | | | | |
| 47 | 0.89 | INTERPRO | PTP_cat | 4 | 427 | 38 | 20827 | 0.92 |
| 47 | 0.89 | SMART | PTPc | 4 | 246 | 37 | 10704 | 0.92 |
| 47 | 0.89 | UP_SEQ_FEATURE | DOMAIN:Tyrosine specific protein phosphatases | 5 | 422 | 69 | 20669 | 1.15 |
| 47 | 0.89 | UP_SEQ_FEATURE | DOMAIN:Tyrosine-protein phosphatase | 5 | 422 | 78 | 20669 | 1.15 |
| 47 | 0.89 | INTERPRO | Tyr_Pase_dom | 5 | 427 | 90 | 20827 | 1.15 |
| 47 | 0.89 | UP_SEQ_FEATURE | ACT_SITE:Phosphocysteine intermediate | 5 | 422 | 91 | 20669 | 1.15 |
| 47 | 0.89 | INTERPRO | Tyr_Pase_cat | 4 | 427 | 63 | 20827 | 0.92 |
| 47 | 0.89 | INTERPRO | Prot-tyrosine_phosphatase-like | 5 | 427 | 105 | 20827 | 1.15 |
| 47 | 0.89 | GOTERM_MF_DIRECT | protein tyrosine phosphatase activity | 5 | 417 | 101 | 19272 | 1.15 |
| 47 | 0.89 | SMART | PTPc_motif | 4 | 246 | 63 | 10704 | 0.92 |
| 47 | 0.89 | INTERPRO | Tyr_Pase_AS | 4 | 427 | 77 | 20827 | 0.92 |
| 47 | 0.89 | GOTERM_MF_DIRECT | phosphoprotein phosphatase activity | 5 | 417 | 153 | 19272 | 1.15 |
| 47 | 0.89 | UP_KW_MOLECULAR_FUNCTION | Protein phosphatase | 5 | 284 | 144 | 12059 | 1.15 |
| 47 | 0.89 | GOTERM_MF_DIRECT | phosphatase activity | 3 | 417 | 82 | 19272 | 0.69 |
| 48 | 0.86 | UP_SEQ_FEATURE | DOMAIN:bHLH | 6 | 422 | 114 | 20669 | 1.38 |
| 48 | 0.86 | INTERPRO | HLH_DNA-bd_sf | 6 | 427 | 116 | 20827 | 1.38 |
| 48 | 0.86 | INTERPRO | bHLH_dom | 6 | 427 | 116 | 20827 | 1.38 |
| 48 | 0.86 | SMART | HLH | 6 | 246 | 114 | 10704 | 1.38 |
| 48 | 0.86 | UP_SEQ_FEATURE | DOMAIN:BHLH | 4 | 422 | 75 | 20669 | 0.92 |
| 48 | 0.86 | GOTERM_MF_DIRECT | protein dimerization activity | 6 | 417 | 199 | 19272 | 1.38 |
| 49 | 0.86 | GOTERM_BP_DIRECT | positive regulation of DNA-templated transcription | 33 | 409 | 791 | 19512 | 7.57 |
| 49 | 0.86 | GOTERM_CC_DIRECT | transcription regulator complex | 15 | 415 | 251 | 20808 | 3.44 |
| 49 | 0.86 | GOTERM_MF_DIRECT | DNA-binding transcription factor activity | 26 | 417 | 739 | 19272 | 5.96 |
| 49 | 0.86 | UP_KW_MOLECULAR_FUNCTION | Activator | 26 | 284 | 733 | 12059 | 5.96 |
| 49 | 0.86 | GOTERM_MF_DIRECT | DNA-binding transcription | 17 | 417 | 456 | 19272 | 3.90 |

| Cluster | Cluster Enrichment Score | Category | Term | Count | List Total | Pop Hits | Pop Total | % |
|---|---|---|---|---|---|---|---|---|
| | | | activator activity, RNA polymerase II-specific | | | | | |
| 49 | 0.86 | GOTERM_MF_DIRECT | transcription cis-regulatory region binding | 11 | 417 | 286 | 19272 | 2.52 |
| 49 | 0.86 | GOTERM_BP_DIRECT | positive regulation of transcription by RNA polymerase II | 33 | 409 | 1243 | 19512 | 7.57 |
| 49 | 0.86 | GOTERM_CC_DIRECT | chromatin | 29 | 415 | 1159 | 20808 | 6.65 |
| 49 | 0.86 | GOTERM_MF_DIRECT | sequence-specific DNA binding | 15 | 417 | 514 | 19272 | 3.44 |
| 49 | 0.86 | GOTERM_BP_DIRECT | regulation of DNA-templated transcription | 37 | 409 | 1526 | 19512 | 8.49 |
| 49 | 0.86 | GOTERM_BP_DIRECT | negative regulation of transcription by RNA polymerase II | 26 | 409 | 1034 | 19512 | 5.96 |
| 49 | 0.86 | GOTERM_MF_DIRECT | RNA polymerase II cis-regulatory region sequence-specific DNA binding | 27 | 417 | 1100 | 19272 | 6.19 |
| 49 | 0.86 | GOTERM_MF_DIRECT | DNA-binding transcription factor activity, RNA polymerase II-specific | 29 | 417 | 1247 | 19272 | 6.65 |
| 49 | 0.86 | GOTERM_MF_DIRECT | chromatin binding | 13 | 417 | 524 | 19272 | 2.98 |
| 49 | 0.86 | GOTERM_BP_DIRECT | regulation of transcription by RNA polymerase II | 31 | 409 | 1643 | 19512 | 7.11 |
| 49 | 0.86 | UP_KW_BIOLOGICAL_PROCESS | Transcription regulation | 46 | 278 | 2444 | 11605 | 10.5 |
| 49 | 0.86 | GOTERM_MF_DIRECT | DNA binding | 38 | 417 | 2370 | 19272 | 8.72 |
| 49 | 0.86 | UP_KW_BIOLOGICAL_PROCESS | Transcription | 46 | 278 | 2516 | 11605 | 10.5 |
| 49 | 0.86 | UP_KW_MOLECULAR_FUNCTION | DNA-binding | 35 | 284 | 2143 | 12059 | 8.03 |
| 49 | 0.86 | UP_KW_CELLULAR_COMPONENT | Nucleus | 87 | 387 | 5969 | 18195 | 19.9 |
| 50 | 0.85 | GOTERM_BP_DIRECT | lipoprotein metabolic process | 4 | 409 | 27 | 19512 | 0.92 |
| 50 | 0.85 | GOTERM_BP_DIRECT | lipid transport | 6 | 409 | 193 | 19512 | 1.38 |
| 50 | 0.85 | UP_KW_BIOLOGICAL_PROCESS | Lipid transport | 6 | 278 | 177 | 11605 | 1.38 |
| 51 | 0.80 | KEGG_PATHWAY | Yersinia infection | 9 | 253 | 138 | 9496 | 2.06 |
| 51 | 0.80 | KEGG_PATHWAY | Pathogenic Escherichia coli infection | 9 | 253 | 203 | 9496 | 2.06 |
| 51 | 0.80 | KEGG_PATHWAY | Salmonella infection | 9 | 253 | 251 | 9496 | 2.06 |

| Cluster | Cluster Enrichment Score | Category | Term | Count | List Total | Pop Hits | Pop Total | % |
|---|---|---|---|---|---|---|---|---|
| 51 | 0.80 | KEGG_PATHWAY | Shigellosis | 9 | 253 | 253 | 9496 | 2.06 |
| 52 | 0.77 | UP_KW_MOLECULAR_FUNCTION | Antimicrobial | 7 | 284 | 116 | 12059 | 1.61 |
| 52 | 0.77 | GOTERM_BP_DIRECT | antibacterial humoral response | 4 | 409 | 75 | 19512 | 0.92 |
| 52 | 0.77 | UP_KW_MOLECULAR_FUNCTION | Antibiotic | 4 | 284 | 100 | 12059 | 0.92 |
| 53 | 0.77 | GOTERM_BP_DIRECT | fatty acid metabolic process | 11 | 409 | 214 | 19512 | 2.52 |
| 53 | 0.77 | UP_KW_BIOLOGICAL_PROCESS | Fatty acid metabolism | 7 | 278 | 167 | 11605 | 1.61 |
| 53 | 0.77 | GOTERM_BP_DIRECT | fatty acid biosynthetic process | 4 | 409 | 83 | 19512 | 0.92 |
| 53 | 0.77 | UP_KW_BIOLOGICAL_PROCESS | Fatty acid biosynthesis | 3 | 278 | 56 | 11605 | 0.69 |
| 53 | 0.77 | KEGG_PATHWAY | Fatty acid metabolism | 3 | 253 | 57 | 9496 | 0.69 |
| 54 | 0.77 | GOTERM_MF_DIRECT | extracellular matrix structural constituent | 6 | 417 | 127 | 19272 | 1.38 |
| 54 | 0.77 | GOTERM_CC_DIRECT | basement membrane | 5 | 415 | 102 | 20808 | 1.15 |
| 54 | 0.77 | UP_KW_CELLULAR_COMPONENT | Basement membrane | 3 | 387 | 44 | 18195 | 0.69 |
| 55 | 0.75 | INTERPRO | SIGLEC | 3 | 427 | 13 | 20827 | 0.69 |
| 55 | 0.75 | GOTERM_MF_DIRECT | sialic acid binding | 3 | 417 | 22 | 19272 | 0.69 |
| 55 | 0.75 | INTERPRO | C-type_lectin-like/link_sf | 5 | 427 | 106 | 20827 | 1.15 |
| 55 | 0.75 | INTERPRO | CTDL_fold | 5 | 427 | 114 | 20827 | 1.15 |
| 55 | 0.75 | GOTERM_MF_DIRECT | carbohydrate binding | 8 | 417 | 231 | 19272 | 1.83 |
| 55 | 0.75 | INTERPRO | C-type_lectin_CS | 3 | 427 | 46 | 20827 | 0.69 |
| 55 | 0.75 | UP_KW_LIGAND | Lectin | 7 | 165 | 182 | 7011 | 1.61 |
| 55 | 0.75 | UP_SEQ_FEATURE | DOMAIN:C-type lectin | 4 | 422 | 89 | 20669 | 0.92 |
| 55 | 0.75 | INTERPRO | C-type_lectin-like | 4 | 427 | 90 | 20827 | 0.92 |
| 55 | 0.75 | SMART | CLECT | 4 | 246 | 88 | 10704 | 0.92 |
| 56 | 0.73 | UP_KW_DOMAIN | Laminin EGF-like domain | 3 | 313 | 32 | 14656 | 0.69 |
| 56 | 0.73 | SMART | EGF_Lam | 3 | 246 | 36 | 10704 | 0.69 |
| 56 | 0.73 | INTERPRO | LE_dom | 3 | 427 | 42 | 20827 | 0.69 |
| 57 | 0.72 | INTERPRO | Peptidase_M12B_N | 3 | 427 | 35 | 20827 | 0.69 |
| 57 | 0.72 | UP_KW_MOLECULAR_FUNCTION | Metalloprotease | 7 | 284 | 156 | 12059 | 1.61 |
| 57 | 0.72 | GOTERM_MF_DIRECT | metallopeptidase activity | 7 | 417 | 171 | 19272 | 1.61 |
| 57 | 0.72 | UP_SEQ_FEATURE | DOMAIN:Peptidase M12B | 3 | 422 | 40 | 20669 | 0.69 |

| Cluster | Cluster Enrichment Score | Category | Term | Count | List Total | Pop Hits | Pop Total | % |
|---|---|---|---|---|---|---|---|---|
| 57 | 0.72 | INTERPRO | Peptidase_M12B | 3 | 427 | 40 | 20827 | 0.69 |
| 57 | 0.72 | GOTERM_MF_DIRECT | metalloendopeptidase activity | 5 | 417 | 109 | 19272 | 1.15 |
| 57 | 0.72 | UP_SEQ_FEATURE | DOMAIN:Disintegrin | 3 | 422 | 42 | 20669 | 0.69 |
| 57 | 0.72 | INTERPRO | MetalloPept_cat_dom_sf | 4 | 427 | 84 | 20827 | 0.92 |
| 58 | 0.69 | INTERPRO | PH-like_dom_sf | 14 | 427 | 452 | 20827 | 3.21 |
| 58 | 0.69 | INTERPRO | PH_domain | 9 | 427 | 276 | 20827 | 2.06 |
| 58 | 0.69 | UP_SEQ_FEATURE | DOMAIN:PH | 9 | 422 | 283 | 20669 | 2.06 |
| 58 | 0.69 | SMART | PH | 9 | 246 | 267 | 10704 | 2.06 |
| 59 | 0.69 | GOTERM_MF_DIRECT | phosphorus-oxygen lyase activity | 3 | 417 | 19 | 19272 | 0.69 |
| 59 | 0.69 | KEGG_PATHWAY | Salivary secretion | 5 | 253 | 97 | 9496 | 1.15 |
| 59 | 0.69 | KEGG_PATHWAY | Pancreatic secretion | 4 | 253 | 106 | 9496 | 0.92 |
| 60 | 0.69 | UP_SEQ_FEATURE | DOMAIN:Immunoglobulin | 5 | 422 | 70 | 20669 | 1.15 |
| 60 | 0.69 | GOTERM_BP_DIRECT | immune response-regulating signaling pathway | 3 | 409 | 64 | 19512 | 0.69 |
| 60 | 0.69 | INTERPRO | Ig-like_Receptors_ImmuneReg | 3 | 427 | 69 | 20827 | 0.69 |
| 61 | 0.68 | GOTERM_CC_DIRECT | autophagosome | 5 | 415 | 86 | 20808 | 1.15 |
| 61 | 0.68 | GOTERM_BP_DIRECT | autophagosome assembly | 5 | 409 | 98 | 19512 | 1.15 |
| 61 | 0.68 | KEGG_PATHWAY | Autophagy - animal | 5 | 253 | 169 | 9496 | 1.15 |
| 62 | 0.68 | GOTERM_CC_DIRECT | lumenal side of endoplasmic reticulum membrane | 3 | 415 | 35 | 20808 | 0.69 |
| 62 | 0.68 | KEGG_PATHWAY | Antigen processing and presentation | 5 | 253 | 82 | 9496 | 1.15 |
| 62 | 0.68 | GOTERM_CC_DIRECT | ER to Golgi transport vesicle membrane | 3 | 415 | 62 | 20808 | 0.69 |
| 63 | 0.67 | UP_KW_CELLULAR_COMPONENT | Lipid droplet | 5 | 387 | 72 | 18195 | 1.15 |
| 63 | 0.67 | GOTERM_MF_DIRECT | acyltransferase activity | 6 | 417 | 181 | 19272 | 1.38 |
| 63 | 0.67 | UP_KW_MOLECULAR_FUNCTION | Acyltransferase | 6 | 284 | 183 | 12059 | 1.38 |
| 64 | 0.63 | GOTERM_BP_DIRECT | blood coagulation | 6 | 409 | 111 | 19512 | 1.38 |
| 64 | 0.63 | GOTERM_BP_DIRECT | hemostasis | 3 | 409 | 55 | 19512 | 0.69 |
| 64 | 0.63 | UP_KW_BIOLOGICAL_PROCESS | Blood coagulation | 3 | 278 | 50 | 11605 | 0.69 |
| 64 | 0.63 | UP_KW_BIOLOGICAL_PROCESS | Hemostasis | 3 | 278 | 50 | 11605 | 0.69 |

| Cluster | Cluster Enrichment Score | Category | Term | Count | List Total | Pop Hits | Pop Total | % |
|---|---|---|---|---|---|---|---|---|
| 65 | 0.59 | INTERPRO | VWF_A | 4 | 427 | 79 | 20827 | 0.92 |
| 65 | 0.59 | SMART | VWA | 4 | 246 | 74 | 10704 | 0.92 |
| 65 | 0.59 | UP_SEQ_FEATURE | DOMAIN:VWFA | 4 | 422 | 85 | 20669 | 0.92 |
| 65 | 0.59 | INTERPRO | vWFA_dom_sf | 4 | 427 | 102 | 20827 | 0.92 |
| 66 | 0.56 | GOTERM_MF_DIRECT | actin filament binding | 8 | 417 | 213 | 19272 | 1.83 |
| 66 | 0.56 | GOTERM_MF_DIRECT | actin binding | 12 | 417 | 412 | 19272 | 2.75 |
| 66 | 0.56 | UP_KW_MOLECULAR_FUNCTION | Actin-binding | 9 | 284 | 300 | 12059 | 2.06 |
| 67 | 0.51 | GOTERM_BP_DIRECT | calcium ion transmembrane transport | 7 | 409 | 150 | 19512 | 1.61 |
| 67 | 0.51 | GOTERM_BP_DIRECT | calcium ion transport | 7 | 409 | 158 | 19512 | 1.61 |
| 67 | 0.51 | UP_KW_BIOLOGICAL_PROCESS | Calcium transport | 6 | 278 | 109 | 11605 | 1.38 |
| 67 | 0.51 | GOTERM_BP_DIRECT | monoatomic ion transport | 19 | 409 | 672 | 19512 | 4.36 |
| 67 | 0.51 | UP_KW_MOLECULAR_FUNCTION | Calcium channel | 4 | 284 | 75 | 12059 | 0.92 |
| 67 | 0.51 | GOTERM_MF_DIRECT | monoatomic cation channel activity | 3 | 417 | 51 | 19272 | 0.69 |
| 67 | 0.51 | GOTERM_CC_DIRECT | monoatomic ion channel complex | 5 | 415 | 150 | 20808 | 1.15 |
| 67 | 0.51 | UP_KW_BIOLOGICAL_PROCESS | Ion transport | 18 | 278 | 647 | 11605 | 4.13 |
| 67 | 0.51 | UP_KW_MOLECULAR_FUNCTION | Voltage-gated channel | 5 | 284 | 141 | 12059 | 1.15 |
| 67 | 0.51 | GOTERM_MF_DIRECT | calcium channel activity | 4 | 417 | 111 | 19272 | 0.92 |
| 67 | 0.51 | UP_KW_MOLECULAR_FUNCTION | Ion channel | 10 | 284 | 356 | 12059 | 2.29 |
| 67 | 0.51 | GOTERM_BP_DIRECT | monoatomic ion transmembrane transport | 10 | 409 | 413 | 19512 | 2.29 |
| 67 | 0.51 | GOTERM_MF_DIRECT | monoatomic ion channel activity | 4 | 417 | 189 | 19272 | 0.92 |
| 67 | 0.51 | UP_KW_BIOLOGICAL_PROCESS | Transport | 46 | 278 | 2133 | 11605 | 10.5 |
| 68 | 0.47 | GOTERM_MF_DIRECT | GTPase activity | 13 | 417 | 364 | 19272 | 2.98 |
| 68 | 0.47 | UP_KW_LIGAND | GTP-binding | 13 | 165 | 375 | 7011 | 2.98 |
| 68 | 0.47 | GOTERM_MF_DIRECT | GTP binding | 13 | 417 | 407 | 19272 | 2.98 |
| 68 | 0.47 | INTERPRO | Small_GTPase | 5 | 427 | 144 | 20827 | 1.15 |
| 68 | 0.47 | SMART | RHO | 5 | 246 | 132 | 10704 | 1.15 |
| 68 | 0.47 | UP_SEQ_FEATURE | LIPID:S-geranylgeranyl cysteine | 4 | 422 | 112 | 20669 | 0.92 |

| Cluster | Cluster Enrichment Score | Category | Term | Count | List Total | Pop Hits | Pop Total | % |
|---|---|---|---|---|---|---|---|---|
| 68 | 0.47 | UP_KW_PTM | Prenylation | 6 | 343 | 177 | 14452 | 1.38 |
| 68 | 0.47 | SMART | RAS | 5 | 246 | 143 | 10704 | 1.15 |
| 68 | 0.47 | INTERPRO | Small_GTP-bd | 5 | 427 | 176 | 20827 | 1.15 |
| 68 | 0.47 | SMART | RAB | 5 | 246 | 159 | 10704 | 1.15 |
| 68 | 0.47 | INTERPRO | P-loop_NTPase | 19 | 427 | 917 | 20827 | 4.36 |
| 68 | 0.47 | SMART | RAN | 3 | 246 | 91 | 10704 | 0.69 |
| 69 | 0.47 | UP_SEQ_FEATURE | DOMAIN:PX | 3 | 422 | 51 | 20669 | 0.69 |
| 69 | 0.47 | INTERPRO | PX_dom_sf | 3 | 427 | 55 | 20827 | 0.69 |
| 69 | 0.47 | GOTERM_MF_DIRECT | phosphatidylinositol binding | 4 | 417 | 116 | 19272 | 0.92 |
| 70 | 0.45 | GOTERM_BP_DIRECT | protein autophosphorylation | 7 | 409 | 119 | 19512 | 1.61 |
| 70 | 0.45 | GOTERM_BP_DIRECT | peptidyl-tyrosine phosphorylation | 3 | 409 | 41 | 19512 | 0.69 |
| 70 | 0.45 | GOTERM_MF_DIRECT | non-membrane spanning protein tyrosine kinase activity | 3 | 417 | 46 | 19272 | 0.69 |
| 70 | 0.45 | GOTERM_BP_DIRECT | cell surface receptor protein tyrosine kinase signaling pathway | 5 | 409 | 125 | 19512 | 1.15 |
| 70 | 0.45 | INTERPRO | Tyr_kinase_AS | 4 | 427 | 98 | 20827 | 0.92 |
| 70 | 0.45 | INTERPRO | Tyr_kinase_cat_dom | 3 | 427 | 91 | 20827 | 0.69 |
| 70 | 0.45 | SMART | TyrKc | 3 | 246 | 91 | 10704 | 0.69 |
| 70 | 0.45 | UP_KW_MOLECULAR_FUNCTION | Tyrosine-protein kinase | 3 | 284 | 118 | 12059 | 0.69 |
| 70 | 0.45 | GOTERM_MF_DIRECT | protein tyrosine kinase activity | 3 | 417 | 140 | 19272 | 0.69 |
| 70 | 0.45 | INTERPRO | Ser-Thr/Tyr_kinase_cat_dom | 3 | 427 | 151 | 20827 | 0.69 |
| 71 | 0.45 | UP_SEQ_FEATURE | REPEAT:4 | 7 | 422 | 191 | 20669 | 1.61 |
| 71 | 0.45 | UP_SEQ_FEATURE | REPEAT:1 | 8 | 422 | 250 | 20669 | 1.83 |
| 71 | 0.45 | UP_SEQ_FEATURE | REPEAT:2 | 8 | 422 | 253 | 20669 | 1.83 |
| 71 | 0.45 | UP_SEQ_FEATURE | REPEAT:7 | 5 | 422 | 137 | 20669 | 1.15 |
| 71 | 0.45 | UP_SEQ_FEATURE | REPEAT:3 | 7 | 422 | 225 | 20669 | 1.61 |
| 71 | 0.45 | UP_SEQ_FEATURE | REPEAT:6 | 5 | 422 | 150 | 20669 | 1.15 |
| 71 | 0.45 | UP_SEQ_FEATURE | REPEAT:5 | 5 | 422 | 167 | 20669 | 1.15 |
| 71 | 0.45 | UP_SEQ_FEATURE | REPEAT:9 | 3 | 422 | 111 | 20669 | 0.69 |

| Cluster | Cluster Enrichment Score | Category | Term | Count | List Total | Pop Hits | Pop Total | % |
|---|---|---|---|---|---|---|---|---|
| 71 | 0.45 | UP_SEQ_FEATURE | REPEAT:8 | 3 | 422 | 126 | 20669 | 0.69 |
| 72 | 0.44 | GOTERM_MF_DIRECT | iron ion binding | 6 | 417 | 148 | 19272 | 1.38 |
| 72 | 0.44 | GOTERM_MF_DIRECT | monooxygenase activity | 4 | 417 | 96 | 19272 | 0.92 |
| 72 | 0.44 | COG_ONTOLOGY | Secondary metabolites biosynthesis, transport, and catabolism | 3 | 40 | 66 | 2025 | 0.69 |
| 72 | 0.44 | UP_KW_MOLECULAR_FUNCTION | Monooxygenase | 4 | 284 | 92 | 12059 | 0.92 |
| 72 | 0.44 | INTERPRO | Cyt_P450 | 3 | 427 | 66 | 20827 | 0.69 |
| 72 | 0.44 | INTERPRO | Cyt_P450_sf | 3 | 427 | 66 | 20827 | 0.69 |
| 72 | 0.44 | GOTERM_MF_DIRECT | oxidoreductase activity, acting on paired donors, with incorporation or reduction of molecular oxygen | 3 | 417 | 82 | 19272 | 0.69 |
| 73 | 0.44 | UP_KW_DOMAIN | Sushi | 3 | 313 | 58 | 14656 | 0.69 |
| 73 | 0.44 | INTERPRO | Sushi_SCR_CCP_dom | 3 | 427 | 61 | 20827 | 0.69 |
| 73 | 0.44 | UP_SEQ_FEATURE | DOMAIN:Sushi | 3 | 422 | 62 | 20669 | 0.69 |
| 73 | 0.44 | INTERPRO | Sushi/SCR/CCP_sf | 3 | 427 | 62 | 20827 | 0.69 |
| 73 | 0.44 | SMART | CCP | 3 | 246 | 60 | 10704 | 0.69 |
| 74 | 0.43 | GOTERM_BP_DIRECT | chemotaxis | 12 | 409 | 163 | 19512 | 2.75 |
| 74 | 0.43 | GOTERM_BP_DIRECT | dendritic cell chemotaxis | 3 | 409 | 16 | 19512 | 0.69 |
| 74 | 0.43 | GOTERM_BP_DIRECT | positive regulation of cytosolic calcium ion concentration | 7 | 409 | 139 | 19512 | 1.61 |
| 74 | 0.43 | GOTERM_BP_DIRECT | cell chemotaxis | 5 | 409 | 87 | 19512 | 1.15 |
| 74 | 0.43 | SMART | 7TM_GPCR_Srsx | 4 | 246 | 124 | 10704 | 0.92 |
| 74 | 0.43 | UP_SEQ_FEATURE | TRANSMEM:Helical; Name=6 | 21 | 422 | 1046 | 20669 | 4.82 |
| 74 | 0.43 | UP_SEQ_FEATURE | TRANSMEM:Helical; Name=2 | 22 | 422 | 1106 | 20669 | 5.05 |
| 74 | 0.43 | UP_SEQ_FEATURE | TRANSMEM:Helical; Name=5 | 21 | 422 | 1056 | 20669 | 4.82 |
| 74 | 0.43 | UP_SEQ_FEATURE | TRANSMEM:Helical; Name=7 | 19 | 422 | 968 | 20669 | 4.36 |
| 74 | 0.43 | UP_SEQ_FEATURE | TRANSMEM:Helical; Name=3 | 21 | 422 | 1082 | 20669 | 4.82 |
| 74 | 0.43 | UP_SEQ_FEATURE | TRANSMEM:Helical; Name=4 | 21 | 422 | 1084 | 20669 | 4.82 |
| 74 | 0.43 | UP_SEQ_FEATURE | TRANSMEM:Helical; Name=1 | 21 | 422 | 1099 | 20669 | 4.82 |
| 74 | 0.43 | GOTERM_BP_DIRECT | G protein-coupled receptor signaling pathway | 19 | 409 | 1041 | 19512 | 4.36 |

| Cluster | Cluster Enrichment Score | Category | Term | Count | List Total | Pop Hits | Pop Total | % |
|---|---|---|---|---|---|---|---|---|
| 74 | 0.43 | UP_SEQ_FEATURE | DOMAIN:G-protein coupled receptors family 1 profile | 9 | 422 | 635 | 20669 | 2.06 |
| 74 | 0.43 | INTERPRO | GPCR_Rhodpsn | 10 | 427 | 739 | 20827 | 2.29 |
| 74 | 0.43 | INTERPRO | GPCR_Rhodpsn_7TM | 10 | 427 | 758 | 20827 | 2.29 |
| 74 | 0.43 | GOTERM_MF_DIRECT | G protein-coupled receptor activity | 11 | 417 | 869 | 19272 | 2.52 |
| 74 | 0.43 | UP_KW_MOLECULAR_FUNCTION | G-protein coupled receptor | 11 | 284 | 862 | 12059 | 2.52 |
| 74 | 0.43 | UP_KW_MOLECULAR_FUNCTION | Transducer | 12 | 284 | 927 | 12059 | 2.75 |
| 75 | 0.40 | UP_SEQ_FEATURE | PROPEP:Removed in mature form | 10 | 422 | 297 | 20669 | 2.29 |
| 75 | 0.40 | GOTERM_CC_DIRECT | side of membrane | 4 | 415 | 150 | 20808 | 0.92 |
| 75 | 0.40 | UP_KW_PTM | GPI-anchor | 4 | 343 | 151 | 14452 | 0.92 |
| 76 | 0.38 | INTERPRO | Cys-rich_flank_reg_C | 5 | 427 | 83 | 20827 | 1.15 |
| 76 | 0.38 | UP_SEQ_FEATURE | DOMAIN:LRRCT | 6 | 422 | 126 | 20669 | 1.38 |
| 76 | 0.38 | SMART | LRRCT | 5 | 246 | 83 | 10704 | 1.15 |
| 76 | 0.38 | INTERPRO | Leu-rich_rpt_typical-subtyp | 6 | 427 | 182 | 20827 | 1.38 |
| 76 | 0.38 | UP_SEQ_FEATURE | DOMAIN:LRRNT | 4 | 422 | 101 | 20669 | 0.92 |
| 76 | 0.38 | SMART | LRR_TYP | 6 | 246 | 182 | 10704 | 1.38 |
| 76 | 0.38 | INTERPRO | Leu-rich_rpt | 7 | 427 | 268 | 20827 | 1.61 |
| 76 | 0.38 | UP_SEQ_FEATURE | REPEAT:LRR 8 | 5 | 422 | 175 | 20669 | 1.15 |
| 76 | 0.38 | UP_SEQ_FEATURE | REPEAT:LRR 3 | 7 | 422 | 277 | 20669 | 1.61 |
| 76 | 0.38 | UP_SEQ_FEATURE | REPEAT:LRR 6 | 6 | 422 | 231 | 20669 | 1.38 |
| 76 | 0.38 | UP_SEQ_FEATURE | REPEAT:LRR 2 | 7 | 422 | 286 | 20669 | 1.61 |
| 76 | 0.38 | UP_SEQ_FEATURE | REPEAT:LRR 1 | 7 | 422 | 289 | 20669 | 1.61 |
| 76 | 0.38 | UP_SEQ_FEATURE | REPEAT:LRR 11 | 3 | 422 | 93 | 20669 | 0.69 |
| 76 | 0.38 | UP_SEQ_FEATURE | REPEAT:LRR 7 | 5 | 422 | 206 | 20669 | 1.15 |
| 76 | 0.38 | UP_SEQ_FEATURE | REPEAT:LRR 5 | 6 | 422 | 262 | 20669 | 1.38 |
| 76 | 0.38 | UP_SEQ_FEATURE | REPEAT:LRR 4 | 6 | 422 | 264 | 20669 | 1.38 |
| 76 | 0.38 | UP_SEQ_FEATURE | REPEAT:LRR 9 | 4 | 422 | 159 | 20669 | 0.92 |
| 76 | 0.38 | UP_SEQ_FEATURE | REPEAT:LRR 10 | 3 | 422 | 109 | 20669 | 0.69 |
| 76 | 0.38 | UP_KW_DOMAIN | Leucine-rich repeat | 7 | 313 | 321 | 14656 | 1.61 |

| Cluster | Cluster Enrichment Score | Category | Term | Count | List Total | Pop Hits | Pop Total | % |
|---|---|---|---|---|---|---|---|---|
| 76 | 0.38 | INTERPRO | LRR_dom_sf | 7 | 427 | 340 | 20827 | 1.61 |
| 77 | 0.35 | GOTERM_CC_DIRECT | stereocilium | 4 | 415 | 56 | 20808 | 0.92 |
| 77 | 0.35 | UP_KW_DISEASE | Non-syndromic deafness | 4 | 125 | 139 | 4888 | 0.92 |
| 77 | 0.35 | GOTERM_BP_DIRECT | sensory perception of sound | 4 | 409 | 171 | 19512 | 0.92 |
| 77 | 0.35 | UP_KW_DISEASE | Deafness | 7 | 125 | 305 | 4888 | 1.61 |
| 78 | 0.35 | GOTERM_BP_DIRECT | positive regulation of NF-kappaB transcription factor activity | 6 | 409 | 94 | 19512 | 1.38 |
| 78 | 0.35 | GOTERM_BP_DIRECT | positive regulation of DNA-binding transcription factor activity | 3 | 409 | 46 | 19512 | 0.69 |
| 78 | 0.35 | INTERPRO | Butyrophylin_SPRY | 3 | 427 | 76 | 20827 | 0.69 |
| 78 | 0.35 | INTERPRO | SPRY_dom | 3 | 427 | 98 | 20827 | 0.69 |
| 78 | 0.35 | UP_SEQ_FEATURE | DOMAIN:B30.2/SPRY | 3 | 422 | 102 | 20669 | 0.69 |
| 78 | 0.35 | INTERPRO | B30.2/SPRY | 3 | 427 | 102 | 20827 | 0.69 |
| 78 | 0.35 | SMART | SPRY | 3 | 246 | 93 | 10704 | 0.69 |
| 78 | 0.35 | INTERPRO | B30.2/SPRY_sf | 3 | 427 | 112 | 20827 | 0.69 |
| 78 | 0.35 | INTERPRO | ConA-like_dom_sf | 5 | 427 | 237 | 20827 | 1.15 |
| 78 | 0.35 | GOTERM_MF_DIRECT | ubiquitin protein ligase activity | 6 | 417 | 383 | 19272 | 1.38 |
| 79 | 0.33 | UP_SEQ_FEATURE | DOMAIN:EF-hand 1 | 6 | 422 | 188 | 20669 | 1.38 |
| 79 | 0.33 | INTERPRO | EF_hand_dom | 7 | 427 | 233 | 20827 | 1.61 |
| 79 | 0.33 | UP_SEQ_FEATURE | DOMAIN:EF-hand | 6 | 422 | 199 | 20669 | 1.38 |
| 79 | 0.33 | INTERPRO | EF_Hand_1_Ca_BS | 5 | 427 | 181 | 20827 | 1.15 |
| 79 | 0.33 | INTERPRO | EF-hand-dom_pair | 7 | 427 | 276 | 20827 | 1.61 |
| 79 | 0.33 | UP_SEQ_FEATURE | DOMAIN:EF-hand 2 | 5 | 422 | 190 | 20669 | 1.15 |
| 79 | 0.33 | GOTERM_MF_DIRECT | calcium ion binding | 17 | 417 | 744 | 19272 | 3.90 |
| 79 | 0.33 | SMART | EFh | 4 | 246 | 149 | 10704 | 0.92 |
| 80 | 0.33 | KEGG_PATHWAY | Morphine addiction | 6 | 253 | 91 | 9496 | 1.38 |
| 80 | 0.33 | KEGG_PATHWAY | Glutamatergic synapse | 5 | 253 | 116 | 9496 | 1.15 |
| 80 | 0.33 | KEGG_PATHWAY | Cholinergic synapse | 5 | 253 | 116 | 9496 | 1.15 |
| 80 | 0.33 | KEGG_PATHWAY | Relaxin signaling pathway | 4 | 253 | 130 | 9496 | 0.92 |

| Cluster | Cluster Enrichment Score | Category | Term | Count | List Total | Pop Hits | Pop Total | % |
|---|---|---|---|---|---|---|---|---|
| 80 | 0.33 | KEGG_PATHWAY | GABAergic synapse | 3 | 253 | 89 | 9496 | 0.69 |
| 80 | 0.33 | KEGG_PATHWAY | Dopaminergic synapse | 4 | 253 | 133 | 9496 | 0.92 |
| 80 | 0.33 | KEGG_PATHWAY | Circadian entrainment | 3 | 253 | 97 | 9496 | 0.69 |
| 80 | 0.33 | KEGG_PATHWAY | Retrograde endocannabinoid signaling | 4 | 253 | 149 | 9496 | 0.92 |
| 81 | 0.32 | UP_SEQ_FEATURE | DOMAIN:B box-type | 3 | 422 | 63 | 20669 | 0.69 |
| 81 | 0.32 | INTERPRO | Znf_B-box | 3 | 427 | 88 | 20827 | 0.69 |
| 81 | 0.32 | SMART | BBOX | 3 | 246 | 80 | 10704 | 0.69 |
| 82 | 0.30 | GOTERM_BP_DIRECT | positive regulation of defense response to virus by host | 5 | 409 | 34 | 19512 | 1.15 |
| 82 | 0.30 | GOTERM_BP_DIRECT | protein K48-linked ubiquitination | 4 | 409 | 106 | 19512 | 0.92 |
| 82 | 0.30 | UP_SEQ_FEATURE | ZN_FING:RING-type | 6 | 422 | 219 | 20669 | 1.38 |
| 82 | 0.30 | UP_SEQ_FEATURE | DOMAIN:RING-type | 5 | 422 | 210 | 20669 | 1.15 |
| 82 | 0.30 | SMART | RING | 6 | 246 | 259 | 10704 | 1.38 |
| 82 | 0.30 | INTERPRO | Znf_RING_CS | 4 | 427 | 183 | 20827 | 0.92 |
| 82 | 0.30 | INTERPRO | Znf_RING | 6 | 427 | 314 | 20827 | 1.38 |
| 82 | 0.30 | INTERPRO | Znf_RING/FYVE/PHD | 9 | 427 | 491 | 20827 | 2.06 |
| 82 | 0.30 | GOTERM_BP_DIRECT | ubiquitin-dependent protein catabolic process | 5 | 409 | 261 | 19512 | 1.15 |
| 82 | 0.30 | GOTERM_MF_DIRECT | ubiquitin protein ligase activity | 6 | 417 | 383 | 19272 | 1.38 |
| 82 | 0.30 | GOTERM_BP_DIRECT | proteasome-mediated ubiquitin-dependent protein catabolic process | 4 | 409 | 356 | 19512 | 0.92 |
| 82 | 0.30 | GOTERM_BP_DIRECT | protein ubiquitination | 6 | 409 | 550 | 19512 | 1.38 |
| 82 | 0.30 | UP_KW_BIOLOGICAL_PROCESS | Ubl conjugation pathway | 10 | 278 | 802 | 11605 | 2.29 |
| 83 | 0.27 | GOTERM_BP_DIRECT | protein autophosphorylation | 7 | 409 | 119 | 19512 | 1.61 |
| 83 | 0.27 | GOTERM_BP_DIRECT | protein phosphorylation | 8 | 409 | 261 | 19512 | 1.83 |
| 83 | 0.27 | INTERPRO | Tyr_kinase_AS | 4 | 427 | 98 | 20827 | 0.92 |
| 83 | 0.27 | UP_SEQ_FEATURE | ACT_SITE:Proton acceptor | 21 | 422 | 909 | 20669 | 4.82 |
| 83 | 0.27 | GOTERM_MF_DIRECT | kinase activity | 18 | 417 | 729 | 19272 | 4.13 |
| 83 | 0.27 | GOTERM_MF_DIRECT | nucleotide binding | 42 | 417 | 1839 | 19272 | 9.63 |

| Cluster | Cluster Enrichment Score | Category | Term | Count | List Total | Pop Hits | Pop Total | % |
|---|---|---|---|---|---|---|---|---|
| 83 | 0.27 | INTERPRO | Protein_kinase_ATP_BS | 9 | 427 | 390 | 20827 | 2.06 |
| 83 | 0.27 | INTERPRO | Prot_kinase_dom | 11 | 427 | 499 | 20827 | 2.52 |
| 83 | 0.27 | UP_SEQ_FEATURE | DOMAIN:Protein kinase | 11 | 422 | 504 | 20669 | 2.52 |
| 83 | 0.27 | GOTERM_MF_DIRECT | protein kinase activity | 12 | 417 | 526 | 19272 | 2.75 |
| 83 | 0.27 | UP_KW_MOLECULAR_FUNCTION | Kinase | 18 | 284 | 755 | 12059 | 4.13 |
| 83 | 0.27 | INTERPRO | Kinase-like_dom_sf | 11 | 427 | 545 | 20827 | 2.52 |
| 83 | 0.27 | SMART | S_TKc | 8 | 246 | 364 | 10704 | 1.83 |
| 83 | 0.27 | UP_KW_LIGAND | Nucleotide-binding | 41 | 165 | 1888 | 7011 | 9.40 |
| 83 | 0.27 | GOTERM_MF_DIRECT | protein serine kinase activity | 7 | 417 | 365 | 19272 | 1.61 |
| 83 | 0.27 | GOTERM_MF_DIRECT | ATP binding | 29 | 417 | 1549 | 19272 | 6.65 |
| 83 | 0.27 | INTERPRO | Ser/Thr_kinase_AS | 5 | 427 | 315 | 20827 | 1.15 |
| 83 | 0.27 | GOTERM_MF_DIRECT | protein serine/threonine kinase activity | 7 | 417 | 423 | 19272 | 1.61 |
| 83 | 0.27 | UP_KW_MOLECULAR_FUNCTION | Serine/threonine-protein kinase | 7 | 284 | 398 | 12059 | 1.61 |
| 83 | 0.27 | UP_KW_LIGAND | ATP-binding | 27 | 165 | 1452 | 7011 | 6.19 |
| 84 | 0.26 | INTERPRO | SAM/pointed_sf | 4 | 427 | 126 | 20827 | 0.92 |
| 84 | 0.26 | UP_SEQ_FEATURE | DOMAIN:SAM | 3 | 422 | 95 | 20669 | 0.69 |
| 84 | 0.26 | INTERPRO | SAM | 3 | 427 | 95 | 20827 | 0.69 |
| 85 | 0.22 | GOTERM_CC_DIRECT | focal adhesion | 10 | 415 | 438 | 20808 | 2.29 |
| 85 | 0.22 | GOTERM_CC_DIRECT | anchoring junction | 9 | 415 | 423 | 20808 | 2.06 |
| 85 | 0.22 | UP_KW_CELLULAR_COMPONENT | Cell junction | 9 | 387 | 425 | 18195 | 2.06 |
| 86 | 0.22 | KEGG_PATHWAY | Leukocyte transendothelial migration | 5 | 253 | 116 | 9496 | 1.15 |
| 86 | 0.22 | KEGG_PATHWAY | Diabetic cardiomyopathy | 6 | 253 | 205 | 9496 | 1.38 |
| 86 | 0.22 | KEGG_PATHWAY | Prion disease | 5 | 253 | 278 | 9496 | 1.15 |
| 87 | 0.21 | KEGG_PATHWAY | Aldosterone synthesis and secretion | 4 | 253 | 98 | 9496 | 0.92 |
| 87 | 0.21 | KEGG_PATHWAY | Cortisol synthesis and secretion | 3 | 253 | 65 | 9496 | 0.69 |
| 87 | 0.21 | KEGG_PATHWAY | Cushing syndrome | 3 | 253 | 155 | 9496 | 0.69 |
| 88 | 0.19 | KEGG_PATHWAY | Endocrine resistance | 4 | 253 | 99 | 9496 | 0.92 |

| Cluster | Cluster Enrichment Score | Category | Term | Count | List Total | Pop Hits | Pop Total | % |
|---|---|---|---|---|---|---|---|---|
| 88 | 0.19 | KEGG_PATHWAY | Relaxin signaling pathway | 4 | 253 | 130 | 9496 | 0.92[illegible] |
| 88 | 0.19 | KEGG_PATHWAY | Growth hormone synthesis, secretion and action | 3 | 253 | 122 | 9496 | 0.69[illegible] |
| 89 | 0.18 | GOTERM_CC_DIRECT | endoplasmic reticulum membrane | 26 | 415 | 1228 | 20808 | 5.96[illegible] |
| 89 | 0.18 | GOTERM_CC_DIRECT | endoplasmic reticulum | 34 | 415 | 1760 | 20808 | 7.80[illegible] |
| 89 | 0.18 | UP_KW_CELLULAR_COMPONENT | Endoplasmic reticulum | 26 | 387 | 1439 | 18195 | 5.96[illegible] |
| 90 | 0.16 | KEGG_PATHWAY | Peroxisome | 3 | 253 | 83 | 9496 | 0.69[illegible] |
| 90 | 0.16 | UP_KW_CELLULAR_COMPONENT | Peroxisome | 3 | 387 | 111 | 18195 | 0.69[illegible] |
| 90 | 0.16 | GOTERM_CC_DIRECT | peroxisome | 3 | 415 | 133 | 20808 | 0.69[illegible] |
| 91 | 0.16 | GOTERM_CC_DIRECT | mitochondrial matrix | 10 | 415 | 464 | 20808 | 2.29[illegible] |
| 91 | 0.16 | UP_SEQ_FEATURE | TRANSIT:Mitochondrion | 11 | 422 | 562 | 20669 | 2.52[illegible] |
| 91 | 0.16 | UP_KW_DOMAIN | Transit peptide | 12 | 313 | 590 | 14656 | 2.75[illegible] |
| 91 | 0.16 | UP_KW_CELLULAR_COMPONENT | Mitochondrion | 27 | 387 | 1403 | 18195 | 6.19[illegible] |
| 92 | 0.13 | GOTERM_BP_DIRECT | visual perception | 6 | 409 | 221 | 19512 | 1.38[illegible] |
| 92 | 0.13 | UP_KW_BIOLOGICAL_PROCESS | Vision | 3 | 278 | 127 | 11605 | 0.69[illegible] |
| 92 | 0.13 | UP_KW_BIOLOGICAL_PROCESS | Sensory transduction | 3 | 278 | 634 | 11605 | 0.69[illegible] |
| 93 | 0.12 | GOTERM_MF_DIRECT | metal ion binding | 94 | 417 | 4143 | 19272 | 21.5[illegible] |
| 93 | 0.12 | GOTERM_MF_DIRECT | zinc ion binding | 43 | 417 | 2237 | 19272 | 9.86[illegible] |
| 93 | 0.12 | UP_KW_LIGAND | Metal-binding | 94 | 165 | 4245 | 7011 | 21.5[illegible] |
| 93 | 0.12 | UP_KW_LIGAND | Zinc | 48 | 165 | 2593 | 7011 | 11.0[illegible] |
| 93 | 0.12 | UP_KW_DOMAIN | Zinc-finger | 23 | 313 | 1842 | 14656 | 5.28[illegible] |
| 94 | 0.11 | UP_SEQ_FEATURE | REPEAT:ANK 5 | 4 | 422 | 158 | 20669 | 0.92[illegible] |
| 94 | 0.11 | UP_SEQ_FEATURE | REPEAT:ANK 6 | 3 | 422 | 112 | 20669 | 0.69[illegible] |
| 94 | 0.11 | UP_SEQ_FEATURE | REPEAT:ANK 4 | 4 | 422 | 188 | 20669 | 0.92[illegible] |
| 94 | 0.11 | UP_SEQ_FEATURE | REPEAT:ANK | 4 | 422 | 198 | 20669 | 0.92[illegible] |
| 94 | 0.11 | UP_SEQ_FEATURE | REPEAT:ANK 1 | 5 | 422 | 260 | 20669 | 1.15[illegible] |
| 94 | 0.11 | UP_SEQ_FEATURE | REPEAT:ANK 2 | 5 | 422 | 261 | 20669 | 1.15[illegible] |
| 94 | 0.11 | INTERPRO | Ankyrin_rpt | 5 | 427 | 269 | 20827 | 1.15[illegible] |
| 94 | 0.11 | INTERPRO | Ankyrin_rpt-contain_sf | 5 | 427 | 275 | 20827 | 1.15[illegible] |

| Cluster | Cluster Enrichment Score | Category | Term | Count | List Total | Pop Hits | Pop Total | % |
|---|---|---|---|---|---|---|---|---|
| 94 | 0.11 | UP_KW_DOMAIN | ANK repeat | 5 | 313 | 267 | 14656 | 1.15 |
| 94 | 0.11 | UP_SEQ_FEATURE | REPEAT:ANK 3 | 4 | 422 | 222 | 20669 | 0.92 |
| 94 | 0.11 | SMART | ANK | 5 | 246 | 258 | 10704 | 1.15 |
| 95 | 0.10 | KEGG_PATHWAY | Oocyte meiosis | 4 | 253 | 139 | 9496 | 0.92 |
| 95 | 0.10 | KEGG_PATHWAY | Progesterone-mediated oocyte maturation | 3 | 253 | 111 | 9496 | 0.69 |
| 95 | 0.10 | KEGG_PATHWAY | Thermogenesis | 5 | 253 | 234 | 9496 | 1.15 |
| 96 | 0.08 | INTERPRO | Homeobox_CS | 4 | 427 | 194 | 20827 | 0.92 |
| 96 | 0.08 | INTERPRO | HD | 5 | 427 | 258 | 20827 | 1.15 |
| 96 | 0.08 | UP_KW_DOMAIN | Homeobox | 5 | 313 | 256 | 14656 | 1.15 |
| 96 | 0.08 | SMART | HOX | 5 | 246 | 252 | 10704 | 1.15 |
| 96 | 0.08 | INTERPRO | Homeodomain-like_sf | 5 | 427 | 328 | 20827 | 1.15 |
| 96 | 0.08 | UP_SEQ_FEATURE | DNA_BIND:Homeobox | 3 | 422 | 234 | 20669 | 0.69 |
| 97 | 0.07 | UP_SEQ_FEATURE | DOMAIN:PDZ | 3 | 422 | 152 | 20669 | 0.69 |
| 97 | 0.07 | INTERPRO | PDZ | 3 | 427 | 159 | 20827 | 0.69 |
| 97 | 0.07 | INTERPRO | PDZ_sf | 3 | 427 | 162 | 20827 | 0.69 |
| 97 | 0.07 | SMART | PDZ | 3 | 246 | 156 | 10704 | 0.69 |
| 98 | 0.01 | UP_SEQ_FEATURE | REPEAT:WD 7 | 3 | 422 | 206 | 20669 | 0.69 |
| 98 | 0.01 | UP_SEQ_FEATURE | REPEAT:WD | 3 | 422 | 211 | 20669 | 0.69 |
| 98 | 0.01 | INTERPRO | WD40/YVTN_repeat-like_dom_sf | 5 | 427 | 358 | 20827 | 1.15 |
| 98 | 0.01 | UP_SEQ_FEATURE | REPEAT:WD 6 | 3 | 422 | 258 | 20669 | 0.69 |
| 98 | 0.01 | UP_SEQ_FEATURE | REPEAT:WD 5 | 3 | 422 | 264 | 20669 | 0.69 |
| 98 | 0.01 | UP_SEQ_FEATURE | REPEAT:WD 4 | 3 | 422 | 267 | 20669 | 0.69 |
| 98 | 0.01 | UP_SEQ_FEATURE | REPEAT:WD 3 | 3 | 422 | 270 | 20669 | 0.69 |
| 98 | 0.01 | INTERPRO | WD40_rpt | 3 | 427 | 276 | 20827 | 0.69 |
| 98 | 0.01 | UP_SEQ_FEATURE | REPEAT:WD 2 | 3 | 422 | 278 | 20669 | 0.69 |
| 98 | 0.01 | UP_SEQ_FEATURE | REPEAT:WD 1 | 3 | 422 | 278 | 20669 | 0.69 |
| 98 | 0.01 | UP_KW_DOMAIN | WD repeat | 3 | 313 | 286 | 14656 | 0.69 |
| 98 | 0.01 | SMART | WD40 | 3 | 246 | 275 | 10704 | 0.69 |

| Cluster | Cluster Enrichment Score | Category | Term | Count | List Total | Pop Hits | Pop Total | % |
|---|---|---|---|---|---|---|---|---|
| 98 | 0.01 | INTERPRO | WD40_repeat_dom_sf | 3 | 427 | 312 | 20827 | 0.69 |
| 99 | 0.00 | GOTERM_BP_DIRECT | DNA damage response | 8 | 409 | 584 | 19512 | 1.83 |
| 99 | 0.00 | UP_KW_BIOLOGICAL_PROCESS | DNA damage | 4 | 278 | 445 | 11605 | 0.92 |
| 99 | 0.00 | UP_KW_BIOLOGICAL_PROCESS | DNA repair | 3 | 278 | 368 | 11605 | 0.69 |
| 99 | 0.00 | GOTERM_BP_DIRECT | DNA repair | 3 | 409 | 437 | 19512 | 0.69 |
| 100 | 0.00 | UP_SEQ_FEATURE | DOMAIN:C2H2-type | 4 | 422 | 521 | 20669 | 0.92 |
| 100 | 0.00 | UP_SEQ_FEATURE | ZN_FING:C2H2-type 2 | 5 | 422 | 633 | 20669 | 1.15 |
| 100 | 0.00 | UP_SEQ_FEATURE | ZN_FING:C2H2-type 4 | 4 | 422 | 608 | 20669 | 0.92 |
| 100 | 0.00 | INTERPRO | Znf_C2H2_sf | 6 | 427 | 796 | 20827 | 1.38 |
| 100 | 0.00 | INTERPRO | Znf_C2H2_type | 6 | 427 | 828 | 20827 | 1.38 |
| 100 | 0.00 | UP_SEQ_FEATURE | ZN_FING:C2H2-type 3 | 4 | 422 | 655 | 20669 | 0.92 |
| 100 | 0.00 | UP_SEQ_FEATURE | ZN_FING:C2H2-type 1 | 3 | 422 | 560 | 20669 | 0.69 |
| 100 | 0.00 | UP_SEQ_FEATURE | ZN_FING:C2H2-type 5 | 3 | 422 | 572 | 20669 | 0.69 |
| 100 | 0.00 | SMART | ZnF_C2H2 | 5 | 246 | 776 | 10704 | 1.15 |
