## Supplemental Table 2 for "CMPK2 restricts *Mycobacterium tuberculosis* replication and regulates macrophage gene expression"

| Cluster | Cluster Enrichment Score | Category | Term | Count | List Total | Pop Hits | Pop Total | % | [illegible] |
|---|---|---|---|---|---|---|---|---|---|
| 1 | 18.43 | INTERPRO | Cadherin_CBD | 22 | 314 | 37 | 20827 | 6.85 | [illegible] |
| 1 | 18.43 | INTERPRO | Cadherin_C | 21 | 314 | 41 | 20827 | 6.54 | [illegible] |
| 1 | 18.43 | INTERPRO | Protocadherin/Cadherin-CA | 24 | 314 | 73 | 20827 | 7.48 | [illegible] |
| 1 | 18.43 | INTERPRO | Cadherin_N | 23 | 314 | 65 | 20827 | 7.17 | [illegible] |
| 1 | 18.43 | UP_SEQ_FEATURE | DOMAIN:Cadherin 6 | 24 | 312 | 82 | 20669 | 7.48 | [illegible] |
| 1 | 18.43 | UP_SEQ_FEATURE | DOMAIN:Cadherin 1 | 26 | 312 | 114 | 20669 | 8.10 | [illegible] |
| 1 | 18.43 | UP_SEQ_FEATURE | DOMAIN:Cadherin 2 | 26 | 312 | 114 | 20669 | 8.10 | [illegible] |
| 1 | 18.43 | UP_SEQ_FEATURE | DOMAIN:Cadherin 5 | 25 | 312 | 104 | 20669 | 7.79 | [illegible] |
| 1 | 18.43 | INTERPRO | Cadherin-like_dom | 26 | 314 | 118 | 20827 | 8.10 | [illegible] |
| 1 | 18.43 | INTERPRO | Cadherin-like_sf | 26 | 314 | 121 | 20827 | 8.10 | [illegible] |
| 1 | 18.43 | UP_SEQ_FEATURE | DOMAIN:Cadherin 3 | 25 | 312 | 111 | 20669 | 7.79 | [illegible] |
| 1 | 18.43 | UP_SEQ_FEATURE | DOMAIN:Cadherin 4 | 25 | 312 | 111 | 20669 | 7.79 | [illegible] |
| 1 | 18.43 | INTERPRO | Cadherin_CS | 25 | 314 | 113 | 20827 | 7.79 | [illegible] |
| 1 | 18.43 | GOTERM_BP_DIRECT | homophilic cell-cell adhesion | 28 | 296 | 171 | 19512 | 8.72 | [illegible] |
| 1 | 18.43 | GOTERM_MF_DIRECT | cell adhesion molecule binding | 26 | 306 | 134 | 19272 | 8.10 | [illegible] |
| 1 | 18.43 | SMART | CA | 26 | 212 | 116 | 10704 | 8.10 | [illegible] |
| 1 | 18.43 | UP_KW_BIOLOGICAL_PROCESS | Cell adhesion | 43 | 193 | 494 | 11605 | 13.40 | [illegible] |
| 1 | 18.43 | GOTERM_BP_DIRECT | cell adhesion | 46 | 296 | 677 | 19512 | 14.33 | [illegible] |
| 1 | 18.43 | GOTERM_BP_DIRECT | nervous system development | 38 | 296 | 649 | 19512 | 11.84 | [illegible] |
| 1 | 18.43 | UP_KW_LIGAND | Calcium | 50 | 137 | 994 | 7011 | 15.58 | [illegible] |
| 1 | 18.43 | GOTERM_MF_DIRECT | calcium ion binding | 38 | 306 | 744 | 19272 | 11.84 | [illegible] |
| 1 | 18.43 | UP_SEQ_FEATURE | DOMAIN:Cadherin | 12 | 312 | 81 | 20669 | 3.74 | [illegible] |
| 1 | 18.43 | UP_SEQ_FEATURE | COMPBIAS:Basic residues | 55 | 312 | 2512 | 20669 | 17.13 | [illegible] |
| 2 | 5.88 | GOTERM_CC_DIRECT | plasma membrane | 146 | 308 | 5927 | 20808 | 45.48 | [illegible] |
| 2 | 5.88 | UP_SEQ_FEATURE | CARBOHYD:N-linked (GlcNAc...) asparagine | 113 | 312 | 4425 | 20669 | 35.20 | [illegible] |
| 2 | 5.88 | UP_KW_CELLULAR_COMPONENT | Cell membrane | 107 | 289 | 4200 | 18195 | 33.33 | [illegible] |
| 2 | 5.88 | GOTERM_CC_DIRECT | membrane | 179 | 308 | 9038 | 20808 | 55.76 | [illegible] |

| Cluster | Cluster Enrichment Score | Category | Term | Count | List Total | Pop Hits | Pop Total | % | [illegible] |
|---|---|---|---|---|---|---|---|---|---|
| 2 | 5.88 | UP_SEQ_FEATURE | TOPO_DOM:Cytoplasmic | 96 | 312 | 3928 | 20669 | 29.91 | [illegible] |
| 2 | 5.88 | UP_SEQ_FEATURE | TOPO_DOM:Extracellular | 79 | 312 | 3012 | 20669 | 24.61 | [illegible] |
| 2 | 5.88 | UP_KW_DOMAIN | Signal | 104 | 237 | 4426 | 14656 | 32.40 | [illegible] |
| 2 | 5.88 | UP_KW_CELLULAR_COMPONENT | Membrane | 169 | 289 | 8428 | 18195 | 52.65 | [illegible] |
| 2 | 5.88 | UP_KW_PTM | Glycoprotein | 119 | 258 | 4892 | 14452 | 37.07 | [illegible] |
| 2 | 5.88 | UP_SEQ_FEATURE | TRANSMEM:Helical | 110 | 312 | 5434 | 20669 | 34.27 | [illegible] |
| 2 | 5.88 | UP_KW_DOMAIN | Transmembrane helix | 121 | 237 | 5895 | 14656 | 37.69 | [illegible] |
| 2 | 5.88 | UP_KW_DOMAIN | Transmembrane | 121 | 237 | 5971 | 14656 | 37.69 | [illegible] |
| 3 | 4.91 | GOTERM_CC_DIRECT | anchoring junction | 22 | 308 | 423 | 20808 | 6.85 | [illegible] |
| 3 | 4.91 | UP_KW_CELLULAR_COMPONENT | Cell junction | 22 | 289 | 425 | 18195 | 6.85 | [illegible] |
| 3 | 4.91 | GOTERM_CC_DIRECT | focal adhesion | 18 | 308 | 438 | 20808 | 5.61 | [illegible] |
| 4 | 2.99 | KEGG_PATHWAY | Focal adhesion | 12 | 138 | 203 | 9496 | 3.74 | [illegible] |
| 4 | 2.99 | GOTERM_MF_DIRECT | integrin binding | 11 | 306 | 161 | 19272 | 3.43 | [illegible] |
| 4 | 2.99 | KEGG_PATHWAY | Regulation of actin cytoskeleton | 12 | 138 | 232 | 9496 | 3.74 | [illegible] |
| 4 | 2.99 | KEGG_PATHWAY | PI3K-Akt signaling pathway | 15 | 138 | 362 | 9496 | 4.67 | [illegible] |
| 4 | 2.99 | KEGG_PATHWAY | Human papillomavirus infection | 14 | 138 | 333 | 9496 | 4.36 | [illegible] |
| 4 | 2.99 | KEGG_PATHWAY | ECM-receptor interaction | 7 | 138 | 89 | 9496 | 2.18 | [illegible] |
| 4 | 2.99 | KEGG_PATHWAY | Small cell lung cancer | 5 | 138 | 93 | 9496 | 1.56 | [illegible] |
| 5 | 2.08 | INTERPRO | RGS_subdom1/3 | 5 | 314 | 18 | 20827 | 1.56 | [illegible] |
| 5 | 2.08 | GOTERM_BP_DIRECT | regulation of G protein-coupled receptor signaling pathway | 6 | 296 | 48 | 19512 | 1.87 | [illegible] |
| 5 | 2.08 | GOTERM_BP_DIRECT | negative regulation of signal transduction | 7 | 296 | 79 | 19512 | 2.18 | [illegible] |
| 5 | 2.08 | INTERPRO | RGS | 5 | 314 | 36 | 20827 | 1.56 | [illegible] |
| 5 | 2.08 | UP_SEQ_FEATURE | DOMAIN:RGS | 5 | 312 | 36 | 20669 | 1.56 | [illegible] |
| 5 | 2.08 | INTERPRO | RGS_subdomain_2 | 5 | 314 | 38 | 20827 | 1.56 | [illegible] |
| 5 | 2.08 | INTERPRO | RGS_sf | 5 | 314 | 39 | 20827 | 1.56 | [illegible] |
| 5 | 2.08 | SMART | RGS | 5 | 212 | 35 | 10704 | 1.56 | [illegible] |
| 5 | 2.08 | GOTERM_MF_DIRECT | calmodulin binding | 10 | 306 | 208 | 19272 | 3.12 | [illegible] |

| Cluster | Cluster Enrichment Score | Category | Term | Count | List Total | Pop Hits | Pop Total | % | [illegible] |
|---|---|---|---|---|---|---|---|---|---|
| 5 | 2.08 | UP_KW_MOLECULAR_FUNCTION | Signal transduction inhibitor | 5 | 187 | 49 | 12059 | 1.56 | [illegible] |
| 5 | 2.08 | GOTERM_MF_DIRECT | G-protein alpha-subunit binding | 4 | 306 | 30 | 19272 | 1.25 | [illegible] |
| 5 | 2.08 | GOTERM_BP_DIRECT | negative regulation of G protein-coupled receptor signaling pathway | 3 | 296 | 17 | 19512 | 0.93 | [illegible] |
| 5 | 2.08 | GOTERM_MF_DIRECT | GTPase activity | 11 | 306 | 364 | 19272 | 3.43 | [illegible] |
| 5 | 2.08 | GOTERM_MF_DIRECT | GTPase activator activity | 8 | 306 | 292 | 19272 | 2.49 | [illegible] |
| 5 | 2.08 | UP_KW_MOLECULAR_FUNCTION | GTPase activation | 6 | 187 | 216 | 12059 | 1.87 | [illegible] |
| 5 | 2.08 | GOTERM_BP_DIRECT | G protein-coupled receptor signaling pathway | 11 | 296 | 1041 | 19512 | 3.43 | [illegible] |
| 6 | 2.02 | GOTERM_BP_DIRECT | mesodermal cell differentiation | 5 | 296 | 14 | 19512 | 1.56 | [illegible] |
| 6 | 2.02 | GOTERM_MF_DIRECT | integrin binding | 11 | 306 | 161 | 19272 | 3.43 | [illegible] |
| 6 | 2.02 | GOTERM_BP_DIRECT | integrin-mediated signaling pathway | 9 | 296 | 114 | 19512 | 2.80 | [illegible] |
| 6 | 2.02 | GOTERM_BP_DIRECT | positive regulation of fibroblast migration | 4 | 296 | 14 | 19512 | 1.25 | [illegible] |
| 6 | 2.02 | GOTERM_MF_DIRECT | fibronectin binding | 5 | 306 | 30 | 19272 | 1.56 | [illegible] |
| 6 | 2.02 | KEGG_PATHWAY | ECM-receptor interaction | 7 | 138 | 89 | 9496 | 2.18 | [illegible] |
| 6 | 2.02 | KEGG_PATHWAY | Dilated cardiomyopathy | 7 | 138 | 105 | 9496 | 2.18 | [illegible] |
| 6 | 2.02 | UP_KW_MOLECULAR_FUNCTION | Integrin | 5 | 187 | 46 | 12059 | 1.56 | [illegible] |
| 6 | 2.02 | INTERPRO | Integrin_dom_sf | 4 | 314 | 26 | 20827 | 1.25 | [illegible] |
| 6 | 2.02 | GOTERM_BP_DIRECT | cell-substrate adhesion | 4 | 296 | 26 | 19512 | 1.25 | [illegible] |
| 6 | 2.02 | GOTERM_CC_DIRECT | integrin complex | 4 | 308 | 28 | 20808 | 1.25 | [illegible] |
| 6 | 2.02 | GOTERM_BP_DIRECT | heterotypic cell-cell adhesion | 4 | 296 | 31 | 19512 | 1.25 | [illegible] |
| 6 | 2.02 | GOTERM_BP_DIRECT | cell adhesion mediated by integrin | 4 | 296 | 39 | 19512 | 1.25 | [illegible] |
| 6 | 2.02 | GOTERM_MF_DIRECT | protease binding | 6 | 306 | 113 | 19272 | 1.87 | [illegible] |
| 6 | 2.02 | KEGG_PATHWAY | Arrhythmogenic right ventricular cardiomyopathy | 5 | 138 | 86 | 9496 | 1.56 | [illegible] |
| 6 | 2.02 | UP_SEQ_FEATURE | DOMAIN:VWFA | 5 | 312 | 85 | 20669 | 1.56 | [illegible] |
| 6 | 2.02 | KEGG_PATHWAY | Cytoskeleton in muscle | 8 | 138 | 233 | 9496 | 2.49 | [illegible] |

| Cluster | Cluster Enrichment Score | Category | Term | Count | List Total | Pop Hits | Pop Total | % | [illegible] |
|---|---|---|---|---|---|---|---|---|---|
| | | | cells | | | | | | |
| 6 | 2.02 | KEGG_PATHWAY | Hypertrophic cardiomyopathy | 5 | 138 | 99 | 9496 | 1.56 | [illegible] |
| 6 | 2.02 | GOTERM_BP_DIRECT | cell-cell adhesion | 8 | 296 | 225 | 19512 | 2.49 | [illegible] |
| 6 | 2.02 | INTERPRO | vWFA_dom_sf | 5 | 314 | 102 | 20827 | 1.56 | [illegible] |
| 6 | 2.02 | GOTERM_BP_DIRECT | cell-matrix adhesion | 5 | 296 | 108 | 19512 | 1.56 | [illegible] |
| 6 | 2.02 | GOTERM_CC_DIRECT | synaptic membrane | 4 | 308 | 73 | 20808 | 1.25 | [illegible] |
| 6 | 2.02 | GOTERM_MF_DIRECT | signaling receptor activity | 6 | 306 | 339 | 19272 | 1.87 | [illegible] |
| 7 | 1.80 | INTERPRO | WH1/EVH1_dom | 3 | 314 | 12 | 20827 | 0.93 | [illegible] |
| 7 | 1.80 | UP_SEQ_FEATURE | DOMAIN:WH1 | 3 | 312 | 13 | 20669 | 0.93 | [illegible] |
| 7 | 1.80 | SMART | WH1 | 3 | 212 | 11 | 10704 | 0.93 | [illegible] |
| 8 | 1.72 | INTERPRO | PH-like_dom_sf | 18 | 314 | 452 | 20827 | 5.61 | [illegible] |
| 8 | 1.72 | INTERPRO | PH_domain | 10 | 314 | 276 | 20827 | 3.12 | [illegible] |
| 8 | 1.72 | UP_SEQ_FEATURE | DOMAIN:PH | 9 | 312 | 283 | 20669 | 2.80 | [illegible] |
| 8 | 1.72 | SMART | PH | 9 | 212 | 267 | 10704 | 2.80 | [illegible] |
| 9 | 1.72 | GOTERM_CC_DIRECT | Schaffer collateral - CA1 synapse | 8 | 308 | 106 | 20808 | 2.49 | [illegible] |
| 9 | 1.72 | GOTERM_BP_DIRECT | modulation of chemical synaptic transmission | 7 | 296 | 123 | 19512 | 2.18 | [illegible] |
| 9 | 1.72 | GOTERM_CC_DIRECT | presynaptic membrane | 3 | 308 | 148 | 20808 | 0.93 | [illegible] |
| 10 | 1.50 | GOTERM_BP_DIRECT | response to wounding | 7 | 296 | 82 | 19512 | 2.18 | [illegible] |
| 10 | 1.50 | GOTERM_BP_DIRECT | platelet activation | 5 | 296 | 69 | 19512 | 1.56 | [illegible] |
| 10 | 1.50 | KEGG_PATHWAY | Complement and coagulation cascades | 4 | 138 | 88 | 9496 | 1.25 | [illegible] |
| 10 | 1.50 | GOTERM_BP_DIRECT | blood coagulation | 4 | 296 | 111 | 19512 | 1.25 | [illegible] |
| 11 | 1.45 | KEGG_PATHWAY | Fluid shear stress and atherosclerosis | 9 | 138 | 142 | 9496 | 2.80 | [illegible] |
| 11 | 1.45 | GOTERM_BP_DIRECT | blood vessel development | 6 | 296 | 71 | 19512 | 1.87 | [illegible] |
| 11 | 1.45 | GOTERM_BP_DIRECT | pituitary gland development | 4 | 296 | 30 | 19512 | 1.25 | [illegible] |
| 11 | 1.45 | GOTERM_BP_DIRECT | cellular response to BMP stimulus | 4 | 296 | 33 | 19512 | 1.25 | [illegible] |
| 11 | 1.45 | GOTERM_BP_DIRECT | stem cell differentiation | 5 | 296 | 64 | 19512 | 1.56 | [illegible] |

| Cluster | Cluster Enrichment Score | Category | Term | Count | List Total | Pop Hits | Pop Total | % |
|---|---|---|---|---|---|---|---|---|
| 11 | 1.45 | GOTERM_BP_DIRECT | heart morphogenesis | 5 | 296 | 66 | 19512 | 1.56 |
| 11 | 1.45 | GOTERM_BP_DIRECT | mesoderm formation | 4 | 296 | 40 | 19512 | 1.25 |
| 11 | 1.45 | KEGG_PATHWAY | Hippo signaling pathway | 7 | 138 | 157 | 9496 | 2.18 |
| 11 | 1.45 | GOTERM_BP_DIRECT | positive regulation of SMAD protein signal transduction | 4 | 296 | 43 | 19512 | 1.25 |
| 11 | 1.45 | GOTERM_BP_DIRECT | positive regulation of bone mineralization | 4 | 296 | 43 | 19512 | 1.25 |
| 11 | 1.45 | GOTERM_BP_DIRECT | lung alveolus development | 4 | 296 | 50 | 19512 | 1.25 |
| 11 | 1.45 | GOTERM_BP_DIRECT | positive regulation of cell differentiation | 5 | 296 | 86 | 19512 | 1.56 |
| 11 | 1.45 | GOTERM_BP_DIRECT | regulation of multicellular organismal development | 3 | 296 | 22 | 19512 | 0.93 |
| 11 | 1.45 | KEGG_PATHWAY | Signaling pathways regulating pluripotency of stem cells | 6 | 138 | 144 | 9496 | 1.87 |
| 11 | 1.45 | GOTERM_BP_DIRECT | osteoblast differentiation | 6 | 296 | 138 | 19512 | 1.87 |
| 11 | 1.45 | GOTERM_BP_DIRECT | embryonic digit morphogenesis | 4 | 296 | 60 | 19512 | 1.25 |
| 11 | 1.45 | GOTERM_BP_DIRECT | outflow tract morphogenesis | 4 | 296 | 63 | 19512 | 1.25 |
| 11 | 1.45 | KEGG_PATHWAY | TGF-beta signaling pathway | 5 | 138 | 110 | 9496 | 1.56 |
| 11 | 1.45 | GOTERM_BP_DIRECT | chondrocyte differentiation | 4 | 296 | 65 | 19512 | 1.25 |
| 11 | 1.45 | GOTERM_BP_DIRECT | cellular response to growth factor stimulus | 4 | 296 | 79 | 19512 | 1.25 |
| 11 | 1.45 | GOTERM_BP_DIRECT | ventricular septum morphogenesis | 3 | 296 | 41 | 19512 | 0.93 |
| 11 | 1.45 | GOTERM_BP_DIRECT | BMP signaling pathway | 4 | 296 | 91 | 19512 | 1.25 |
| 11 | 1.45 | GOTERM_BP_DIRECT | positive regulation of osteoblast differentiation | 3 | 296 | 80 | 19512 | 0.93 |
| 12 | 1.45 | GOTERM_MF_DIRECT | protein kinase activity | 19 | 306 | 526 | 19272 | 5.92 |
| 12 | 1.45 | GOTERM_MF_DIRECT | nucleotide binding | 46 | 306 | 1839 | 19272 | 14.33 |
| 12 | 1.45 | UP_KW_MOLECULAR_FUNCTION | Tyrosine-protein kinase | 8 | 187 | 118 | 12059 | 2.49 |
| 12 | 1.45 | INTERPRO | Protein_kinase_ATP_BS | 15 | 314 | 390 | 20827 | 4.67 |

| Cluster | Cluster Enrichment Score | Category | Term | Count | List Total | Pop Hits | Pop Total | % | [illegible] |
|---|---|---|---|---|---|---|---|---|---|
| 12 | 1.45 | GOTERM_BP_DIRECT | peptidyl-tyrosine phosphorylation | 5 | 296 | 41 | 19512 | 1.56 | [illegible] |
| 12 | 1.45 | INTERPRO | Tyr_kinase_AS | 7 | 314 | 98 | 20827 | 2.18 | [illegible] |
| 12 | 1.45 | INTERPRO | Prot_kinase_dom | 17 | 314 | 499 | 20827 | 5.30 | [illegible] |
| 12 | 1.45 | INTERPRO | Kinase-like_dom_sf | 18 | 314 | 545 | 20827 | 5.61 | [illegible] |
| 12 | 1.45 | UP_SEQ_FEATURE | DOMAIN:Protein kinase | 17 | 312 | 504 | 20669 | 5.30 | [illegible] |
| 12 | 1.45 | GOTERM_MF_DIRECT | protein tyrosine kinase activity | 8 | 306 | 140 | 19272 | 2.49 | [illegible] |
| 12 | 1.45 | GOTERM_MF_DIRECT | ATP binding | 38 | 306 | 1549 | 19272 | 11.84 | [illegible] |
| 12 | 1.45 | INTERPRO | Ser-Thr/ Tyr_kinase_cat_dom | 8 | 314 | 151 | 20827 | 2.49 | [illegible] |
| 12 | 1.45 | INTERPRO | Tyr_kinase_cat_dom | 6 | 314 | 91 | 20827 | 1.87 | [illegible] |
| 12 | 1.45 | GOTERM_MF_DIRECT | kinase activity | 20 | 306 | 729 | 19272 | 6.23 | [illegible] |
| 12 | 1.45 | SMART | TyrKc | 6 | 212 | 91 | 10704 | 1.87 | [illegible] |
| 12 | 1.45 | GOTERM_MF_DIRECT | non-membrane spanning protein tyrosine kinase activity | 4 | 306 | 46 | 19272 | 1.25 | [illegible] |
| 12 | 1.45 | GOTERM_BP_DIRECT | cell surface receptor protein tyrosine kinase signaling pathway | 6 | 296 | 125 | 19512 | 1.87 | [illegible] |
| 12 | 1.45 | UP_KW_MOLECULAR_FUNCTION | Kinase | 19 | 187 | 755 | 12059 | 5.92 | [illegible] |
| 12 | 1.45 | GOTERM_BP_DIRECT | protein phosphorylation | 9 | 296 | 261 | 19512 | 2.80 | [illegible] |
| 12 | 1.45 | UP_KW_LIGAND | ATP-binding | 37 | 137 | 1452 | 7011 | 11.53 | [illegible] |
| 12 | 1.45 | UP_KW_LIGAND | Nucleotide-binding | 46 | 137 | 1888 | 7011 | 14.33 | [illegible] |
| 12 | 1.45 | UP_SEQ_FEATURE | ACT_SITE:Proton acceptor | 20 | 312 | 909 | 20669 | 6.23 | [illegible] |
| 12 | 1.45 | INTERPRO | RTK | 3 | 314 | 44 | 20827 | 0.93 | [illegible] |
| 12 | 1.45 | GOTERM_MF_DIRECT | transmembrane receptor protein tyrosine kinase activity | 3 | 306 | 53 | 19272 | 0.93 | [illegible] |
| 12 | 1.45 | GOTERM_BP_DIRECT | protein autophosphorylation | 4 | 296 | 119 | 19512 | 1.25 | [illegible] |
| 12 | 1.45 | UP_KW_MOLECULAR_FUNCTION | Serine/threonine-protein kinase | 9 | 187 | 398 | 12059 | 2.80 | [illegible] |
| 12 | 1.45 | GOTERM_MF_DIRECT | protein serine/threonine kinase activity | 9 | 306 | 423 | 19272 | 2.80 | [illegible] |

| Cluster | Cluster Enrichment Score | Category | Term | Count | List Total | Pop Hits | Pop Total | % | [illegible] |
|---|---|---|---|---|---|---|---|---|---|
| 12 | 1.45 | GOTERM_MF_DIRECT | transferase activity | 33 | 306 | 1913 | 19272 | 10.28 | [illegible] |
| 12 | 1.45 | SMART | S_TKc | 9 | 212 | 364 | 10704 | 2.80 | [illegible] |
| 12 | 1.45 | UP_KW_MOLECULAR_FUNCTION | Transferase | 34 | 187 | 2093 | 12059 | 10.59 | [illegible] |
| 12 | 1.45 | INTERPRO | Immunoglobulin_dom | 3 | 314 | 110 | 20827 | 0.93 | [illegible] |
| 12 | 1.45 | GOTERM_MF_DIRECT | protein serine kinase activity | 6 | 306 | 365 | 19272 | 1.87 | [illegible] |
| 12 | 1.45 | INTERPRO | Ser/Thr_kinase_AS | 5 | 314 | 315 | 20827 | 1.56 | [illegible] |
| 13 | 1.42 | UP_KW_BIOLOGICAL_PROCESS | Endocytosis | 7 | 193 | 130 | 11605 | 2.18 | [illegible] |
| 13 | 1.42 | GOTERM_BP_DIRECT | endocytosis | 9 | 296 | 226 | 19512 | 2.80 | [illegible] |
| 13 | 1.42 | GOTERM_BP_DIRECT | receptor-mediated endocytosis | 4 | 296 | 80 | 19512 | 1.25 | [illegible] |
| 14 | 1.36 | UP_KW_BIOLOGICAL_PROCESS | Endocytosis | 7 | 193 | 130 | 11605 | 2.18 | [illegible] |
| 14 | 1.36 | GOTERM_MF_DIRECT | clathrin adaptor activity | 3 | 306 | 18 | 19272 | 0.93 | [illegible] |
| 14 | 1.36 | GOTERM_MF_DIRECT | clathrin binding | 3 | 306 | 38 | 19272 | 0.93 | [illegible] |
| 15 | 1.36 | GOTERM_BP_DIRECT | odontogenesis of dentin-containing tooth | 5 | 296 | 56 | 19512 | 1.56 | [illegible] |
| 15 | 1.36 | GOTERM_BP_DIRECT | positive regulation of miRNA transcription | 5 | 296 | 61 | 19512 | 1.56 | [illegible] |
| 15 | 1.36 | GOTERM_BP_DIRECT | endoderm development | 4 | 296 | 34 | 19512 | 1.25 | [illegible] |
| 15 | 1.36 | GOTERM_BP_DIRECT | epithelial cell proliferation | 5 | 296 | 72 | 19512 | 1.56 | [illegible] |
| 15 | 1.36 | KEGG_PATHWAY | Hippo signaling pathway | 7 | 138 | 157 | 9496 | 2.18 | [illegible] |
| 15 | 1.36 | GOTERM_BP_DIRECT | positive regulation of SMAD protein signal transduction | 4 | 296 | 43 | 19512 | 1.25 | [illegible] |
| 15 | 1.36 | GOTERM_BP_DIRECT | hematopoietic progenitor cell differentiation | 5 | 296 | 76 | 19512 | 1.56 | [illegible] |
| 15 | 1.36 | GOTERM_BP_DIRECT | positive regulation of cell differentiation | 5 | 296 | 86 | 19512 | 1.56 | [illegible] |
| 15 | 1.36 | KEGG_PATHWAY | TGF-beta signaling pathway | 5 | 138 | 110 | 9496 | 1.56 | [illegible] |
| 15 | 1.36 | GOTERM_BP_DIRECT | chondrocyte differentiation | 4 | 296 | 65 | 19512 | 1.25 | [illegible] |
| 15 | 1.36 | GOTERM_BP_DIRECT | positive regulation of epithelial cell proliferation | 4 | 296 | 74 | 19512 | 1.25 | [illegible] |
| 15 | 1.36 | GOTERM_BP_DIRECT | cellular response to growth factor stimulus | 4 | 296 | 79 | 19512 | 1.25 | [illegible] |

| Cluster | Cluster Enrichment Score | Category | Term | Count | List Total | Pop Hits | Pop Total | % | [illegible] |
|---|---|---|---|---|---|---|---|---|---|
| 15 | 1.36 | GOTERM_BP_DIRECT | negative regulation of gene expression | 5 | 296 | 351 | 19512 | 1.56 | [illegible] |
| 16 | 1.24 | INTERPRO | Tyr_Pase_cat | 5 | 314 | 63 | 20827 | 1.56 | [illegible] |
| 16 | 1.24 | INTERPRO | Prot-tyrosine_phosphatase-like | 6 | 314 | 105 | 20827 | 1.87 | [illegible] |
| 16 | 1.24 | GOTERM_MF_DIRECT | protein tyrosine phosphatase activity | 6 | 306 | 101 | 19272 | 1.87 | [illegible] |
| 16 | 1.24 | SMART | PTPc_motif | 5 | 212 | 63 | 10704 | 1.56 | [illegible] |
| 16 | 1.24 | INTERPRO | Tyr_Pase_dom | 5 | 314 | 90 | 20827 | 1.56 | [illegible] |
| 16 | 1.24 | UP_KW_MOLECULAR_FUNCTION | Protein phosphatase | 6 | 187 | 144 | 12059 | 1.87 | [illegible] |
| 16 | 1.24 | GOTERM_MF_DIRECT | phosphoprotein phosphatase activity | 6 | 306 | 153 | 19272 | 1.87 | [illegible] |
| 16 | 1.24 | UP_SEQ_FEATURE | DOMAIN:Tyrosine-protein phosphatase | 4 | 312 | 78 | 20669 | 1.25 | [illegible] |
| 16 | 1.24 | UP_SEQ_FEATURE | ACT_SITE:Phosphocysteine intermediate | 4 | 312 | 91 | 20669 | 1.25 | [illegible] |
| 16 | 1.24 | UP_SEQ_FEATURE | DOMAIN:Tyrosine specific protein phosphatases | 3 | 312 | 69 | 20669 | 0.93 | [illegible] |
| 17 | 1.22 | UP_SEQ_FEATURE | MOTIF:Cell attachment site | 8 | 312 | 88 | 20669 | 2.49 | [illegible] |
| 17 | 1.22 | INTERPRO | EGF-like_dom | 11 | 314 | 254 | 20827 | 3.43 | [illegible] |
| 17 | 1.22 | UP_KW_DOMAIN | EGF-like domain | 11 | 237 | 269 | 14656 | 3.43 | [illegible] |
| 17 | 1.22 | GOTERM_CC_DIRECT | extracellular matrix | 15 | 308 | 492 | 20808 | 4.67 | [illegible] |
| 17 | 1.22 | SMART | EGF | 10 | 212 | 200 | 10704 | 3.12 | [illegible] |
| 17 | 1.22 | UP_SEQ_FEATURE | DOMAIN:EGF-like | 8 | 312 | 221 | 20669 | 2.49 | [illegible] |
| 17 | 1.22 | GOTERM_MF_DIRECT | extracellular matrix structural constituent | 6 | 306 | 127 | 19272 | 1.87 | [illegible] |
| 17 | 1.22 | INTERPRO | Growth_fac_rcpt_cys_sf | 6 | 314 | 145 | 20827 | 1.87 | [illegible] |
| 17 | 1.22 | INTERPRO | cEGF | 3 | 314 | 30 | 20827 | 0.93 | [illegible] |
| 17 | 1.22 | INTERPRO | NOTCH1_EGF-like | 4 | 314 | 80 | 20827 | 1.25 | [illegible] |
| 17 | 1.22 | UP_KW_CELLULAR_COMPONENT | Extracellular matrix | 8 | 289 | 282 | 18195 | 2.49 | [illegible] |
| 17 | 1.22 | UP_SEQ_FEATURE | DOMAIN:EGF-like 5; calcium-binding | 3 | 312 | 49 | 20669 | 0.93 | [illegible] |
| 17 | 1.22 | UP_SEQ_FEATURE | DOMAIN:EGF-like 2; calcium-binding | 3 | 312 | 57 | 20669 | 0.93 | [illegible] |

| Cluster | Cluster Enrichment Score | Category | Term | Count | List Total | Pop Hits | Pop Total | % | [illegible] |
|---|---|---|---|---|---|---|---|---|---|
| 17 | 1.22 | INTERPRO | EGF-type_Asp/ Asn_hydroxyl_site | 4 | 314 | 107 | 20827 | 1.25 | [illegible] |
| 17 | 1.22 | INTERPRO | EGF_Ca-bd_CS | 4 | 314 | 109 | 20827 | 1.25 | [illegible] |
| 17 | 1.22 | UP_SEQ_FEATURE | DOMAIN:EGF-like 1 | 4 | 312 | 129 | 20669 | 1.25 | [illegible] |
| 17 | 1.22 | INTERPRO | EGF-like_Ca-bd_dom | 4 | 314 | 134 | 20827 | 1.25 | [illegible] |
| 17 | 1.22 | SMART | EGF_CA | 4 | 212 | 134 | 10704 | 1.25 | [illegible] |
| 18 | 1.18 | UP_SEQ_FEATURE | DOMAIN:SH3 | 8 | 312 | 217 | 20669 | 2.49 | [illegible] |
| 18 | 1.18 | INTERPRO | SH3-like_dom_sf | 8 | 314 | 220 | 20827 | 2.49 | [illegible] |
| 18 | 1.18 | INTERPRO | SH3_domain | 8 | 314 | 229 | 20827 | 2.49 | [illegible] |
| 18 | 1.18 | UP_KW_DOMAIN | SH3 domain | 8 | 237 | 229 | 14656 | 2.49 | [illegible] |
| 18 | 1.18 | SMART | SH3 | 8 | 212 | 208 | 10704 | 2.49 | [illegible] |
| 19 | 1.12 | GOTERM_CC_DIRECT | clathrin-coated vesicle membrane | 5 | 308 | 35 | 20808 | 1.56 | [illegible] |
| 19 | 1.12 | GOTERM_BP_DIRECT | actin filament organization | 5 | 296 | 158 | 19512 | 1.56 | [illegible] |
| 19 | 1.12 | GOTERM_MF_DIRECT | actin filament binding | 6 | 306 | 213 | 19272 | 1.87 | [illegible] |
| 19 | 1.12 | GOTERM_CC_DIRECT | clathrin-coated vesicle | 3 | 308 | 83 | 20808 | 0.93 | [illegible] |
| 20 | 0.99 | KEGG_PATHWAY | Platelet activation | 10 | 138 | 126 | 9496 | 3.12 | [illegible] |
| 20 | 0.99 | KEGG_PATHWAY | Human cytomegalovirus infection | 11 | 138 | 227 | 9496 | 3.43 | [illegible] |
| 20 | 0.99 | KEGG_PATHWAY | Endocrine resistance | 6 | 138 | 99 | 9496 | 1.87 | [illegible] |
| 20 | 0.99 | KEGG_PATHWAY | Yersinia infection | 7 | 138 | 138 | 9496 | 2.18 | [illegible] |
| 20 | 0.99 | KEGG_PATHWAY | Apelin signaling pathway | 7 | 138 | 140 | 9496 | 2.18 | [illegible] |
| 20 | 0.99 | KEGG_PATHWAY | Oxytocin signaling pathway | 7 | 138 | 155 | 9496 | 2.18 | [illegible] |
| 20 | 0.99 | KEGG_PATHWAY | Parathyroid hormone synthesis, secretion and action | 6 | 138 | 115 | 9496 | 1.87 | [illegible] |
| 20 | 0.99 | KEGG_PATHWAY | Growth hormone synthesis, secretion and action | 6 | 138 | 122 | 9496 | 1.87 | [illegible] |
| 20 | 0.99 | KEGG_PATHWAY | cGMP-PKG signaling pathway | 7 | 138 | 166 | 9496 | 2.18 | [illegible] |
| 20 | 0.99 | KEGG_PATHWAY | Relaxin signaling pathway | 6 | 138 | 130 | 9496 | 1.87 | [illegible] |
| 20 | 0.99 | KEGG_PATHWAY | Gap junction | 5 | 138 | 92 | 9496 | 1.56 | [illegible] |
| 20 | 0.99 | KEGG_PATHWAY | GnRH signaling pathway | 5 | 138 | 93 | 9496 | 1.56 | [illegible] |

| Cluster | Cluster Enrichment Score | Category | Term | Count | List Total | Pop Hits | Pop Total | % | [illegible] |
|---|---|---|---|---|---|---|---|---|---|
| 20 | 0.99 | KEGG_PATHWAY | Regulation of lipolysis in adipocytes | 4 | 138 | 59 | 9496 | 1.25 | [illegible] |
| 20 | 0.99 | KEGG_PATHWAY | Melanogenesis | 5 | 138 | 101 | 9496 | 1.56 | [illegible] |
| 20 | 0.99 | KEGG_PATHWAY | Leukocyte transendothelial migration | 5 | 138 | 116 | 9496 | 1.56 | [illegible] |
| 20 | 0.99 | KEGG_PATHWAY | Glutamatergic synapse | 5 | 138 | 116 | 9496 | 1.56 | [illegible] |
| 20 | 0.99 | KEGG_PATHWAY | Gastric acid secretion | 4 | 138 | 76 | 9496 | 1.25 | [illegible] |
| 20 | 0.99 | KEGG_PATHWAY | cAMP signaling pathway | 7 | 138 | 226 | 9496 | 2.18 | [illegible] |
| 20 | 0.99 | KEGG_PATHWAY | Chemokine signaling pathway | 6 | 138 | 193 | 9496 | 1.87 | [illegible] |
| 20 | 0.99 | KEGG_PATHWAY | Calcium signaling pathway | 7 | 138 | 254 | 9496 | 2.18 | [illegible] |
| 20 | 0.99 | KEGG_PATHWAY | Cushing syndrome | 5 | 138 | 155 | 9496 | 1.56 | [illegible] |
| 20 | 0.99 | KEGG_PATHWAY | Chemical carcinogenesis - receptor activation | 6 | 138 | 217 | 9496 | 1.87 | [illegible] |
| 20 | 0.99 | KEGG_PATHWAY | Hormone signaling | 6 | 138 | 219 | 9496 | 1.87 | [illegible] |
| 20 | 0.99 | KEGG_PATHWAY | Progesterone-mediated oocyte maturation | 4 | 138 | 111 | 9496 | 1.25 | [illegible] |
| 20 | 0.99 | KEGG_PATHWAY | Vascular smooth muscle contraction | 4 | 138 | 134 | 9496 | 1.25 | [illegible] |
| 20 | 0.99 | KEGG_PATHWAY | Retrograde endocannabinoid signaling | 4 | 138 | 149 | 9496 | 1.25 | [illegible] |
| 20 | 0.99 | KEGG_PATHWAY | GABAergic synapse | 3 | 138 | 89 | 9496 | 0.93 | [illegible] |
| 20 | 0.99 | KEGG_PATHWAY | Human immunodeficiency virus 1 infection | 5 | 138 | 214 | 9496 | 1.56 | [illegible] |
| 20 | 0.99 | KEGG_PATHWAY | Morphine addiction | 3 | 138 | 91 | 9496 | 0.93 | [illegible] |
| 20 | 0.99 | KEGG_PATHWAY | Adrenergic signaling in cardiomyocytes | 4 | 138 | 154 | 9496 | 1.25 | [illegible] |
| 20 | 0.99 | KEGG_PATHWAY | Circadian entrainment | 3 | 138 | 97 | 9496 | 0.93 | [illegible] |
| 20 | 0.99 | KEGG_PATHWAY | Inflammatory mediator regulation of TRP channels | 3 | 138 | 99 | 9496 | 0.93 | [illegible] |
| 20 | 0.99 | KEGG_PATHWAY | Cholinergic synapse | 3 | 138 | 116 | 9496 | 0.93 | [illegible] |
| 20 | 0.99 | GOTERM_BP_DIRECT | adenylate cyclase-activating G protein-coupled receptor signaling pathway | 4 | 296 | 176 | 19512 | 1.25 | [illegible] |

| Cluster | Cluster Enrichment Score | Category | Term | Count | List Total | Pop Hits | Pop Total | % | [illegible] |
|---|---|---|---|---|---|---|---|---|---|
| 20 | 0.99 | KEGG_PATHWAY | Neuroactive ligand signaling | 4 | 138 | 199 | 9496 | 1.25 | [illegible] |
| 20 | 0.99 | KEGG_PATHWAY | Dopaminergic synapse | 3 | 138 | 133 | 9496 | 0.93 | [illegible] |
| 20 | 0.99 | KEGG_PATHWAY | Estrogen signaling pathway | 3 | 138 | 139 | 9496 | 0.93 | [illegible] |
| 20 | 0.99 | KEGG_PATHWAY | Oocyte meiosis | 3 | 138 | 139 | 9496 | 0.93 | [illegible] |
| 20 | 0.99 | KEGG_PATHWAY | Thermogenesis | 3 | 138 | 234 | 9496 | 0.93 | [illegible] |
| 21 | 0.93 | KEGG_PATHWAY | Th17 cell differentiation | 6 | 138 | 109 | 9496 | 1.87 | [illegible] |
| 21 | 0.93 | KEGG_PATHWAY | Th1 and Th2 cell differentiation | 5 | 138 | 93 | 9496 | 1.56 | [illegible] |
| 21 | 0.93 | KEGG_PATHWAY | PD-L1 expression and PD-1 checkpoint pathway in cancer | 3 | 138 | 90 | 9496 | 0.93 | [illegible] |
| 21 | 0.93 | KEGG_PATHWAY | T cell receptor signaling pathway | 3 | 138 | 122 | 9496 | 0.93 | [illegible] |
| 22 | 0.92 | KEGG_PATHWAY | Th1 and Th2 cell differentiation | 5 | 138 | 93 | 9496 | 1.56 | [illegible] |
| 22 | 0.92 | KEGG_PATHWAY | C-type lectin receptor signaling pathway | 4 | 138 | 105 | 9496 | 1.25 | [illegible] |
| 22 | 0.92 | KEGG_PATHWAY | Lipid and atherosclerosis | 6 | 138 | 216 | 9496 | 1.87 | [illegible] |
| 23 | 0.90 | GOTERM_CC_DIRECT | endoplasmic reticulum | 34 | 308 | 1760 | 20808 | 10.59 | [illegible] |
| 23 | 0.90 | GOTERM_CC_DIRECT | endoplasmic reticulum membrane | 24 | 308 | 1228 | 20808 | 7.48 | [illegible] |
| 23 | 0.90 | UP_KW_CELLULAR_COMPONENT | Endoplasmic reticulum | 29 | 289 | 1439 | 18195 | 9.03 | [illegible] |
| 24 | 0.90 | UP_SEQ_FEATURE | DOMAIN:Calponin-homology (CH) | 4 | 312 | 79 | 20669 | 1.25 | [illegible] |
| 24 | 0.90 | INTERPRO | CH_dom | 4 | 314 | 81 | 20827 | 1.25 | [illegible] |
| 24 | 0.90 | INTERPRO | CH_dom_sf | 4 | 314 | 83 | 20827 | 1.25 | [illegible] |
| 24 | 0.90 | SMART | CH | 4 | 212 | 66 | 10704 | 1.25 | [illegible] |
| 25 | 0.89 | KEGG_PATHWAY | Toxoplasmosis | 6 | 138 | 112 | 9496 | 1.87 | [illegible] |
| 25 | 0.89 | KEGG_PATHWAY | Chagas disease | 5 | 138 | 103 | 9496 | 1.56 | [illegible] |
| 25 | 0.89 | KEGG_PATHWAY | Pertussis | 4 | 138 | 78 | 9496 | 1.25 | [illegible] |
| 25 | 0.89 | KEGG_PATHWAY | Leishmaniasis | 4 | 138 | 79 | 9496 | 1.25 | [illegible] |
| 25 | 0.89 | GOTERM_BP_DIRECT | cellular response to virus | 3 | 296 | 96 | 19512 | 0.93 | [illegible] |
| 25 | 0.89 | KEGG_PATHWAY | Tuberculosis | 3 | 138 | 182 | 9496 | 0.93 | [illegible] |

| Cluster | Cluster Enrichment Score | Category | Term | Count | List Total | Pop Hits | Pop Total | % | [illegible] |
|---|---|---|---|---|---|---|---|---|---|
| 26 | 0.84 | KEGG_PATHWAY | Cytokine-cytokine receptor interaction | 11 | 138 | 298 | 9496 | 3.43 | [illegible] |
| 26 | 0.84 | INTERPRO | TGF-beta-rel | 3 | 314 | 32 | 20827 | 0.93 | [illegible] |
| 26 | 0.84 | INTERPRO | TGF-b_C | 3 | 314 | 37 | 20827 | 0.93 | [illegible] |
| 26 | 0.84 | SMART | TGFB | 3 | 212 | 36 | 10704 | 0.93 | [illegible] |
| 26 | 0.84 | GOTERM_MF_DIRECT | cytokine activity | 7 | 306 | 240 | 19272 | 2.18 | [illegible] |
| 26 | 0.84 | UP_KW_MOLECULAR_FUNCTION | Cytokine | 6 | 187 | 195 | 12059 | 1.87 | [illegible] |
| 26 | 0.84 | GOTERM_MF_DIRECT | growth factor activity | 5 | 306 | 164 | 19272 | 1.56 | [illegible] |
| 26 | 0.84 | UP_SEQ_FEATURE | DISULFID:Interchain | 5 | 312 | 180 | 20669 | 1.56 | [illegible] |
| 26 | 0.84 | UP_KW_MOLECULAR_FUNCTION | Growth factor | 4 | 187 | 132 | 12059 | 1.25 | [illegible] |
| 26 | 0.84 | INTERPRO | Cystine-knot_cytokine | 3 | 314 | 80 | 20827 | 0.93 | [illegible] |
| 27 | 0.84 | GOTERM_MF_DIRECT | ATP hydrolysis activity | 13 | 306 | 440 | 19272 | 4.05 | [illegible] |
| 27 | 0.84 | INTERPRO | AAA+_ATPase | 5 | 314 | 141 | 20827 | 1.56 | [illegible] |
| 27 | 0.84 | UP_SEQ_FEATURE | DOMAIN:AAA+ ATPase | 3 | 312 | 53 | 20669 | 0.93 | [illegible] |
| 27 | 0.84 | SMART | AAA | 5 | 212 | 141 | 10704 | 1.56 | [illegible] |
| 28 | 0.82 | GOTERM_MF_DIRECT | ATPase-coupled intramembrane lipid transporter activity | 3 | 306 | 21 | 19272 | 0.93 | [illegible] |
| 28 | 0.82 | GOTERM_BP_DIRECT | phospholipid translocation | 3 | 296 | 27 | 19512 | 0.93 | [illegible] |
| 28 | 0.82 | GOTERM_BP_DIRECT | phospholipid transport | 3 | 296 | 48 | 19512 | 0.93 | [illegible] |
| 28 | 0.82 | UP_KW_MOLECULAR_FUNCTION | Translocase | 4 | 187 | 107 | 12059 | 1.25 | [illegible] |
| 28 | 0.82 | UP_KW_BIOLOGICAL_PROCESS | Lipid transport | 5 | 193 | 177 | 11605 | 1.56 | [illegible] |
| 28 | 0.82 | GOTERM_BP_DIRECT | lipid transport | 5 | 296 | 193 | 19512 | 1.56 | [illegible] |
| 29 | 0.79 | UP_SEQ_FEATURE | DOMAIN:BTB | 6 | 312 | 170 | 20669 | 1.87 | [illegible] |
| 29 | 0.79 | INTERPRO | BTB/POZ_dom | 6 | 314 | 180 | 20827 | 1.87 | [illegible] |
| 29 | 0.79 | INTERPRO | SKP1/BTB/POZ_sf | 6 | 314 | 190 | 20827 | 1.87 | [illegible] |
| 29 | 0.79 | SMART | BTB | 6 | 212 | 176 | 10704 | 1.87 | [illegible] |
| 30 | 0.79 | KEGG_PATHWAY | Hippo signaling pathway | 7 | 138 | 157 | 9496 | 2.18 | [illegible] |
| 30 | 0.79 | GOTERM_BP_DIRECT | response to xenobiotic stimulus | 8 | 296 | 260 | 19512 | 2.49 | [illegible] |
| 30 | 0.79 | KEGG_PATHWAY | Gastric cancer | 4 | 138 | 150 | 9496 | 1.25 | [illegible] |

| Cluster | Cluster Enrichment Score | Category | Term | Count | List Total | Pop Hits | Pop Total | % | [illegible] |
|---|---|---|---|---|---|---|---|---|---|
| 30 | 0.79 | KEGG_PATHWAY | Hepatocellular carcinoma | 3 | 138 | 170 | 9496 | 0.93 | [illegible] |
| 31 | 0.79 | KEGG_PATHWAY | FoxO signaling pathway | 5 | 138 | 133 | 9496 | 1.56 | [illegible] |
| 31 | 0.79 | KEGG_PATHWAY | AGE-RAGE signaling pathway in diabetic complications | 4 | 138 | 101 | 9496 | 1.25 | [illegible] |
| 31 | 0.79 | KEGG_PATHWAY | Cellular senescence | 5 | 138 | 157 | 9496 | 1.56 | [illegible] |
| 32 | 0.67 | INTERPRO | SAM | 4 | 314 | 95 | 20827 | 1.25 | [illegible] |
| 32 | 0.67 | UP_SEQ_FEATURE | DOMAIN:SAM | 4 | 312 | 95 | 20669 | 1.25 | [illegible] |
| 32 | 0.67 | SMART | SAM | 4 | 212 | 86 | 10704 | 1.25 | [illegible] |
| 32 | 0.67 | INTERPRO | SAM/pointed_sf | 4 | 314 | 126 | 20827 | 1.25 | [illegible] |
| 33 | 0.66 | KEGG_PATHWAY | Chagas disease | 5 | 138 | 103 | 9496 | 1.56 | [illegible] |
| 33 | 0.66 | KEGG_PATHWAY | Lipid and atherosclerosis | 6 | 138 | 216 | 9496 | 1.87 | [illegible] |
| 33 | 0.66 | KEGG_PATHWAY | IL-17 signaling pathway | 3 | 138 | 95 | 9496 | 0.93 | [illegible] |
| 33 | 0.66 | KEGG_PATHWAY | Coronavirus disease - COVID-19 | 5 | 138 | 238 | 9496 | 1.56 | [illegible] |
| 34 | 0.62 | UP_SEQ_FEATURE | SITE:Reactive bond | 3 | 312 | 47 | 20669 | 0.93 | [illegible] |
| 34 | 0.62 | UP_KW_MOLECULAR_FUNCTION | Serine protease inhibitor | 4 | 187 | 93 | 12059 | 1.25 | [illegible] |
| 34 | 0.62 | GOTERM_MF_DIRECT | serine-type endopeptidase inhibitor activity | 4 | 306 | 109 | 19272 | 1.25 | [illegible] |
| 34 | 0.62 | UP_KW_MOLECULAR_FUNCTION | Protease inhibitor | 4 | 187 | 133 | 12059 | 1.25 | [illegible] |
| 34 | 0.62 | GOTERM_MF_DIRECT | peptidase inhibitor activity | 4 | 306 | 131 | 19272 | 1.25 | [illegible] |
| 35 | 0.62 | GOTERM_CC_DIRECT | side of membrane | 5 | 308 | 150 | 20808 | 1.56 | [illegible] |
| 35 | 0.62 | UP_SEQ_FEATURE | LIPID:GPI-anchor amidated serine | 3 | 312 | 59 | 20669 | 0.93 | [illegible] |
| 35 | 0.62 | UP_KW_PTM | GPI-anchor | 5 | 258 | 151 | 14452 | 1.56 | [illegible] |
| 35 | 0.62 | UP_SEQ_FEATURE | PROPEP:Removed in mature form | 7 | 312 | 297 | 20669 | 2.18 | [illegible] |
| 36 | 0.61 | UP_SEQ_FEATURE | DOMAIN:PDZ | 5 | 312 | 152 | 20669 | 1.56 | [illegible] |
| 36 | 0.61 | INTERPRO | PDZ | 5 | 314 | 159 | 20827 | 1.56 | [illegible] |
| 36 | 0.61 | INTERPRO | PDZ_sf | 5 | 314 | 162 | 20827 | 1.56 | [illegible] |
| 36 | 0.61 | SMART | PDZ | 5 | 212 | 156 | 10704 | 1.56 | [illegible] |
| 37 | 0.60 | INTERPRO | Ig_I-set | 7 | 314 | 143 | 20827 | 2.18 | [illegible] |

| Cluster | Cluster Enrichment Score | Category | Term | Count | List Total | Pop Hits | Pop Total | % | [illegible] |
|---|---|---|---|---|---|---|---|---|---|
| 37 | 0.60 | UP_SEQ_FEATURE | DOMAIN:Ig-like C2-type 5 | 4 | 312 | 57 | 20669 | 1.25 | [illegible] |
| 37 | 0.60 | INTERPRO | FN3_dom | 7 | 314 | 209 | 20827 | 2.18 | [illegible] |
| 37 | 0.60 | INTERPRO | FN3_sf | 7 | 314 | 211 | 20827 | 2.18 | [illegible] |
| 37 | 0.60 | UP_SEQ_FEATURE | DOMAIN:Fibronectin type-III | 6 | 312 | 174 | 20669 | 1.87 | [illegible] |
| 37 | 0.60 | UP_SEQ_FEATURE | DOMAIN:Ig-like C2-type 4 | 4 | 312 | 83 | 20669 | 1.25 | [illegible] |
| 37 | 0.60 | UP_SEQ_FEATURE | DOMAIN:Fibronectin type-III 3 | 4 | 312 | 86 | 20669 | 1.25 | [illegible] |
| 37 | 0.60 | UP_SEQ_FEATURE | DOMAIN:Ig-like C2-type 3 | 5 | 312 | 133 | 20669 | 1.56 | [illegible] |
| 37 | 0.60 | INTERPRO | Ig_sub2 | 7 | 314 | 262 | 20827 | 2.18 | [illegible] |
| 37 | 0.60 | UP_SEQ_FEATURE | DOMAIN:Ig-like C2-type 1 | 6 | 312 | 219 | 20669 | 1.87 | [illegible] |
| 37 | 0.60 | UP_SEQ_FEATURE | DOMAIN:Ig-like C2-type 2 | 6 | 312 | 219 | 20669 | 1.87 | [illegible] |
| 37 | 0.60 | SMART | FN3 | 5 | 212 | 150 | 10704 | 1.56 | [illegible] |
| 37 | 0.60 | UP_SEQ_FEATURE | DOMAIN:Fibronectin type-III 1 | 4 | 312 | 139 | 20669 | 1.25 | [illegible] |
| 37 | 0.60 | UP_SEQ_FEATURE | DOMAIN:Fibronectin type-III 2 | 4 | 312 | 139 | 20669 | 1.25 | [illegible] |
| 37 | 0.60 | INTERPRO | Ig_sub | 10 | 314 | 528 | 20827 | 3.12 | [illegible] |
| 37 | 0.60 | SMART | IGc2 | 7 | 212 | 262 | 10704 | 2.18 | [illegible] |
| 37 | 0.60 | INTERPRO | Ig-like_fold | 19 | 314 | 1123 | 20827 | 5.92 | [illegible] |
| 37 | 0.60 | SMART | IG | 10 | 212 | 528 | 10704 | 3.12 | [illegible] |
| 37 | 0.60 | UP_SEQ_FEATURE | DOMAIN:Ig-like | 8 | 312 | 676 | 20669 | 2.49 | [illegible] |
| 37 | 0.60 | INTERPRO | Ig-like_dom | 9 | 314 | 768 | 20827 | 2.80 | [illegible] |
| 37 | 0.60 | INTERPRO | Ig-like_dom_sf | 10 | 314 | 869 | 20827 | 3.12 | [illegible] |
| 37 | 0.60 | UP_KW_DOMAIN | Immunoglobulin domain | 10 | 237 | 828 | 14656 | 3.12 | [illegible] |
| 38 | 0.60 | GOTERM_CC_DIRECT | recycling endosome | 5 | 308 | 157 | 20808 | 1.56 | [illegible] |
| 38 | 0.60 | GOTERM_CC_DIRECT | recycling endosome membrane | 4 | 308 | 116 | 20808 | 1.25 | [illegible] |
| 38 | 0.60 | GOTERM_CC_DIRECT | trans-Golgi network | 5 | 308 | 194 | 20808 | 1.56 | [illegible] |
| 39 | 0.59 | UP_KW_MOLECULAR_FUNCTION | Motor protein | 5 | 187 | 138 | 12059 | 1.56 | [illegible] |
| 39 | 0.59 | GOTERM_CC_DIRECT | kinesin complex | 3 | 308 | 50 | 20808 | 0.93 | [illegible] |
| 39 | 0.59 | UP_KW_CELLULAR_COMPONENT | Microtubule | 8 | 289 | 302 | 18195 | 2.49 | [illegible] |

| Cluster | Cluster Enrichment Score | Category | Term | Count | List Total | Pop Hits | Pop Total | % | [illegible] |
|---|---|---|---|---|---|---|---|---|---|
| 39 | 0.59 | GOTERM_CC_DIRECT | microtubule | 8 | 308 | 377 | 20808 | 2.49 | [illegible] |
| 39 | 0.59 | GOTERM_MF_DIRECT | microtubule binding | 5 | 306 | 277 | 19272 | 1.56 | [illegible] |
| 40 | 0.57 | INTERPRO | TPR_rpt | 5 | 314 | 140 | 20827 | 1.56 | [illegible] |
| 40 | 0.57 | UP_SEQ_FEATURE | REPEAT:TPR 3 | 5 | 312 | 146 | 20669 | 1.56 | [illegible] |
| 40 | 0.57 | UP_SEQ_FEATURE | REPEAT:TPR 4 | 4 | 312 | 101 | 20669 | 1.25 | [illegible] |
| 40 | 0.57 | UP_SEQ_FEATURE | REPEAT:TPR 1 | 5 | 312 | 160 | 20669 | 1.56 | [illegible] |
| 40 | 0.57 | UP_SEQ_FEATURE | REPEAT:TPR 2 | 5 | 312 | 160 | 20669 | 1.56 | [illegible] |
| 40 | 0.57 | SMART | TPR | 5 | 212 | 132 | 10704 | 1.56 | [illegible] |
| 40 | 0.57 | UP_KW_DOMAIN | TPR repeat | 5 | 237 | 171 | 14656 | 1.56 | [illegible] |
| 40 | 0.57 | UP_SEQ_FEATURE | REPEAT:TPR 6 | 3 | 312 | 77 | 20669 | 0.93 | [illegible] |
| 40 | 0.57 | UP_SEQ_FEATURE | REPEAT:TPR 5 | 3 | 312 | 82 | 20669 | 0.93 | [illegible] |
| 40 | 0.57 | UP_SEQ_FEATURE | REPEAT:TPR | 3 | 312 | 99 | 20669 | 0.93 | [illegible] |
| 40 | 0.57 | INTERPRO | TPR-like_helical_dom_sf | 5 | 314 | 230 | 20827 | 1.56 | [illegible] |
| 41 | 0.56 | GOTERM_MF_DIRECT | transmembrane transporter activity | 8 | 306 | 231 | 19272 | 2.49 | [illegible] |
| 41 | 0.56 | INTERPRO | MFS_dom | 3 | 314 | 99 | 20827 | 0.93 | [illegible] |
| 41 | 0.56 | INTERPRO | MFS_trans_sf | 3 | 314 | 147 | 20827 | 0.93 | [illegible] |
| 42 | 0.50 | UP_KW_MOLECULAR_FUNCTION | Motor protein | 5 | 187 | 138 | 12059 | 1.56 | [illegible] |
| 42 | 0.50 | INTERPRO | Kinesin_motor_dom_sf | 3 | 314 | 84 | 20827 | 0.93 | [illegible] |
| 42 | 0.50 | KEGG_PATHWAY | Motor proteins | 4 | 138 | 197 | 9496 | 1.25 | [illegible] |
| 43 | 0.45 | KEGG_PATHWAY | Vascular smooth muscle contraction | 4 | 138 | 134 | 9496 | 1.25 | [illegible] |
| 43 | 0.45 | KEGG_PATHWAY | Insulin secretion | 3 | 138 | 86 | 9496 | 0.93 | [illegible] |
| 43 | 0.45 | KEGG_PATHWAY | Salivary secretion | 3 | 138 | 97 | 9496 | 0.93 | [illegible] |
| 44 | 0.38 | GOTERM_BP_DIRECT | positive regulation of transcription by RNA polymerase II | 28 | 296 | 1243 | 19512 | 8.72 | [illegible] |
| 44 | 0.38 | GOTERM_MF_DIRECT | DNA-binding transcription factor activity | 17 | 306 | 739 | 19272 | 5.30 | [illegible] |
| 44 | 0.38 | GOTERM_MF_DIRECT | DNA-binding transcription repressor activity, RNA polymerase II-specific | 8 | 306 | 293 | 19272 | 2.49 | [illegible] |
| 44 | 0.38 | GOTERM_MF_DIRECT | sequence-specific double- | 14 | 306 | 638 | 19272 | 4.36 | [illegible] |

| Cluster | Cluster Enrichment Score | Category | Term | Count | List Total | Pop Hits | Pop Total | % | [illegible] |
|---|---|---|---|---|---|---|---|---|---|
| | | | stranded DNA binding | | | | | | |
| 44 | 0.38 | UP_KW_MOLECULAR_FUNCTION | Activator | 15 | 187 | 733 | 12059 | 4.67 | [illegible] |
| 44 | 0.38 | GOTERM_MF_DIRECT | sequence-specific DNA binding | 11 | 306 | 514 | 19272 | 3.43 | [illegible] |
| 44 | 0.38 | GOTERM_MF_DIRECT | DNA-binding transcription activator activity, RNA polymerase II-specific | 10 | 306 | 456 | 19272 | 3.12 | [illegible] |
| 44 | 0.38 | GOTERM_BP_DIRECT | negative regulation of transcription by RNA polymerase II | 19 | 296 | 1034 | 19512 | 5.92 | [illegible] |
| 44 | 0.38 | GOTERM_MF_DIRECT | RNA polymerase II cis-regulatory region sequence-specific DNA binding | 18 | 306 | 1100 | 19272 | 5.61 | [illegible] |
| 44 | 0.38 | GOTERM_CC_DIRECT | chromatin | 17 | 308 | 1159 | 20808 | 5.30 | [illegible] |
| 44 | 0.38 | GOTERM_MF_DIRECT | DNA-binding transcription factor activity, RNA polymerase II-specific | 19 | 306 | 1247 | 19272 | 5.92 | [illegible] |
| 44 | 0.38 | GOTERM_BP_DIRECT | regulation of DNA-templated transcription | 22 | 296 | 1526 | 19512 | 6.85 | [illegible] |
| 44 | 0.38 | GOTERM_BP_DIRECT | regulation of transcription by RNA polymerase II | 22 | 296 | 1643 | 19512 | 6.85 | [illegible] |
| 44 | 0.38 | UP_KW_BIOLOGICAL_PROCESS | Transcription regulation | 34 | 193 | 2444 | 11605 | 10.59 | [illegible] |
| 44 | 0.38 | UP_KW_MOLECULAR_FUNCTION | DNA-binding | 27 | 187 | 2143 | 12059 | 8.41 | [illegible] |
| 44 | 0.38 | UP_KW_BIOLOGICAL_PROCESS | Transcription | 34 | 193 | 2516 | 11605 | 10.59 | [illegible] |
| 44 | 0.38 | GOTERM_MF_DIRECT | DNA binding | 29 | 306 | 2370 | 19272 | 9.03 | [illegible] |
| 44 | 0.38 | UP_KW_CELLULAR_COMPONENT | Nucleus | 71 | 289 | 5969 | 18195 | 22.12 | [illegible] |
| 45 | 0.38 | GOTERM_MF_DIRECT | endopeptidase activity | 5 | 306 | 96 | 19272 | 1.56 | [illegible] |
| 45 | 0.38 | UP_SEQ_FEATURE | SITE:Cleavage; by autolysis | 4 | 312 | 66 | 20669 | 1.25 | [illegible] |
| 45 | 0.38 | GOTERM_BP_DIRECT | protein processing | 3 | 296 | 96 | 19512 | 0.93 | [illegible] |
| 45 | 0.38 | GOTERM_MF_DIRECT | serine-type endopeptidase activity | 4 | 306 | 192 | 19272 | 1.25 | [illegible] |
| 45 | 0.38 | UP_KW_PTM | Zymogen | 4 | 258 | 227 | 14452 | 1.25 | [illegible] |
| 45 | 0.38 | GOTERM_MF_DIRECT | peptidase activity | 5 | 306 | 544 | 19272 | 1.56 | [illegible] |
| 45 | 0.38 | UP_KW_MOLECULAR_FUNCTION | Protease | 5 | 187 | 586 | 12059 | 1.56 | [illegible] |

| Cluster | Cluster Enrichment Score | Category | Term | Count | List Total | Pop Hits | Pop Total | % | [illegible] |
|---|---|---|---|---|---|---|---|---|---|
| 45 | 0.38 | GOTERM_BP_DIRECT | proteolysis | 5 | 296 | 671 | 19512 | 1.56 | [illegible] |
| 46 | 0.37 | GOTERM_CC_DIRECT | voltage-gated potassium channel complex | 4 | 308 | 88 | 20808 | 1.25 | [illegible] |
| 46 | 0.37 | UP_KW_BIOLOGICAL_PROCESS | Potassium transport | 5 | 193 | 129 | 11605 | 1.56 | [illegible] |
| 46 | 0.37 | GOTERM_BP_DIRECT | potassium ion transport | 5 | 296 | 155 | 19512 | 1.56 | [illegible] |
| 46 | 0.37 | UP_KW_MOLECULAR_FUNCTION | Potassium channel | 3 | 187 | 76 | 12059 | 0.93 | [illegible] |
| 46 | 0.37 | UP_KW_LIGAND | Potassium | 5 | 137 | 151 | 7011 | 1.56 | [illegible] |
| 46 | 0.37 | GOTERM_BP_DIRECT | potassium ion transmembrane transport | 4 | 296 | 151 | 19512 | 1.25 | [illegible] |
| 46 | 0.37 | GOTERM_MF_DIRECT | potassium channel activity | 3 | 306 | 88 | 19272 | 0.93 | [illegible] |
| 46 | 0.37 | GOTERM_BP_DIRECT | monoatomic ion transport | 11 | 296 | 672 | 19512 | 3.43 | [illegible] |
| 46 | 0.37 | UP_KW_MOLECULAR_FUNCTION | Voltage-gated channel | 3 | 187 | 141 | 12059 | 0.93 | [illegible] |
| 46 | 0.37 | GOTERM_CC_DIRECT | monoatomic ion channel complex | 3 | 308 | 150 | 20808 | 0.93 | [illegible] |
| 46 | 0.37 | UP_KW_BIOLOGICAL_PROCESS | Ion transport | 9 | 193 | 647 | 11605 | 2.80 | [illegible] |
| 46 | 0.37 | GOTERM_BP_DIRECT | monoatomic ion transmembrane transport | 5 | 296 | 413 | 19512 | 1.56 | [illegible] |
| 46 | 0.37 | UP_KW_MOLECULAR_FUNCTION | Ion channel | 3 | 187 | 356 | 12059 | 0.93 | [illegible] |
| 47 | 0.35 | GOTERM_MF_DIRECT | carboxylic ester hydrolase activity | 3 | 306 | 60 | 19272 | 0.93 | [illegible] |
| 47 | 0.35 | UP_KW_BIOLOGICAL_PROCESS | Lipid degradation | 3 | 193 | 103 | 11605 | 0.93 | [illegible] |
| 47 | 0.35 | INTERPRO | AB_hydrolase_fold | 3 | 314 | 125 | 20827 | 0.93 | [illegible] |
| 47 | 0.35 | GOTERM_BP_DIRECT | lipid catabolic process | 3 | 296 | 130 | 19512 | 0.93 | [illegible] |
| 48 | 0.34 | GOTERM_BP_DIRECT | sodium ion transport | 5 | 296 | 146 | 19512 | 1.56 | [illegible] |
| 48 | 0.34 | UP_KW_BIOLOGICAL_PROCESS | Sodium transport | 3 | 193 | 131 | 11605 | 0.93 | [illegible] |
| 48 | 0.34 | UP_KW_LIGAND | Sodium | 3 | 137 | 162 | 7011 | 0.93 | [illegible] |
| 49 | 0.33 | UP_SEQ_FEATURE | DOMAIN:C2 1 | 3 | 312 | 61 | 20669 | 0.93 | [illegible] |
| 49 | 0.33 | UP_SEQ_FEATURE | DOMAIN:C2 2 | 3 | 312 | 62 | 20669 | 0.93 | [illegible] |
| 49 | 0.33 | INTERPRO | C2_domain_sf | 4 | 314 | 177 | 20827 | 1.25 | [illegible] |
| 49 | 0.33 | INTERPRO | C2_dom | 3 | 314 | 148 | 20827 | 0.93 | [illegible] |
| 49 | 0.33 | SMART | C2 | 3 | 212 | 126 | 10704 | 0.93 | [illegible] |
| 49 | 0.33 | GOTERM_CC_DIRECT | cytoplasmic vesicle | 3 | 308 | 195 | 20808 | 0.93 | [illegible] |

| Cluster | Cluster Enrichment Score | Category | Term | Count | List Total | Pop Hits | Pop Total | % | [illegible] |
|---|---|---|---|---|---|---|---|---|---|
| | | | membrane | | | | | | |
| 50 | 0.31 | GOTERM_BP_DIRECT | cell division | 9 | 296 | 433 | 19512 | 2.80 | [illegible] |
| 50 | 0.31 | UP_KW_BIOLOGICAL_PROCESS | Cell division | 9 | 193 | 422 | 11605 | 2.80 | [illegible] |
| 50 | 0.31 | UP_KW_BIOLOGICAL_PROCESS | Cell cycle | 12 | 193 | 692 | 11605 | 3.74 | [illegible] |
| 50 | 0.31 | UP_KW_BIOLOGICAL_PROCESS | Mitosis | 5 | 193 | 294 | 11605 | 1.56 | [illegible] |
| 51 | 0.29 | GOTERM_CC_DIRECT | extracellular space | 31 | 308 | 1895 | 20808 | 9.66 | [illegible] |
| 51 | 0.29 | GOTERM_CC_DIRECT | extracellular region | 41 | 308 | 2693 | 20808 | 12.77 | [illegible] |
| 51 | 0.29 | UP_KW_CELLULAR_COMPONENT | Secreted | 34 | 289 | 2233 | 18195 | 10.59 | [illegible] |
| 52 | 0.18 | INTERPRO | Small_GTPase | 4 | 314 | 144 | 20827 | 1.25 | [illegible] |
| 52 | 0.18 | GOTERM_MF_DIRECT | GTP binding | 8 | 306 | 407 | 19272 | 2.49 | [illegible] |
| 52 | 0.18 | SMART | RHO | 3 | 212 | 132 | 10704 | 0.93 | [illegible] |
| 52 | 0.18 | INTERPRO | Small_GTP-bd | 3 | 314 | 176 | 20827 | 0.93 | [illegible] |
| 52 | 0.18 | SMART | RAS | 3 | 212 | 143 | 10704 | 0.93 | [illegible] |
| 52 | 0.18 | SMART | RAB | 3 | 212 | 159 | 10704 | 0.93 | [illegible] |
| 52 | 0.18 | UP_KW_LIGAND | GTP-binding | 6 | 137 | 375 | 7011 | 1.87 | [illegible] |
| 53 | 0.15 | UP_SEQ_FEATURE | REPEAT:3 | 4 | 312 | 225 | 20669 | 1.25 | [illegible] |
| 53 | 0.15 | UP_SEQ_FEATURE | REPEAT:1 | 4 | 312 | 250 | 20669 | 1.25 | [illegible] |
| 53 | 0.15 | UP_SEQ_FEATURE | REPEAT:2 | 4 | 312 | 253 | 20669 | 1.25 | [illegible] |
| 54 | 0.13 | UP_SEQ_FEATURE | REPEAT:ANK 6 | 3 | 312 | 112 | 20669 | 0.93 | [illegible] |
| 54 | 0.13 | UP_SEQ_FEATURE | REPEAT:ANK 3 | 4 | 312 | 222 | 20669 | 1.25 | [illegible] |
| 54 | 0.13 | UP_SEQ_FEATURE | REPEAT:ANK 5 | 3 | 312 | 158 | 20669 | 0.93 | [illegible] |
| 54 | 0.13 | UP_SEQ_FEATURE | REPEAT:ANK 1 | 4 | 312 | 260 | 20669 | 1.25 | [illegible] |
| 54 | 0.13 | UP_SEQ_FEATURE | REPEAT:ANK 2 | 4 | 312 | 261 | 20669 | 1.25 | [illegible] |
| 54 | 0.13 | INTERPRO | Ankyrin_rpt | 4 | 314 | 269 | 20827 | 1.25 | [illegible] |
| 54 | 0.13 | UP_SEQ_FEATURE | REPEAT:ANK 4 | 3 | 312 | 188 | 20669 | 0.93 | [illegible] |
| 54 | 0.13 | INTERPRO | Ankyrin_rpt-contain_sf | 4 | 314 | 275 | 20827 | 1.25 | [illegible] |
| 54 | 0.13 | UP_SEQ_FEATURE | REPEAT:ANK | 3 | 312 | 198 | 20669 | 0.93 | [illegible] |
| 54 | 0.13 | UP_KW_DOMAIN | ANK repeat | 4 | 237 | 267 | 14656 | 1.25 | [illegible] |
| 54 | 0.13 | SMART | ANK | 4 | 212 | 258 | 10704 | 1.25 | [illegible] |

| Cluster | Cluster Enrichment Score | Category | Term | Count | List Total | Pop Hits | Pop Total | % | [illegible] |
|---|---|---|---|---|---|---|---|---|---|
| 55 | 0.13 | UP_KW_BIOLOGICAL_PROCESS | Vision | 3 | 193 | 127 | 11605 | 0.93 | [illegible] |
| 55 | 0.13 | GOTERM_BP_DIRECT | visual perception | 4 | 296 | 221 | 19512 | 1.25 | [illegible] |
| 55 | 0.13 | UP_KW_BIOLOGICAL_PROCESS | Sensory transduction | 3 | 193 | 634 | 11605 | 0.93 | [illegible] |
| 56 | 0.10 | GOTERM_MF_DIRECT | transcription cis-regulatory region binding | 6 | 306 | 286 | 19272 | 1.87 | [illegible] |
| 56 | 0.10 | UP_SEQ_FEATURE | DOMAIN:C2H2-type | 9 | 312 | 521 | 20669 | 2.80 | [illegible] |
| 56 | 0.10 | UP_SEQ_FEATURE | ZN_FING:C2H2-type 1 | 9 | 312 | 560 | 20669 | 2.80 | [illegible] |
| 56 | 0.10 | UP_SEQ_FEATURE | ZN_FING:C2H2-type 2 | 9 | 312 | 633 | 20669 | 2.80 | [illegible] |
| 56 | 0.10 | UP_SEQ_FEATURE | ZN_FING:C2H2-type 3 | 9 | 312 | 655 | 20669 | 2.80 | [illegible] |
| 56 | 0.10 | GOTERM_BP_DIRECT | regulation of transcription by RNA polymerase II | 22 | 296 | 1643 | 19512 | 6.85 | [illegible] |
| 56 | 0.10 | INTERPRO | Znf_C2H2_sf | 9 | 314 | 796 | 20827 | 2.80 | [illegible] |
| 56 | 0.10 | INTERPRO | Znf_C2H2_type | 9 | 314 | 828 | 20827 | 2.80 | [illegible] |
| 56 | 0.10 | UP_SEQ_FEATURE | ZN_FING:C2H2-type 4 | 5 | 312 | 608 | 20669 | 1.56 | [illegible] |
| 56 | 0.10 | SMART | ZnF_C2H2 | 9 | 212 | 776 | 10704 | 2.80 | [illegible] |
| 56 | 0.10 | UP_KW_DOMAIN | Zinc-finger | 17 | 237 | 1842 | 14656 | 5.30 | [illegible] |
| 56 | 0.10 | GOTERM_MF_DIRECT | zinc ion binding | 21 | 306 | 2237 | 19272 | 6.54 | [illegible] |
| 57 | 0.10 | GOTERM_BP_DIRECT | proteasome-mediated ubiquitin-dependent protein catabolic process | 7 | 296 | 356 | 19512 | 2.18 | [illegible] |
| 57 | 0.10 | INTERPRO | Znf_RING_CS | 3 | 314 | 183 | 20827 | 0.93 | [illegible] |
| 57 | 0.10 | UP_SEQ_FEATURE | DOMAIN:RING-type | 3 | 312 | 210 | 20669 | 0.93 | [illegible] |
| 57 | 0.10 | UP_SEQ_FEATURE | ZN_FING:RING-type | 3 | 312 | 219 | 20669 | 0.93 | [illegible] |
| 57 | 0.10 | INTERPRO | Znf_RING | 4 | 314 | 314 | 20827 | 1.25 | [illegible] |
| 57 | 0.10 | SMART | RING | 4 | 212 | 259 | 10704 | 1.25 | [illegible] |
| 57 | 0.10 | GOTERM_MF_DIRECT | ubiquitin protein ligase activity | 4 | 306 | 383 | 19272 | 1.25 | [illegible] |
| 57 | 0.10 | INTERPRO | Znf_RING/FYVE/PHD | 4 | 314 | 491 | 20827 | 1.25 | [illegible] |
| 58 | 0.09 | UP_SEQ_FEATURE | DOMAIN:Homeobox | 3 | 312 | 130 | 20669 | 0.93 | [illegible] |
| 58 | 0.09 | INTERPRO | Homeodomain-like_sf | 5 | 314 | 328 | 20827 | 1.56 | [illegible] |
| 58 | 0.09 | UP_SEQ_FEATURE | DNA_BIND:Homeobox | 3 | 312 | 234 | 20669 | 0.93 | [illegible] |
| 58 | 0.09 | INTERPRO | HD | 3 | 314 | 258 | 20827 | 0.93 | [illegible] |

| Cluster | Cluster Enrichment Score | Category | Term | Count | List Total | Pop Hits | Pop Total | % | [illegible] |
|---|---|---|---|---|---|---|---|---|---|
| 58 | 0.09 | UP_KW_DOMAIN | Homeobox | 3 | 237 | 256 | 14656 | 0.93 | [illegible] |
| 58 | 0.09 | SMART | HOX | 3 | 212 | 252 | 10704 | 0.93 | [illegible] |
| 59 | 0.09 | GOTERM_CC_DIRECT | lysosomal membrane | 7 | 308 | 418 | 20808 | 2.18 | [illegible] |
| 59 | 0.09 | GOTERM_CC_DIRECT | lysosome | 5 | 308 | 532 | 20808 | 1.56 | [illegible] |
| 59 | 0.09 | UP_KW_CELLULAR_COMPONENT | Lysosome | 4 | 289 | 450 | 18195 | 1.25 | [illegible] |
| 60 | 0.09 | GOTERM_MF_DIRECT | metal ion binding | 66 | 306 | 4143 | 19272 | 20.56 | [illegible] |
| 60 | 0.09 | UP_KW_LIGAND | Metal-binding | 64 | 137 | 4245 | 7011 | 19.94 | [illegible] |
| 60 | 0.09 | UP_KW_LIGAND | Zinc | 27 | 137 | 2593 | 7011 | 8.41 | [illegible] |
| 61 | 0.08 | GOTERM_CC_DIRECT | sperm flagellum | 3 | 308 | 161 | 20808 | 0.93 | [illegible] |
| 61 | 0.08 | UP_KW_CELLULAR_COMPONENT | Flagellum | 3 | 289 | 229 | 18195 | 0.93 | [illegible] |
| 61 | 0.08 | UP_KW_CELLULAR_COMPONENT | Cilium | 4 | 289 | 393 | 18195 | 1.25 | [illegible] |
| 62 | 0.06 | UP_SEQ_FEATURE | DOMAIN:EF-hand | 3 | 312 | 199 | 20669 | 0.93 | [illegible] |
| 62 | 0.06 | INTERPRO | EF_hand_dom | 3 | 314 | 233 | 20827 | 0.93 | [illegible] |
| 62 | 0.06 | INTERPRO | EF-hand-dom_pair | 3 | 314 | 276 | 20827 | 0.93 | [illegible] |
| 63 | 0.04 | GOTERM_CC_DIRECT | nucleoplasm | 55 | 308 | 4091 | 20808 | 17.13 | [illegible] |
| 63 | 0.04 | GOTERM_CC_DIRECT | nucleus | 96 | 308 | 7054 | 20808 | 29.91 | [illegible] |
| 63 | 0.04 | UP_KW_CELLULAR_COMPONENT | Nucleus | 71 | 289 | 5969 | 18195 | 22.12 | [illegible] |
| 64 | 0.04 | GOTERM_BP_DIRECT | intracellular protein transport | 4 | 296 | 294 | 19512 | 1.25 | [illegible] |
| 64 | 0.04 | GOTERM_BP_DIRECT | protein transport | 7 | 296 | 703 | 19512 | 2.18 | [illegible] |
| 64 | 0.04 | UP_KW_BIOLOGICAL_PROCESS | Protein transport | 6 | 193 | 661 | 11605 | 1.87 | [illegible] |
| 65 | 0.03 | UP_KW_BIOLOGICAL_PROCESS | Innate immunity | 6 | 193 | 432 | 11605 | 1.87 | [illegible] |
| 65 | 0.03 | GOTERM_BP_DIRECT | innate immune response | 7 | 296 | 621 | 19512 | 2.18 | [illegible] |
| 65 | 0.03 | GOTERM_BP_DIRECT | immune system process | 10 | 296 | 953 | 19512 | 3.12 | [illegible] |
| 65 | 0.03 | UP_KW_BIOLOGICAL_PROCESS | Immunity | 9 | 193 | 988 | 11605 | 2.80 | [illegible] |
| 66 | 0.00 | UP_SEQ_FEATURE | ZN_FING:C2H2-type 4 | 5 | 312 | 608 | 20669 | 1.56 | [illegible] |
| 66 | 0.00 | UP_SEQ_FEATURE | ZN_FING:C2H2-type 6 | 3 | 312 | 521 | 20669 | 0.93 | [illegible] |
| 66 | 0.00 | UP_SEQ_FEATURE | ZN_FING:C2H2-type 5 | 3 | 312 | 572 | 20669 | 0.93 | [illegible] |
| 67 | 0.00 | UP_SEQ_FEATURE | TRANSMEM:Helical; Name=7 | 7 | 312 | 968 | 20669 | 2.18 | [illegible] |

| Cluster | Cluster Enrichment Score | Category | Term | Count | List Total | Pop Hits | Pop Total | % | [illegible] |
|---|---|---|---|---|---|---|---|---|---|
| 67 | 0.00 | UP_SEQ_FEATURE | TRANSMEM:Helical; Name=6 | 7 | 312 | 1046 | 20669 | 2.18 | [illegible] |
| 67 | 0.00 | UP_SEQ_FEATURE | TRANSMEM:Helical; Name=5 | 7 | 312 | 1056 | 20669 | 2.18 | [illegible] |
| 67 | 0.00 | UP_KW_MOLECULAR_FUNCTION | Transducer | 6 | 187 | 927 | 12059 | 1.87 | [illegible] |
| 67 | 0.00 | UP_SEQ_FEATURE | TRANSMEM:Helical; Name=3 | 7 | 312 | 1082 | 20669 | 2.18 | [illegible] |
| 67 | 0.00 | UP_SEQ_FEATURE | TRANSMEM:Helical; Name=4 | 7 | 312 | 1084 | 20669 | 2.18 | [illegible] |
| 67 | 0.00 | UP_SEQ_FEATURE | TRANSMEM:Helical; Name=1 | 7 | 312 | 1099 | 20669 | 2.18 | [illegible] |
| 67 | 0.00 | UP_SEQ_FEATURE | TRANSMEM:Helical; Name=2 | 7 | 312 | 1106 | 20669 | 2.18 | [illegible] |
| 67 | 0.00 | INTERPRO | GPCR_Rhodpsn | 3 | 314 | 739 | 20827 | 0.93 | [illegible] |
| 67 | 0.00 | UP_KW_MOLECULAR_FUNCTION | G-protein coupled receptor | 4 | 187 | 862 | 12059 | 1.25 | [illegible] |
| 67 | 0.00 | INTERPRO | GPCR_Rhodpsn_7TM | 3 | 314 | 758 | 20827 | 0.93 | [illegible] |
| 67 | 0.00 | GOTERM_MF_DIRECT | G protein-coupled receptor activity | 4 | 306 | 869 | 19272 | 1.25 | [illegible] |
